# Rat Leukocyte Population Dynamics Predicts a Window for Intervention in Aging

**DOI:** 10.1101/2022.01.12.476126

**Authors:** Hagai Yanai, Christopher Dunn, Bongsoo Park, Christopher Coletta, Ross A. McDevitt, Taylor McNeely, Michael Leone, Robert P. Wersto, Kathy A. Perdue, Isabel Beerman

**Author notes:** Corresponding authors: Isabel Beerman and Hagai Yanai, BRC, 251 Bayview Blvd, Suite 100/10C220, Baltimore, MD, 21224, USA. Equal contribution (Hagai Yanai, Christopher Dunn).

## Abstract

Age-associated changes in human hematopoiesis have been mostly recapitulated in mouse models; but not much has been explored in rats, a physiologically closer model to humans. To establish whether rat hematopoiesis closely mirrors humans’, we examined the peripheral blood of rats throughout their lifespan. Significant age-associated changes showed distinctive population shifts predictive of age. A divergence between predicted versus chronological age changes was indicative of fragility; thus, these data may be a valuable tool to identify underlying diseases or as a surrogate predictor for intervention efficacy. Notably, several blood parameters and DNA methylation alterations defined specific leverage points during aging, supporting non-linear aging effects and highlighting a roadmap for interventions at these junctures. Overall, we present a simple set of rat blood metrics that can provide a window into their health and inform the implementation of interventions in a model system physiologically relevant for humans.

## Introduction

Among all mammalian tissues, blood is perhaps most easily collected in fairly large quantities for various advanced analyses, with modest discomfort to the donor. This has made blood the prevailing tissue for a wide variety of applications such as clinical diagnosis, immune surveillance, and molecular investigations, including DNA methylation, that require a sizable amount mass of genetic material ^1, 2^.

To date, peripheral blood aging has been largely characterized in humans ^3–5^ and mice ^6^. Studies in the mouse model have provided important insights as this model system allows investigation of the impact of a variety of genetic interventions and hematopoietic stem cell (HSC) transplant assays. Many murine aging phenotypes accurately reflect reports from human studies including a decline in immunogenic response of lymphoid cells ^6^, decreased CD4/CD8 T-cell ratio ^7^, and an increase in myeloid cells at the expense of lymphoid cells ^8^. These phenotypes have been shown to be influenced, at least in part, by intrinsic HSC aging ^9^.

However, the mouse as a model of human hematopoiesis has drawbacks including the fact that the common laboratory strains do not reliably develop spontaneous blood cancers, which is a common aging phenotype in humans ^10^. The differences between the two species are further exemplified by the fact that mouse blood is dominated by lymphocytes whereas human blood is rich in neutrophils ^11^; which in the murine model may dampen the impact of the age-associated decrease in lymphoid potential.

An attractive alternative system is the rat model. Rat physiology is more analogous to humans ^12, 13^, yet still have relatively short lifespans ^14^ in which to study aging phenotypes. Further, the rat and human genomes share genes involved in immunity and hematopoiesis absent in mice ^13^. Importantly, rats, especially the Fischer 344 CDF (F344) strain, display age-related incidence of leukemia ^15^. All the above suggest the rat may be better suited to study the aging of the blood and immune systems. Despite these advantages, use of the rat model has been in decline as mice are smaller, generally cost less to maintain, and transgenic tools to manipulate the murine genomes have been robustly developed. As a result, the impact of aging on the blood compartment in rats is not well characterized.

We sought to describe changes in the composition of the peripheral blood during aging in the F344 male rat using flow cytometry. In addition, we found that CD25^+^ T cell frequency is a novel marker for predicting aging and rats have defined age-associated inflection points linked to altered methylation profiles and cell composition. These results indicate that age-associated blood phenotypes in rats provide relevant insight into human blood aging and point to a critical time period of hematopoiesis which may best be targeted for anti-aging interventions

## Results

### Aging patterns in rat peripheral blood leukocyte populations

We analyzed the peripheral blood composition of 146 male F344 rats ranging in age from 3 to 27 months, using a panel of seven monoclonal antibodies to evaluate major leukocyte populations. As cell-specific gating strategies for rat blood populations are less defined than in humans or mice, we initially evaluated the flow cytometry results in an unbiased manner using t-distributed stochastic neighbor embedding (t-SNE) and Self-Organizing Maps analyses. Cells generally clustered in a manner that allowed for identification of defined populations (Fig. 1a and Supplementary Fig. 1) and clustering analysis helped inform decisions for gating strategies we devised for cell-type classification (Supplementary Fig. 2). Age-related changes were also profiled from single-cell analyses and revealed an accumulation of cells in clusters corresponding to myeloid lineage cells and a reduction of cells in clusters expressing markers associated with B and CD8^+^ T cells (Fig. 1b). Globally CD4^+^ T cells decreased with age, but we identified 2 distinct CD4^+^ populations with different trajectories during aging. The CD4^+^ T cell cluster expressing high levels of CD25 did not show an age-associated decline in cell number and instead increased in frequency over time. t-SNE defined clusters 5 and 9 (brown and cyan, Fig. 1a) also show age-associated increases in frequency (same top left location as Fig 1a, but blue in Fig. 1b); however, these clusters could not be fully identified from this combination of antibodies (Supplementary Fig. 2). Cluster 5, due to a high expression of CD11b/c, is most likely a type of myeloid cell, whereas cluster 9 most likely consists of debris (Supplementary Fig. 1) as it stained equally positive for all antibodies. Most identified cell populations that were categorized encompassed 2 or more clusters, indicating potential subpopulations we were unable to define with the panel of antibodies in this study.

**Figure 1.**
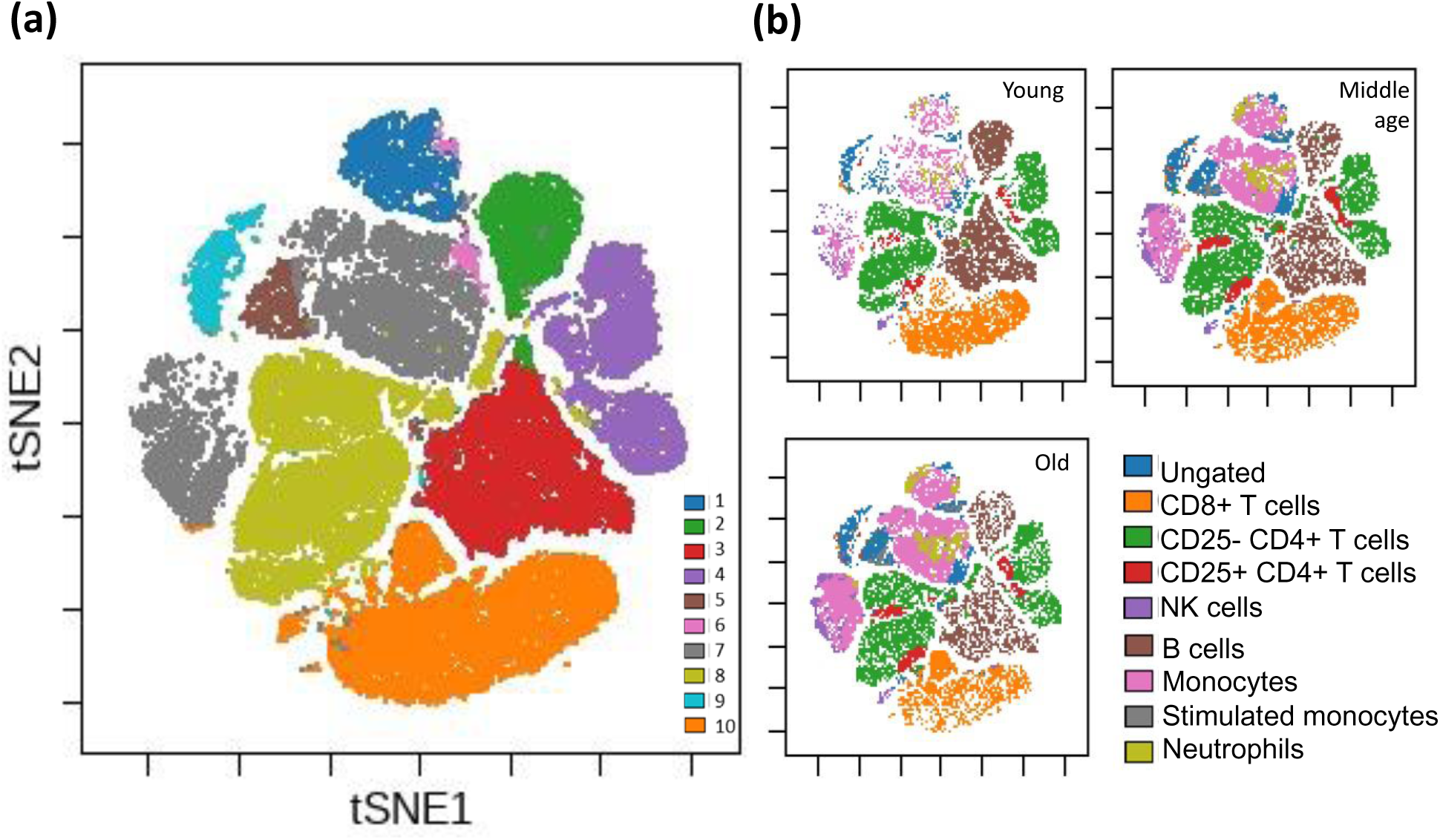
Unsupervised clustering of all F344 rat peripheral blood cells. Clustering of rat peripheral blood using a self-organizing map algorithm. For detailed clustering also see Supplementary Fig. 1. **(a)** Overall clustering of all leukocytes from 146 rats overlaid on a t-SNE map; each cluster is denoted as a different color (see legend). **(b)** Specific blood cell types illustrated on the same t-SNE map for three age groups (n=10,000 cells from each age group; total 30,000 cells). For visualization purposes, rats were divided into age groups: Young (3-4 mo), Middle-aged (15-23 mo) and Old (24+ mo).

We next sought to determine if age-specific blood parameter characteristics could be defined using frequencies of the distinct cell populations identified (Supplementary Fig. 2). Using unsupervised clustering of all rats based on leukocyte population frequencies, animals clustered into three discrete groups (Fig. 2), indicating the overall “fingerprint” of blood composition is age dependent. The cluster classified as “young” is characterized by a high proportion of lymphocytes in general, and T cells in particular, but with a low frequency of CD25^+^ CD4^+^ T cells.

**Figure 2.**
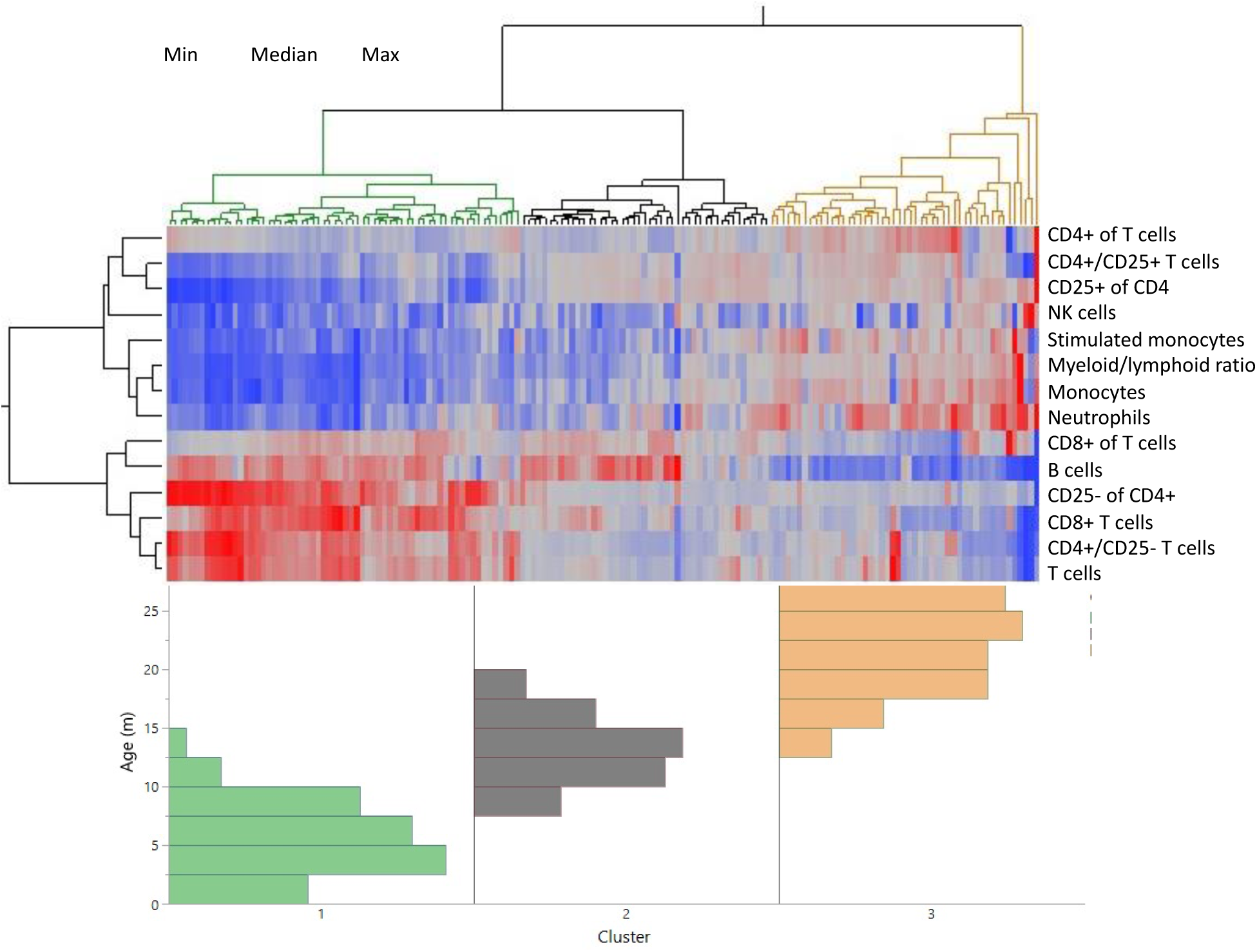
Distinct age clustering of rats based on white blood cell frequencies. Hierarchical agglomerative clustering with distance matrix by Ward’s method. The top lines represent the distance matrix for the individual rats and the three major branches are annotated by green, dark gray, and orange. The heatmap represents the normalized frequency of each population as indicated by the legend. The lines on the left indicate the distance matrix of the frequencies. The bottom histogram represents the age distribution for each of the major clusters in the same colors as above.

In contrast, the “old” cluster, on the far right, is characterized by a low overall lymphocyte frequency, and elevated proportions of both myeloid and CD25^+^ CD4^+^ T cells (Fig. 2). This increase in CD25^+^ CD4^+^ T cells is similar to what is seen in human peripheral blood aging ^16^, while in the murine model CD25^+^ CD4^+^ T cells only accumulate in secondary lymphoid organs (spleen and lymph nodes) but not in the peripheral blood ^17^. This analysis also defines a specific “mid-age” profile that is distinct from both young and old phenotypes, but clusters more closely to the young rat profile than the old.

### Peripheral blood parameters predict age and pathology in rats

To assess the blood cell types most predictive of aging, we fit all leukocyte frequencies in a standard least square model to predict aging in animals with no detected pathology and then eliminated the least significant parameters in a backstep fashion (Fig. 3a). The health status of all rats was recorded throughout the study, and necropsy was performed on all animals, less than a month post blood collection (for full health/pathology report see Supplementary Data 1). This enabled testing of how the health status of each rat fit into the model. One caveat was animals with overt sickness were excluded and euthanized per vet recommendation, so the health analysis captures mostly underlying illnesses only discernable during necropsy. The resulting blood parameter model predicted age with an approximate 2-month margin of error (as indicated by the root mean square error value). The single most predictive parameter of rat age was the proportion of CD25 expression in CD4^+^ T cells (Fig. 3a, right panel). Animals with histological indication of disease, including indications of large granular lymphocytic leukemia (LGLL) in spleen or liver, described in Finesso et al. ^18^, are generally predicted by our model as more variable than their actual age (i.e., have stronger residual age values) (Fig. 3b). When constructing an age prediction model only for ill animals, we found the myeloid/lymphoid ratio was the strongest contributor to age prediction (Fig. 3d). Additionally, we constructed a nominal logistic regression model to predict illness and found that myeloid/lymphoid ratio is the strongest contributor, even more so than age (Supplementary Fig. 3). While the myeloid/lymphoid ratio increases with aging, this phenomenon is more pronounced in sick animals. However, this ratio alone is not sufficient to predict illness (Fig. 3d, left), and must be combined with the other blood parameters that are characteristic of age and how variable the predicted age is from the chronologic age (Fig. 3b, c). We also observed that myeloid/lymphoid ratio increases concurrently with severity of LGLL ^18^ (Fig. 3d, right).

**Figure 3.**
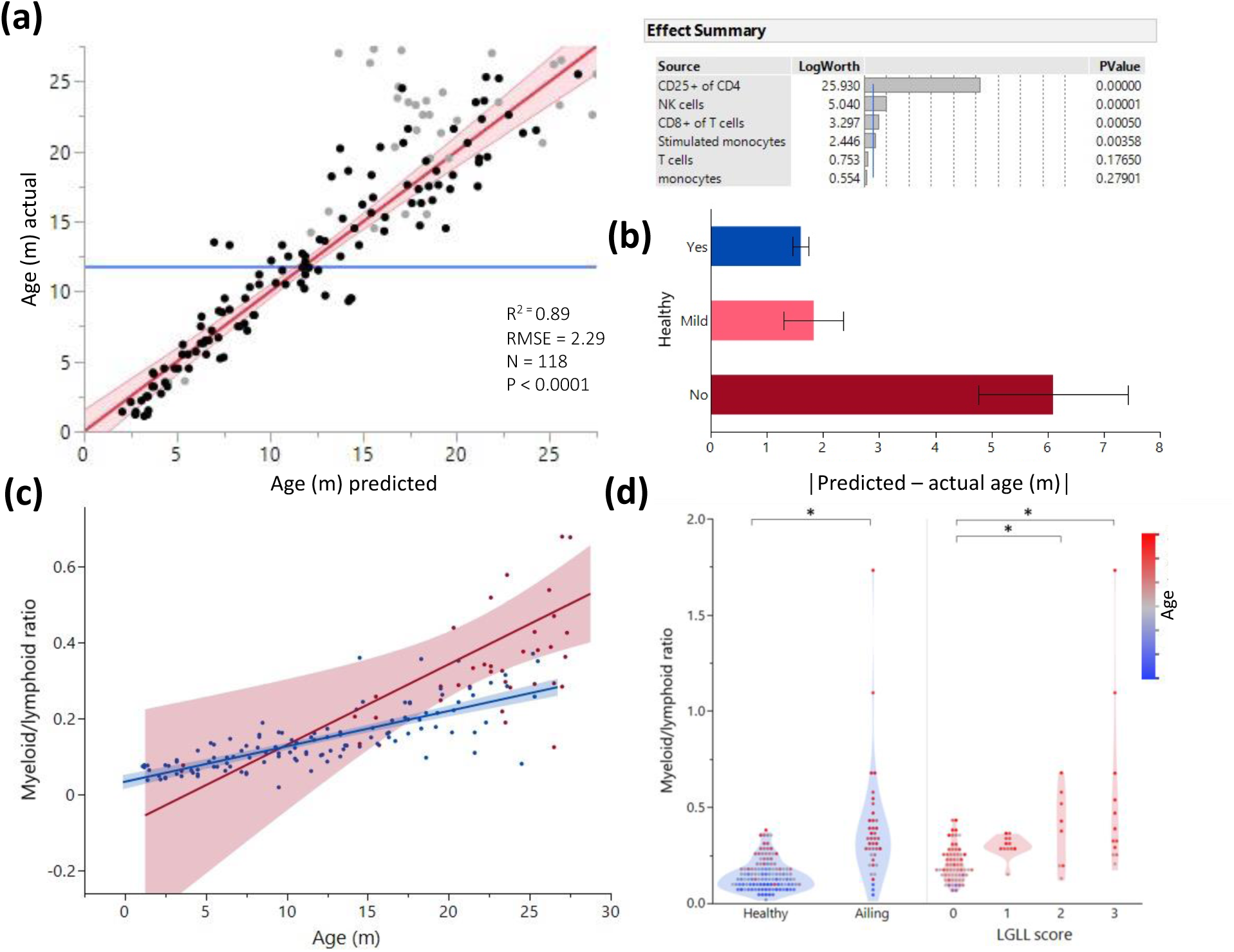
Age and illness prediction by blood leukocyte composition. All blood cell population frequencies were used to create a standard least square model of age prediction. Variables that did not contribute to the model were removed in a backstep fashion. **(a)** Predicted age vs. actual age, the red line indicating the mean and fit. Animals excluded due to illness are depicted in light gray. The effect summary for the variables used is depicted in the right panel (Effect Summary) with the LogWorth cutoff depicted by a blue line. **(b)** Residual age (i.e., the difference between actual age and predicted age) plotted as absolute mean values against assessment of health as determined by necropsy. Yes = no health issues; Mild = mild health problems (e.g., minor foot lesions); No = Clear health issues found during necropsy (Supplementary Data 1). **(c)** The correlation between myeloid/lymphoid ratio and age in healthy (blue) and ill (red) animals. **(d)** Myeloid/lymphoid ratio in the peripheral blood plotted against health status (left panel) and large granular lymphocytic leukemia (LGLL) pathology score as detected in the liver and spleen. Each dot represents a single rat. Dot color denotes age.

As blood samples from the 159 rats were analyzed post-fixation (see Methods), we were interested to determine whether the model generated is also applicable for freshly isolated blood. We thus generated data for both fresh and fixed rat blood, taken from the same donors. We found only small differences (<10%) in population frequencies associated with fixation, with changes in the T and B cell population frequencies post fixation showing the biggest variation (Supplementary Fig. 4a). Despite these differences, applying the age-prediction model, generated from fixed blood samples, on the blood profiles generated on un-fixed samples of varying ages (Supplementary Fig. 4b) demonstrated that the existing model (from fixed blood) could be used in a similar fashion to predict age with only minor mathematical adjustments.

### Regression analysis reveals potential aging intervention points

This study involved a minimum of 5 rats from each month of age starting from 3 months to 27 months (Supplementary Data 2) which enabled us to cross-sectionally examine the dynamics of blood changes during aging. Overall, the changes of leukocyte frequencies with age-matched rats, reflected described changes in aging human blood, including a reduction in B and T cells, and an increase in monocytes, neutrophils, and CD25^+^ CD4^+^ T cells (Supplementary Fig. 5). To better understand the dynamics of these blood composition shifts, we performed a multiple regression analysis for each cell population and identified time points in which there was a major trajectory shift (Fig. 4a and Supplementary Fig. 5b). Most of these shifts appear to converge at approximately 15 months of age. A strong age-associated increase in the overall variance of the different parameters was also observed, which was especially pronounced at close to 24 months of age (Fig. 4b). Interestingly, the first hematopoietic inflections appeared prior to the decline in survival of the classic Gompertz curve for this rat strain ^14^.

**Figure 4.**
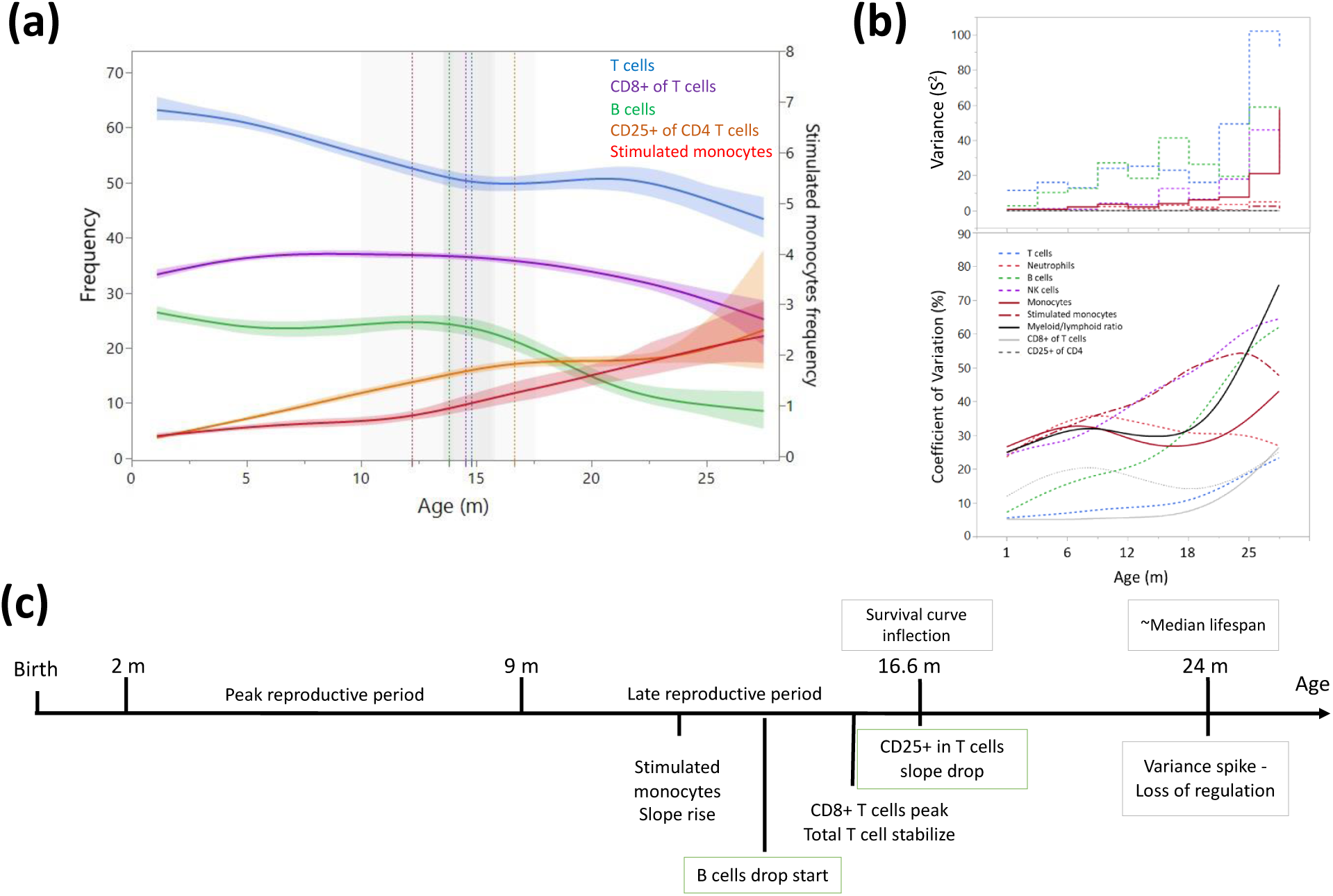
Dynamics of leukocyte frequencies during aging. **(a)** Aging trends of variables with inflection points depicted by a cubic spline regression (λ=0.521). Shaded colored areas indicate the fit. Inflection/breakpoints are depicted as dotted vertical lines with SE in shaded grey. **(b)** Variance (S2) and coefficient of variation (CV) for each parameter as a function of age (in a sliding window of 3 months). **(c)** Summary of leukocyte critical aging points (color coded) in relation to reproductive capacity (orange) and survival curve points (grey) ^14^.

### Peripheral rat blood DNA methylation changes with age

DNA methylation data was generated from the peripheral blood of these rats and was previously analyzed for defining a rat aging “methylation clock” ^2^. We analyzed the epigenic landscapes for hematopoietic-specific changes in methylation associated with aging using the Bismark pipeline. Global, age-dependent hypomethylation (Fig. 5a) occurred, located mostly in intergenic and intronic regions. Promoter regions had a more balanced ratio of sites that either gained or lost methylation (Fig. 5b). Interestingly, overall decreases in methylation were more pronounced in the transition between middle and old age compared to the global methylation changes seen in the shift from young to middle age (Fig. 5a). To gain insight into the difference between the early- and late-aging changes, we performed differentially methylated region (DMR) analysis on locations near transcription start sites (TSS). Most regions with significant DNA methylation changes were progressively altered during aging, which is slightly incongruous with the phase shifts seen in blood composition. However, we also found non-linear changes in methylation using a differential methylation analysis of 3 major age groups (Fig. 5). We identified 140 DMRs unique to the young vs. middle-age comparison and 94 DMRs unique to the middle-aged vs old (Fig. 5c). 456 DMRs appeared as significantly differential only between the young and old animals, which we attribute to a slow progressive change that only passed the cutoff of significance and fold change in the young vs old comparison (Fig 5c). Enrichment analysis of differentially methylated regions showed age-associated hyper-methylation for several processes and included some overlaps between the early (Y-M) and late (M-O) aging phenotypes, such as regulation of T cells and differentiation (Fig. 5d). However, some enrichments were unique to the middle age transition (regulation of apoptosis) and the late age (Notch signaling pathway), again indicating a difference between the early and late aging phenotypes. Of note, while hypomethylated DMRs between young and middle age were identified for several processes (Fig. 5d, right panel), hypomethylated DMRs in the transition from middle-aged to old were not enriched for any specific process (Supplementary Data 3); perhaps suggesting that these changes are more reflective of drift or random alterations and not necessarily concerted DNA methylation changes.

**Figure 5.**
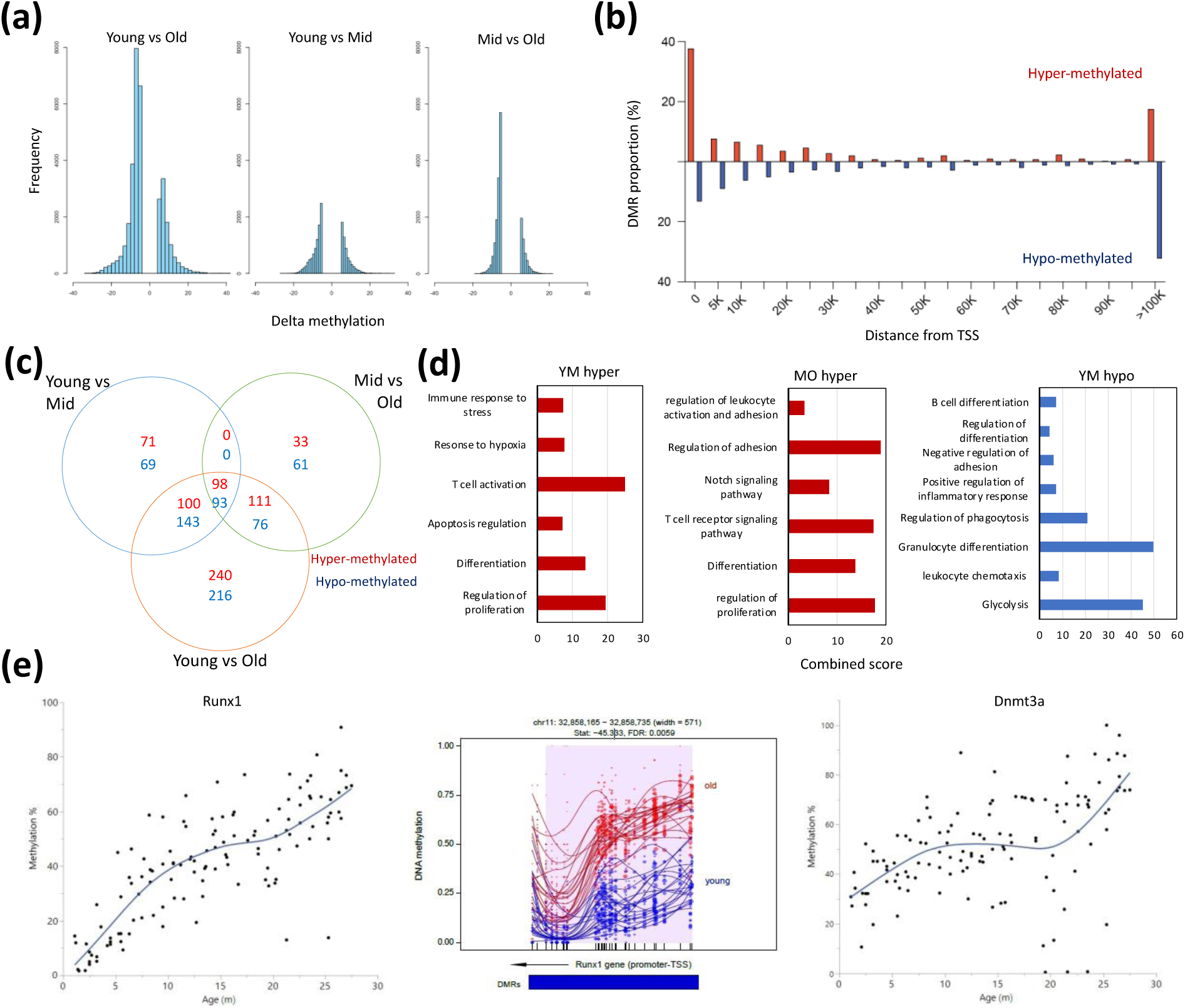
Differentially methylated regions with rat aging. **(a)** Global differential methylation histograms per age group comparison. A cut-off of 5% methylation difference was used. **(b)** Proportion of hyper- and hypo- methylated regions (DMR) compared to distance from the closest transcription start site (TSS). **(c)** Venn diagram for the number of differentially methylated regions within 20k bp distance of the nearest TSS based on age group comparisons. A cut-off of FDR<0.05, >5% CpG mean change, and at least 3 CpG differential DMR site per block was used. Hyper- and hypo-methylated changes are indicated by the noted colors. **(d)** Summary of gene set enrichment analysis for DMRs in proximity of TSSs (<20k bp). Analysis was performed for 3 age groups: Young (3-8 mo), Middle-aged (13-17 mo), and Old (22-27 mo). Combined score was calculated as -log10(FDR) x fold enrichment. YM - Young vs. Middle aged; MO - Middle aged vs. Old; YO - Young vs. Old. The list of hypomethylated DMRs for the MO comparison had statistically significant enrichments. For a full enrichment analysis see Table S4. E. Individual methylation levels of regions near the Runx1 and Dnmt3a genes. Left panel - Runx1; Middle panel - Methylation map in the Runx1 TSS; Right panel - Dnmt3a.

To eliminate a potential bias of pre-determining age groups, we also analyzed DNA methylation at all transcription start sites to determine if changes occurred as a function of age using the Response Screening platform (see Experimental Procedures) (Supplementary Fig. 6). The most significantly changed sites were all located near the TSS of RUNX family transcription factor 1 (*Runx1*) and these sites all show strong hypermethylation with age (Fig. 5e, left panel). The hypermethylation of *Runx1* can be mapped immediately distal to the TSS (Fig. 5e, middle panel). The second most significantly changed DMR was located near the TSS of *Dnmt3a* (Fig. 5e, right panel). A principal component analysis (PCA) based only on the 2,311 DMRs that significantly changed as a function of age (Supplementary Fig. 6) demonstrates a marked increase in variation around the 2-year mark, analogous to the increased variation observed for population frequencies (Fig. 4b).

### Early and late life DNA methylation switches

As the blood population frequencies indicated a mid-life inflection, we explored whether DNA methylation patterns exhibited similar signature transitions. First, we performed both linear and logistic regressions on significant age-associated DMRs near TSSs (Supplementary Fig. 6 and Supplementary Fig. 7). We next selected the regression with the best fit (based on Akaike Information Criterion Shavlakadze, et al. ^19^). 62.9% of the sites changed linearly with age while 37.1% showed a better fit to a logistic regression (Fig. S7). The break point was calculated for sites that did not change linearly with age (Fig. 6, left) and a module enrichment analysis was performed for genes with switches before and after 15 months of age (Fig. 6, right). The methylation switch points defined converged at around 22 months of age and mildly at close to 12 months of age. Genes that had a breakpoint earlier than 15 months of age were enriched for cytokine production, regulation of differentiation and hematopoiesis, and include Apolipoprotein E (*Apoe*) and *Runx1*. Genes identified to have a switch later in life were enriched for myeloid differentiation, leukocyte activation, ion transport, and epithelial development. Similar to the changes in blood parameter composition, these DNA methylation results highlight differences come in waves, notably at around 15 months, rather than occurring in a linear fashion and suggest a key remodeling time-point of the hematopoietic system.

**Figure 6.**
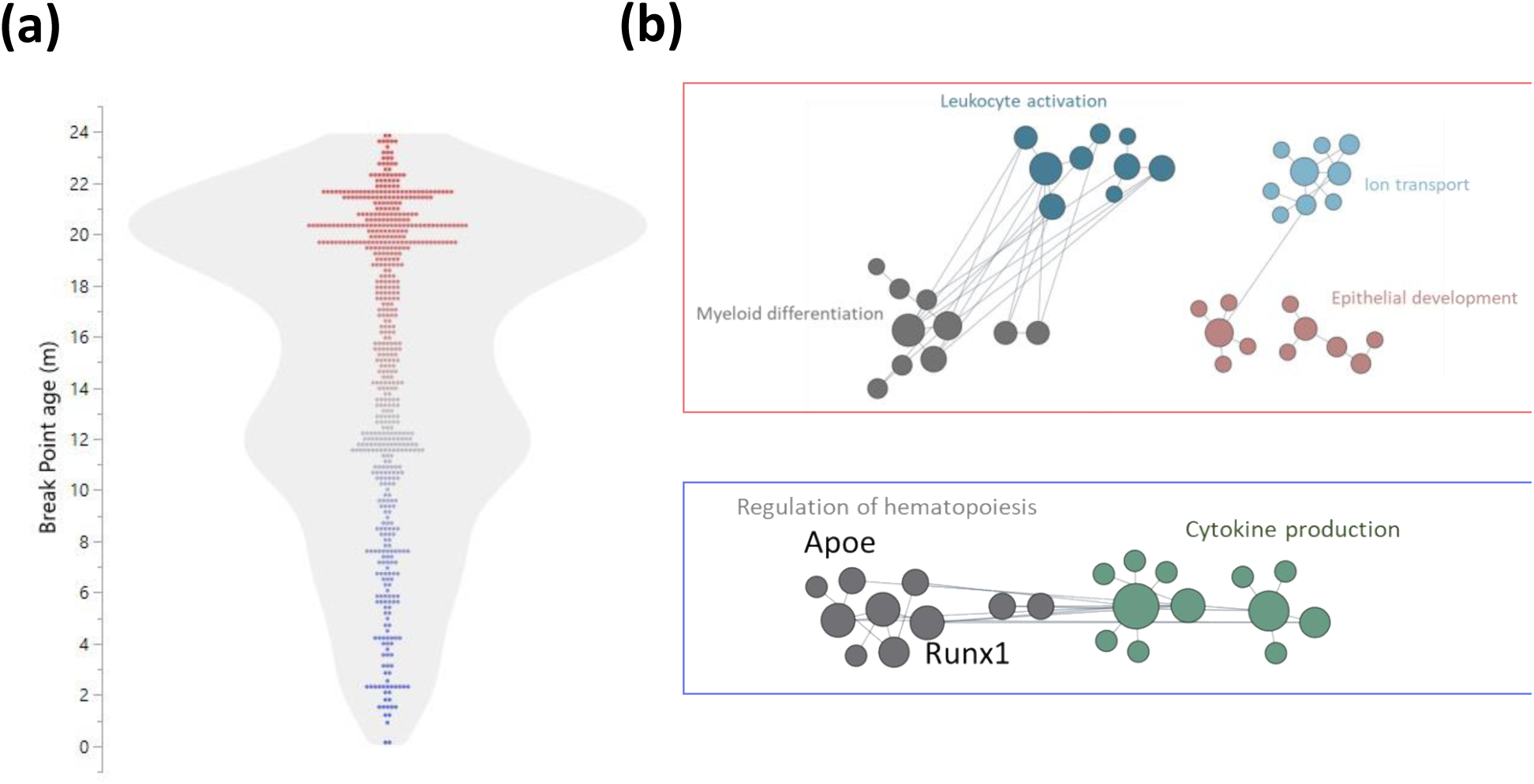
Early and late life methylation breakpoints. **(a)** Distribution of breakpoints by age. Each dot represents the breakpoint of a single gene TSS; color scales by age from young (blue) to old (red). Contour violin plot of the distribution is displayed in gray. **(b)** Functional module networks of the genes with breakpoints before (bottom - blue) and after (top - red) 15 months of age. Each dot represents a single gene with size corresponding to its’ connectivity in the network. The lines indicate interactions (edges). The networks were built for blood tissue, with human orthologs as input. The interaction network is built using the closest gene neighbors and then clustered based on enrichment in GO categories. Networks were generated using HumanBase (https://hb.flatironinstitute.org).

## Discussion

Although the rat peripheral blood compartment has been analyzed in regard to aging ^20^, newer panels of monoclonal antibodies offer improved insight into changes of the hematopoietic system. Here, a relatively simple blood profiling technique using 8 antibodies led us to construct a model that predicts both age and, potentially, pathology. The model can be used to predict the risk of illness as a function of the deviation from predicted age to actual age, and we propose such a model could also be used to determine if intervention experiments mitigate aging phenotypes. While our dataset and analyses were for a specific set of flow cytometric parameters generated on fixed cells, we found we were able to adapt these analyses to also predict the age of freshly isolated blood (using similar flow cytometry markers). Thus, with inclusion of proper controls to account for experimental design variation, this resource of aging rat blood parameters could be adapted for studies of rat aging or pathology without requiring euthanasia of the animals. We believe this would be especially useful for longitudinal or intervention studies given the ease of blood collection at multiple time points.

A strong predictor of age was the accumulation of CD25 on CD4^+^ T cells in the F344 rat model. These results support previous findings that report an age dependent accumulation of regulatory T cells (Tregs) ^17, 21, 22^. This stable increase in the F344 rat is more similar to that observed in humans ^16^ than in mice ^17^ and highlights the relevance of the rat model to study hematopoietic aging. Garg et al. ^23^ have previously suggested that this accumulation is driven by hypomethylation of forkhead box P3 (Foxp3) in CD25^+^CD4^+^ T cells, however in our dataset analyzing total blood cell methylation, we do not see differential methylation of Foxp3. This is likely because CD25^+^CD4^+^ T cells are a rare subpopulation of the total blood and DNA methylation changes in this population would not significantly affect the global blood methylation profiles.

We observed an age-dependent myeloid bias with rat aging, similar to mice ^9^ and humans^8^. The observed myeloid bias was a strong predictor of pathology in general, and of LGLL in particular. Importantly, strong myeloid bias was correlated with illness that was otherwise only noticeable during autopsy and undetectable during routine veterinary observation. A caveat to this observation is that myeloid-bias increases with age, even in healthy animals, meaning that the myeloid/lymphoid ratio can only be useful as a diagnostic parameter when plotted against the expected “normal” age-dependent increase. In this reference dataset, we found a significant association between severity of LGLL and increased myeloid cell numbers. The myeloid bias we report in rats has previously been documented in mice, however most mouse models do not develop age-associated blood diseases seen in humans. Thus, we find the correlation between increased myeloid cells and LGLL severity in rats to be of particular relevance for potential modeling of human blood aging phenotypes.

Our analysis of the age-related changes in leukocyte profiles indicates inflection points in the trajectories occurring at early middle-age (∼15 months) and at ∼24 months of age. The first inflection point is characterized by several shifts in leukocyte frequencies, including a stabilization of T cell decline, peak in CD8^+^ T cells, rapid drop of B cells, deceleration in the accumulation of CD25^+^ CD4^+^ T cells, and an acceleration in the increase of stimulated monocytes. The second shift point we observed is defined by a marked increase in variance between individual rats which alludes to a pan-hematopoietic loss of homeostasis. We hypothesize that interventions aiming at mitigating aging phenotypes would be significantly more effective if performed prior to these defined shift points. The inflection points in the blood are similar to the transition pattens (linear, early logistic, and mid-logistic) of gene expression on tissues isolated from aging Sprague- Dawley rats ^19^. Further supporting a claim for interventions predating the observed switch point are studies of caloric restriction (CR). It is quite clear that early onset CR has more benefits that late onset, but it is yet unclear what the ideal age of onset should be ^24^. In fact, late onset CR has even been shown to be potentially detrimental to cognitive function when applied too late ^25^. However, Chen et al ^26^ reported a beneficial effect of late onset CR to muscle mass and metabolism in Sprague-Dawley rats, indicating that the ideal points of intervention could vary greatly between different systems. In fact, interventions that aim to prevent or delay aging could be fundamentally different than those that aim to reverse it, even to the extent where harm could be caused when one strategy is applied instead of the other ^27^. It is therefore critical to understand which systems display aging switch points, and when, in order to design the appropriate intervention regimen. This is especially true when translating experimental paradigms to human interventions due to the great heterogeneity between the aging rates of both individuals and different systems withing the individual ^28^.

The combined results of changes in population frequencies and the DNA methylome indicate that hematopoietic aging of rats occurs in phases as opposed to a continuous process. However, with the current data, we cannot ascertain the drivers underlying the shift between these aging phases, whether they are triggered by a specific event, or simply manifest when a critical mass of small changes reaches a threshold. Whichever the case, investigating these relatively early time points in which blood profiles begin to irreversibly shift, indicating critical trajectory alterations, is of great interest.

The age-related DNA methylation changes in all leukocytes highlights the potential involvement of epigenetic changes that have occurred in hematopoietic stem cells and/or early progenitors, and are transmitted to all the differentiated progeny, as indicated by the hypermethylation of *Runx1* and the gene enrichment analysis. In addition, the lack of enrichment for hypomethylated transcription start sites in the transition from middle age to old indicates that age-related gains of methylation may be a directed process, whereas loss of methylation later in life is more non-specific and perhaps attributed to drift. If so, mitigating the seemingly random loss of methylation would be an interesting potential aging intervention.

In summary, we present this work as a resource for investigators studying aging in the rat model, which appears to be a robust system for modeling the aging human hematopoietic system. Our research emphasizes the importance of studying changes in homeostasis during aging as we present key trajectory shifts in blood populations that occur at relatively early age (15 months). Finally, our research model describes aging “juncture points.” Given these specific breakpoints in composition of the hematopoietic system, we posit that strategic aging interventions would likely be more robust and beneficial if performed earlier in life to mitigate or stall the major changes that occur under homeostatic conditions. It would be exciting to further explore if an illness prediction model based off of these defined blood parameters could be extrapolated for a broad range of diseases, and ultimately if similar age-associated shifts occur in other organs/tissue systems, animal models, and specifically in humans.

## Methods

### Animals

All experimental procedures were conducted in accordance with the *Guide for the Care and Use of Laboratory Animals* and approved by the NIA Animal Care and Use Committee. Male Fischer 344 CDF (F344) rats were obtained from the NIA Aged Rodent Colony housed at the Charles River Laboratories (Frederick, MD). Animals were housed with Nylabone supplementation and ad libitum access to food (Envigo 2018SX) and water. Rats younger than 3 months were housed in groups of three; all other rats were single housed. All rats were maintained on a 12/12 lighting schedule, with all procedures carried out during the light cycle. Rats were habituated to the facility for at least 3 days before sample collection. The same rats were analyzed in previous studies ^2, 18^.

### Flow cytometry and sample collection

500 µl of whole blood was collected via retro-orbital bleedings for DNA and FACS analysis. Blood for DNA was collected in heparinized tubes, spun, and the plasma removed; buffy coat and red blood cells were frozen at −80°C until DNA extraction. Blood for FACS analysis was collected in EDTA-treated tubes, chilled on ice, and 100 µL was stained and then processed using a Beckman Coulter TQ-prep (fixation step) and the Beckman Coulter immunoprep reagent system. For the fresh sample analysis, blood was drawn as above, ACK treated and immediately stained on ice. Antibodies used for fluorescence analysis are as follows: FITC-conjugated anti-rat CD3 (clone 1F4, Cat#201403), PE-conjugated anti-rat CD25 (clone OX-39, Cat#202105), PerCP-conjugated anti-rat CD8a (clone OX-8, Cat#201712), PE-Cy7 conjugated anti-rat CD11b/c (clone OX-42, Cat#201818), APC-Cy7 conjugated anti-rat CD4 (clone W3/25, Cat#201518) from Biolegend (San Diego, USA), and AF647-conjugated anti-rat RT1B (clone OX-6, Cat#562223), BV421-conjugated anti-rat CD45RA (clone OX-33, Cat#740043), and BV605-conjugated anti-rat CD45 (clone OX-1, Cat#740515) from BD Biosciences (Franklin Lakes, USA). Immunophenotyping data was acquired on a BD FACSCanto II and analyzed using FlowJo (https://www.flowjo.com/).

### DNA methylation analysis

Samples were treated and sequenced for DNA methylation as described in Levine et al.^2^. The raw sequencing datasets are available from GEO (GSE161141). To identify differentially methylated loci: we trimmed reads of adapter dimers using Trim Galore (0.4.3) and quality trimmed with a minimum quality score above >25 (--rrbs -q 25). The attached adapter dimers were trimmed using cutadapter. Firstly, bisulfite-converted index (GA and CT conversion) was generated using F344 rat genome with Bismark build option and trimmed reads were aligned with bismark ^29^. Once we created aligned reads and corresponding locations, we used the bismark_methylation_extrator tool to summarize the level of methylation in CpG sites (bismark_methylation_extractor -p –comprehensive –no_overlap –bedGraph –counts –buffer_size 16G ($Aligned read bam file). Approximately a total 1-2 million sites per sample were predicted with DNA methylated sites (or unmethylated). Differentially methylated regions (DMR) and blocks of differentially methylated sites were identified with a minimum of 3 CpG sites per block and at least >5% Methylation difference (FDR <0.05) using MethylKit [PMID: 23034086^30^] and DMRseq ^31^. Functional annotation was performed using ClusterProfiler ^32^.

To generate RRBS DNA methylation block: RRBS datasets were processed in a uniform way and DNA methylation levels for each sample were extracted. Then, we generated 200 base-pair binned DNA methylation levels across the genome. Each DNA methylation block contains 1-20 CpG sites. We first calculated the average DNAm level per DNAm block for each rat age then used this average DNAm level for mean CpG imputation. We calculated the DNAm level using all methylated and unmethylated sites from the binned CpG sites when the minimum coverage was more than 5. The F344 build of the rat reference genome was used for bisulfite sequencing alignment, and the rn6 genome feature was used to extract genomic annotation information using the Homer package ^33^. Genes, exons, introns, and UTRs were taken from Homer annotation tools (annotatePeaks.pl DMR rn6). TSS-promoter sites were considered the identified DMRs close to TSS (less than 1kb). TSS-proximal sites were considered the identified DMRs close to TSS (less than 20kb) and TSS-distal sites were considered the DNAm block whose distance to TSS is above 20kb. For downstream analysis to find inflection points, we used the TSS-promoter DNAm block for searching inflection point analysis (e.g. Runx1, Dnmt3a).

### Statistical analysis and software used

Population frequencies were determined with FlowJo™ v10.8 Software (BD Life Sciences) ViSNE and Self-Organizing Maps (SOM) analyses were performed using Cytobank ^34^. All statistical analyses except those specifically detailed otherwise were performed using JMP (JMP®, Version 16. SAS Institute Inc., Cary, NC, 1989–2021). The predictive model of age based on leukocyte frequencies was constructed using the least square method with backstep elimination of insignificant variables. Analysis of pathology prediction was performed using the JMP Decision Tree Platform. Determination of regression formulas and breakpoints were performed in JMP according to the formulas indicated in Fig. S7.

## Acknowledgements

Many thanks to Dr. Rafael de Cabo, Dr. Luigi Ferrucci, Dr. Michel Bernier, and all members of TGB for invaluable support. We would like to thank all the members of the NIA Comparative Medicine Section for their consistent efforts and high standards of animal care. A special thank you to Daniel Ariad for invaluable advice. This research was supported entirely by the Intramural Research Program of the NIH, National institute on Aging.

## Author contributions

C.D., T.M., and M.L. conducted experiments; B.P. and C.C. participated in the bioinformatics analysis; R.M. was involved in study design and data analysis; R.W., and K.P. participated in study design; I.B. supervised the study and edited the manuscript. H.Y. conducted experiments, analyzed the data, and wrote the manuscript with input from all authors

## Data availability statement

The data that support the findings of this study are available in the supplementary material and Supplementary Tables of this article. The DNA methylation raw data is available via GEO Accession (GSE161141). Further information and requests for resources and reagents should be directed to and will be replied to by the corresponding authors.

## Supplementary Information

**Supplementary Figure 1.**
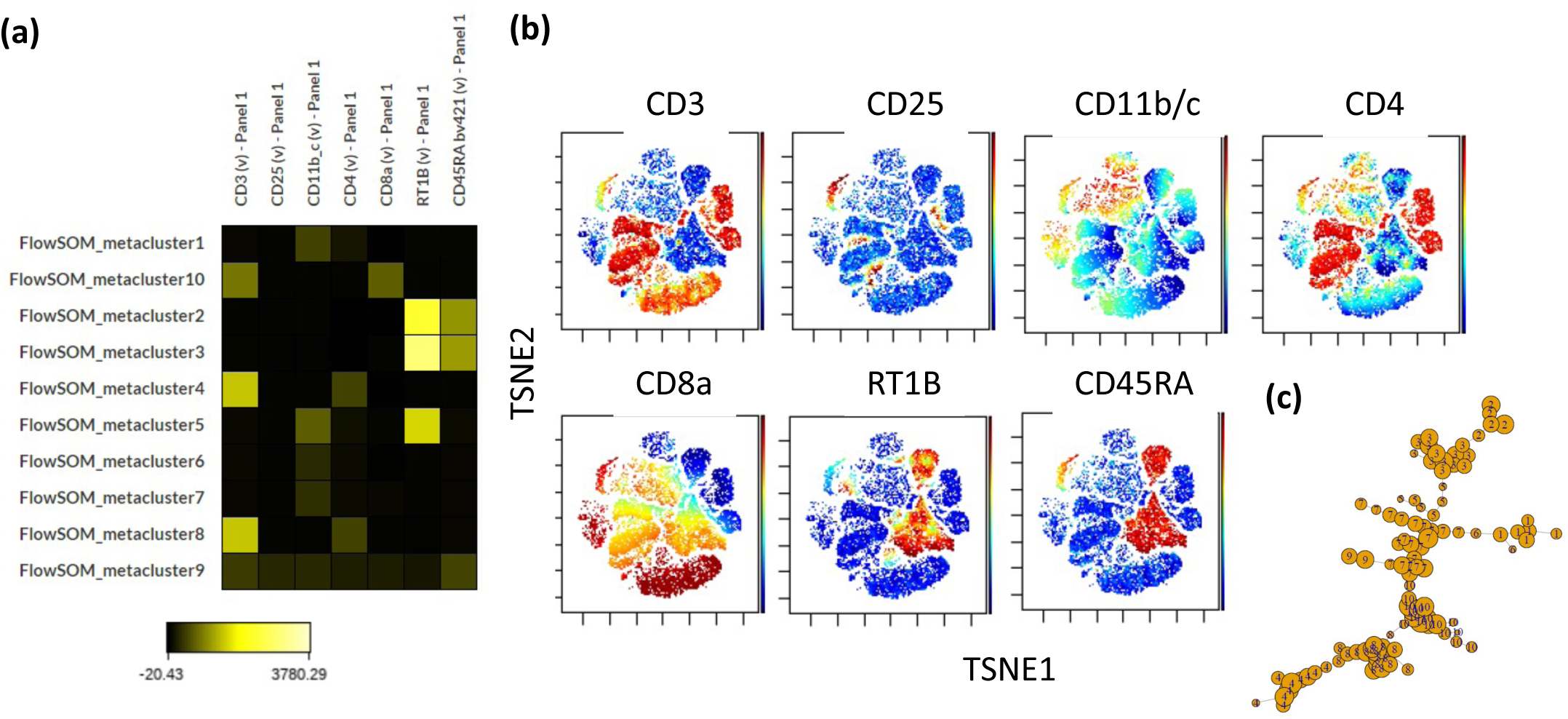
Self organizing map clustering of F344 rat leukocytes. 1.3 million random cells from all samples were chosen to perform a self organizing map analysis on the cytobank platform using default settings. **(a)** Heatmap of Ab fluorophore intensities per cluster. **(b)** Fluorophore intensities (blue = low intensity to red = high intensity) overlaid on a ViSNE plot generated on the same platform using default settings. **(c)** Divergence tree of clustering.

**Supplementary Figure 2.**
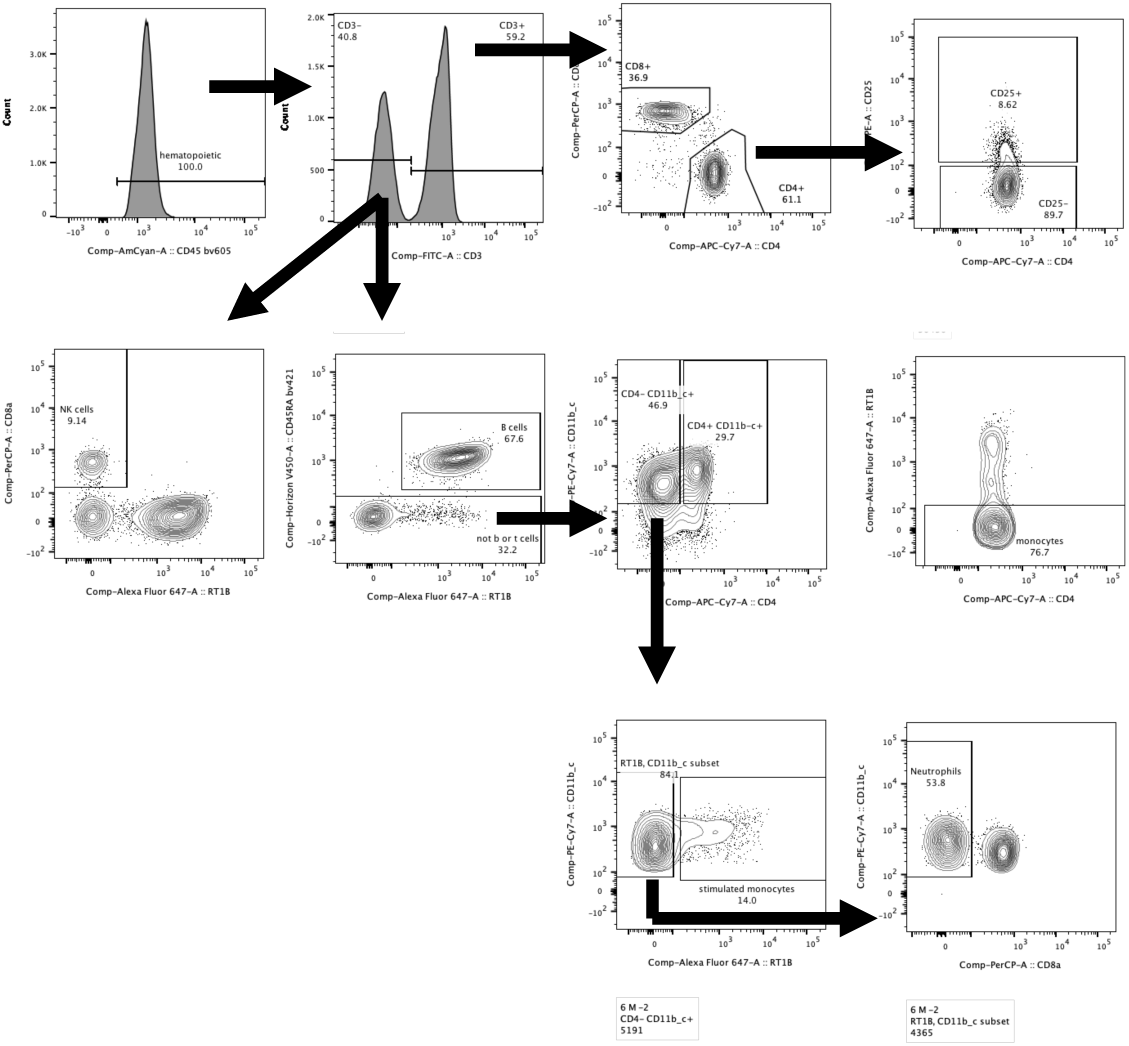
Gating strategy. The staining panel was uniform for all samples and consisted of the following anti-rat Abs: CD45-BV605, CD3-FITC, CD8a-PerCP, CD4-APC.Cy7, CD25-PE, RT1B-AF647, CD45RA-BV421, and CD11b/c-PE.Cy7.

**Supplementary Figure 3.**
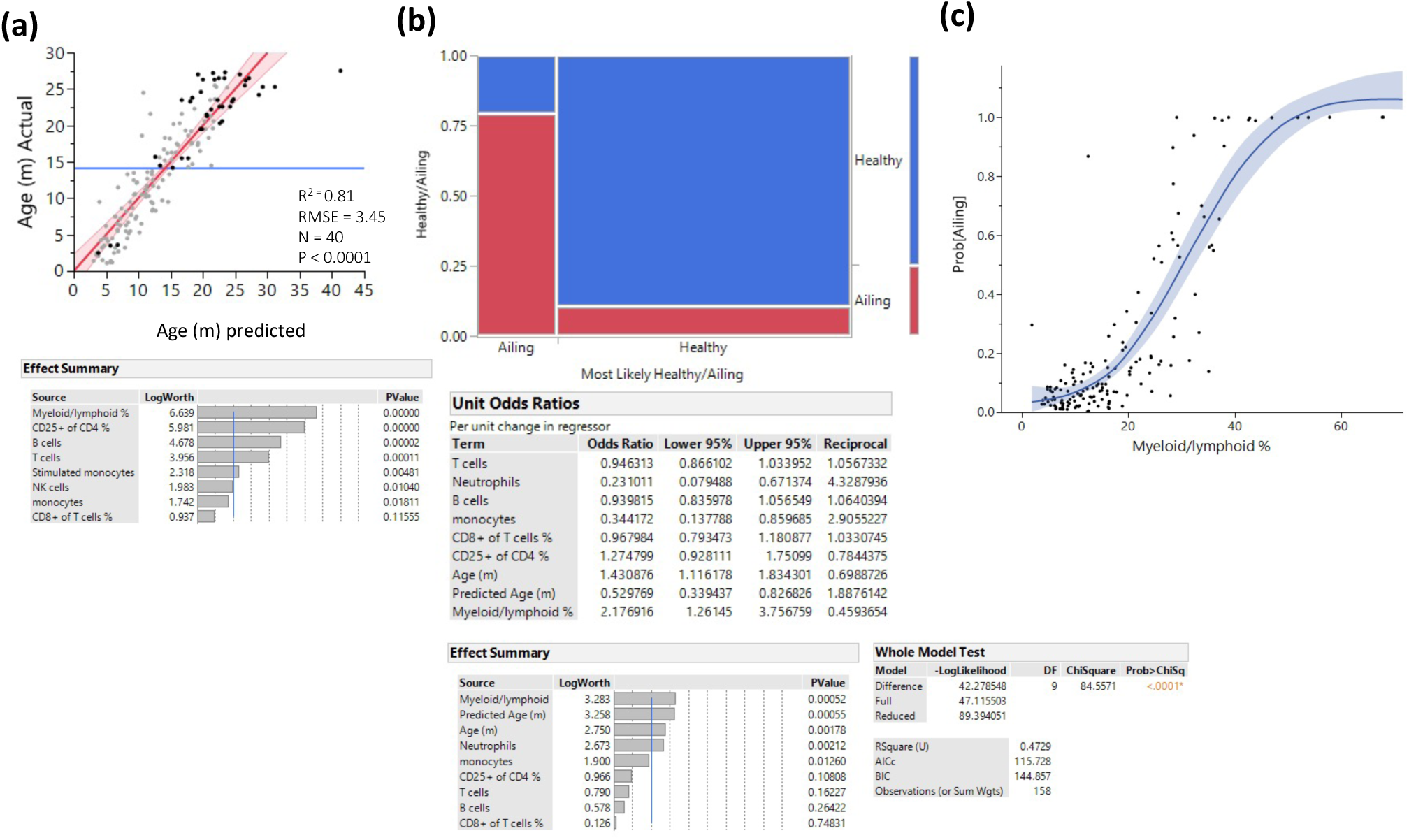
Prediction of pathology. Several methods were used to identify myeloid/lymphoid ratio as a strong predictor of pathology. **(a)** Prediction of age only among animals with any kind of pathology by least square method. Statistics are depicted and the variable contribution is depicted in the lower panel. **(b)** Visualization of a nominal regression model to predict pathology, the X axis is divided into animals that were predicted to be healthy/ailing and the proportions of actual health status are denoted in blue (healthy) and red (ill). The lower panels denote the nominal regression model statistics and parameter contributions and odd ratios. **(c)** Probability of being ill is plotted as a function of myeloid/lymphoid ratio (line denotes a spline regression with the colored area denoting the fit).

**Supplementary Figure 4.**
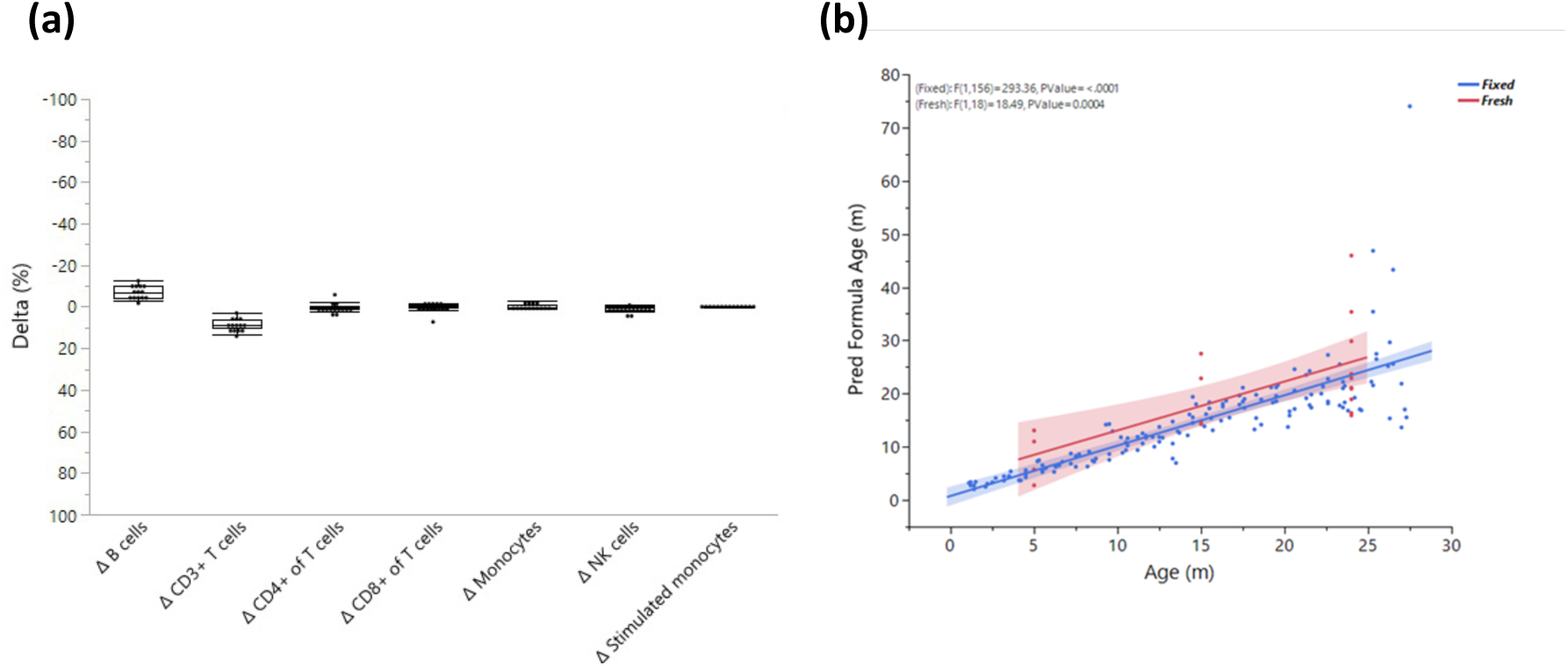
Fresh Vs fixed samples. **(a)** Blood from the same rats (10 months old) was divided and stained after fixation and on fresh live cells. The cells were analyzed by flow cytometry and the delta for each parameter was calculated as Fresh value – Fixed value. Each dot represents the delta for the given parameter in the same rat and the box indicates the mean and quantiles. n = 14. **(b)** Blood from 5 young (5 mo), 5 middle-aged (15 mo), and 11 old (24 mo) rats was collected and analyzed, and the results were fitted (red) to the model generated from fixed samples (blue).

**Supplementary Figure 5.**
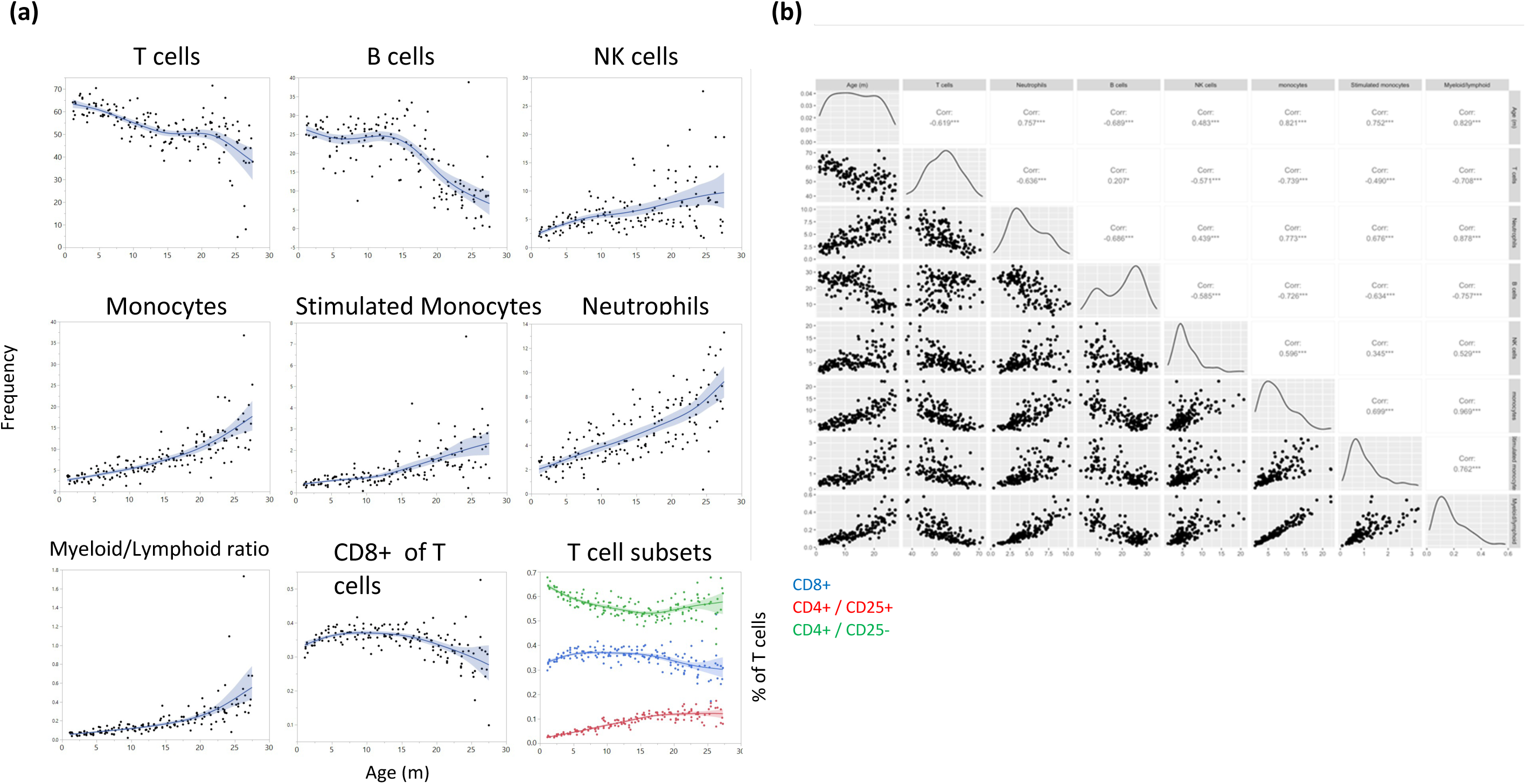
Major leukocyte population frequencies as a function of age. **(a)** Each dot represents a reading from a single rat and the cell type is indicated for each plot. The line represents a spline regression, and the colored area represents the fit. **(b)** Bivariate correlation matrix. The diagonal indicates the univariate distribution for the given variable

**Supplementary Figure 6.**
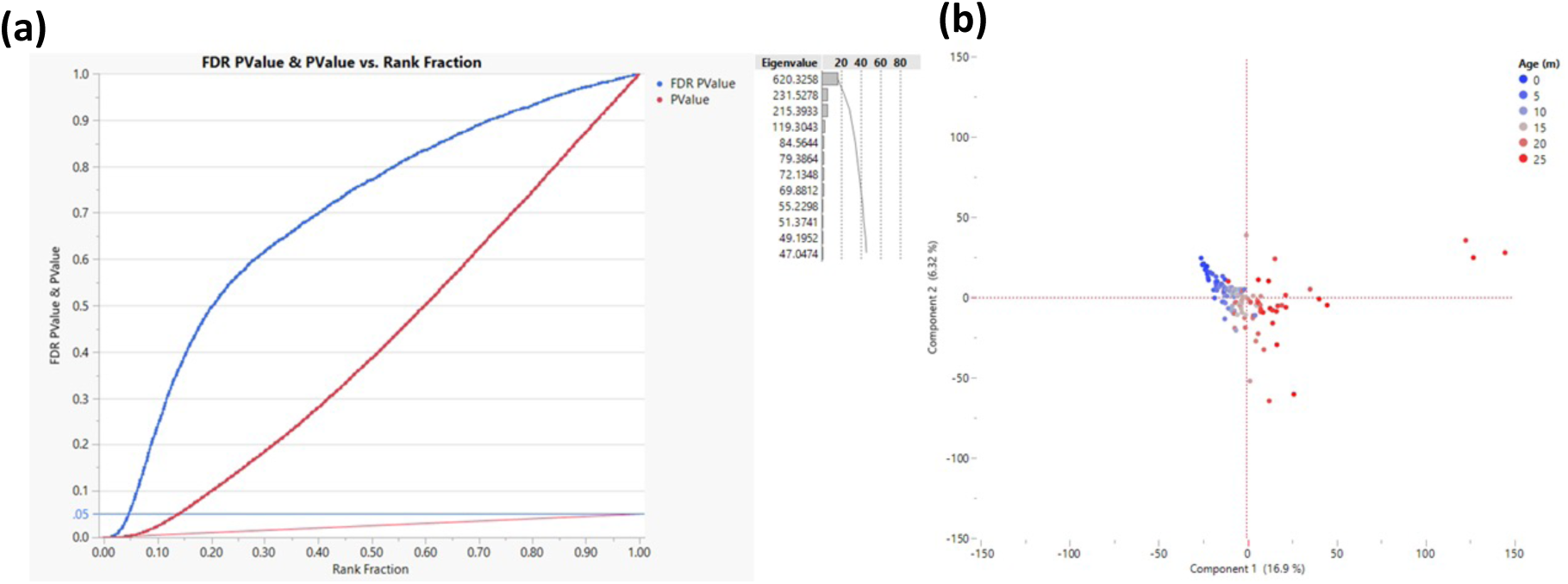
Response Screening to determine transcription start sites that change their methylation as a function of age. The Response Screening function (JMP - https://www.jmp.com) performs multiple correlation analyses and corrects for False Discovery Rate (FDR). The analysis was performed on 200 bp bins that are within 500 bp of TSS. **(a)** Correlation P values are depicted in red and the FDR in blue where each TSS is a single dot. The horizontal blue line indicates the cut-off (FDR p value < 0.05, measured in >50 rats, n = 2,311). Only sites that were successfully sequenced in >50 animals were included (n = 58,000). **(b)** PCA for all rats based on the genes that passed the cut-off; each dot represents a rat and the color represents age as indicated.

**Supplementary Figure 7.**
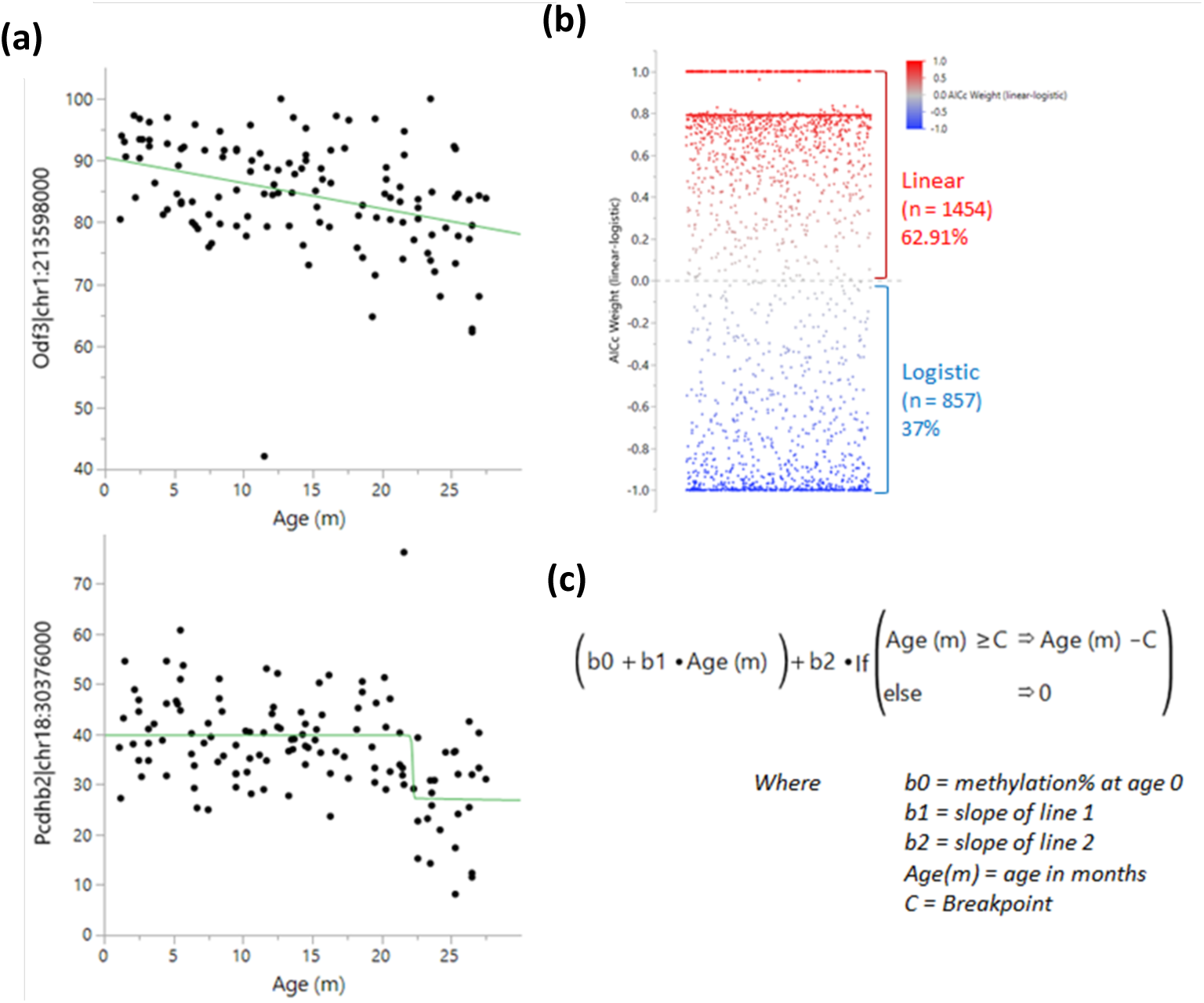
Determination of methylation changes “switch” points. A logistic 4P and linear regression were both tested for each bin near a TSS and the results were compared using the Bayesian Information Criterion (BIC) and Corrected Akaike Information Criterion (AICc). **(a)** Example of a linear (top) and a logistic (bottom) regression. **(b)** Comparison of weighted AICc delta (linear-logistic). Each dot represents a site. Values over 0 (red) indicate a linear regression fits better than a logistic one and vice versa for values under 0 (blue). **(c)** Breakpoint analysis formula used to determine the optimal switch point.

**Suppl Table 1.**
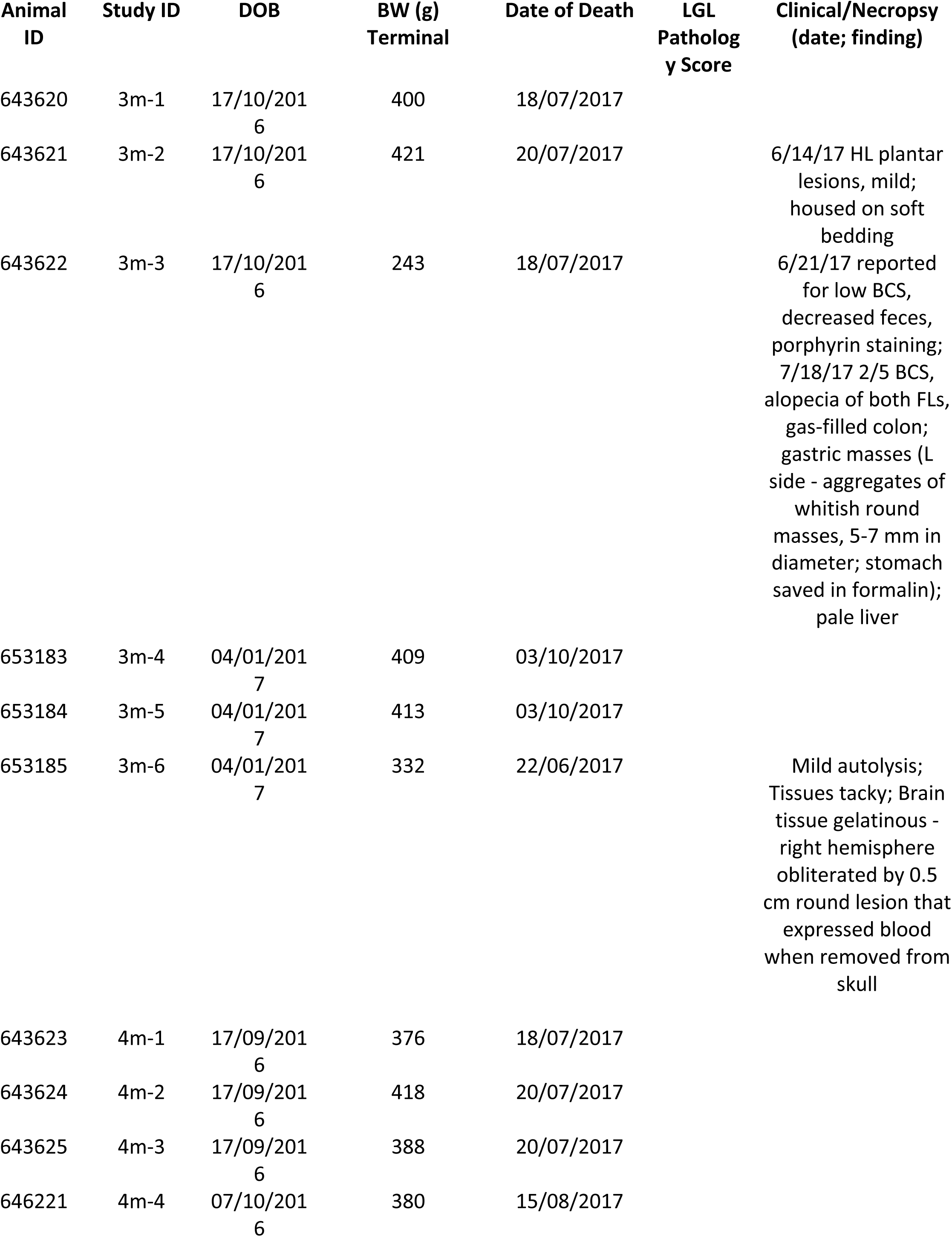
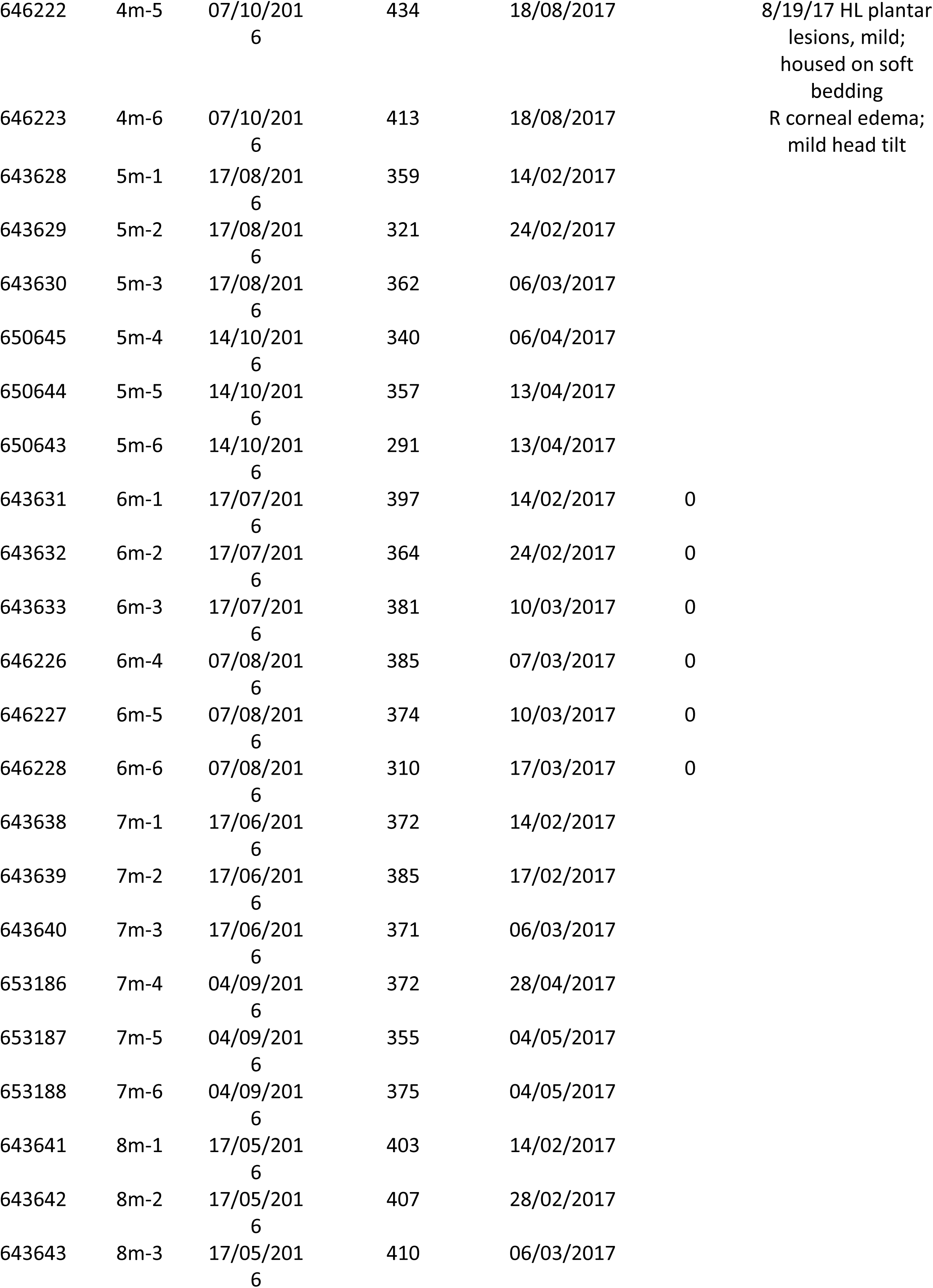
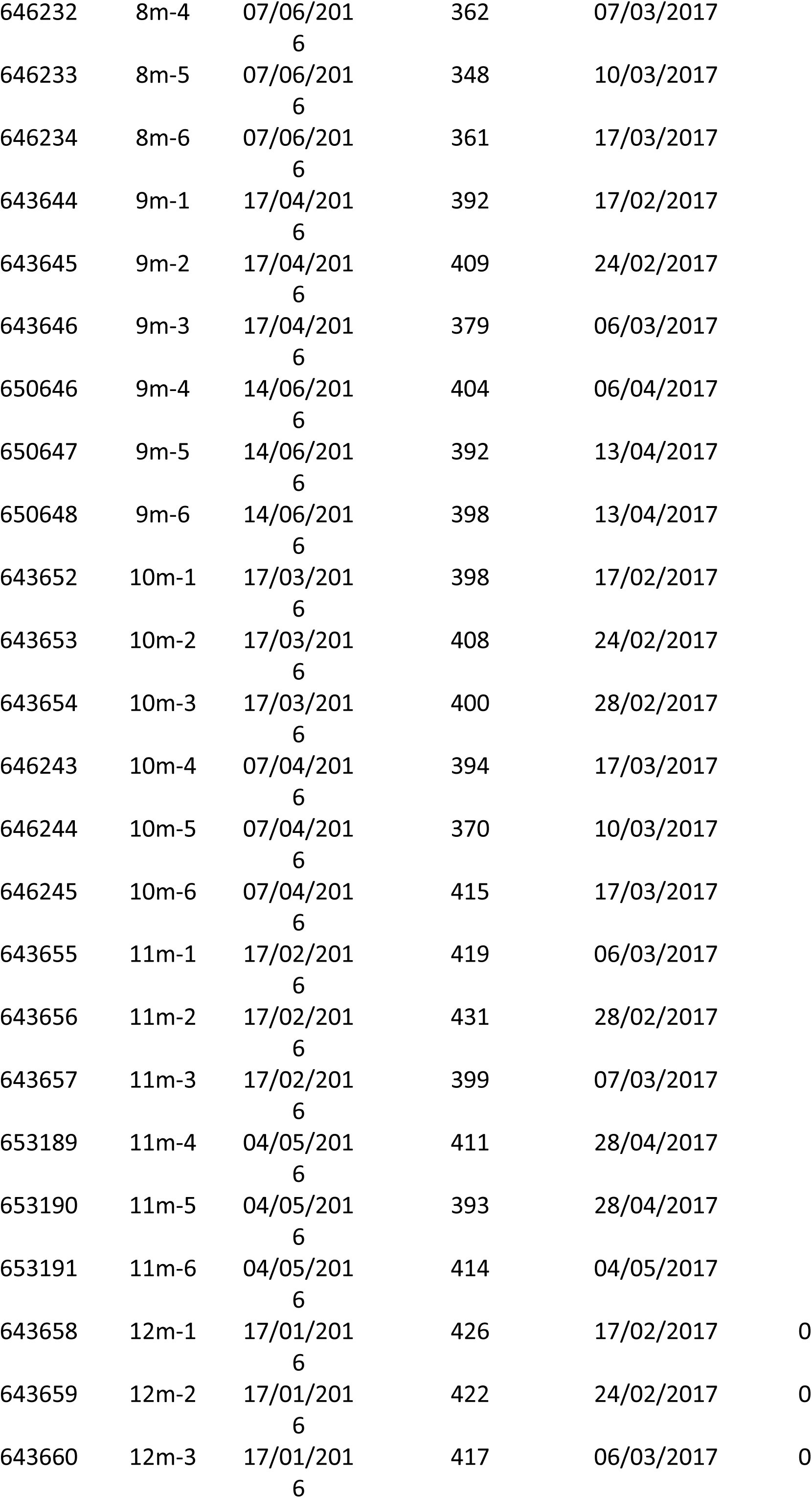
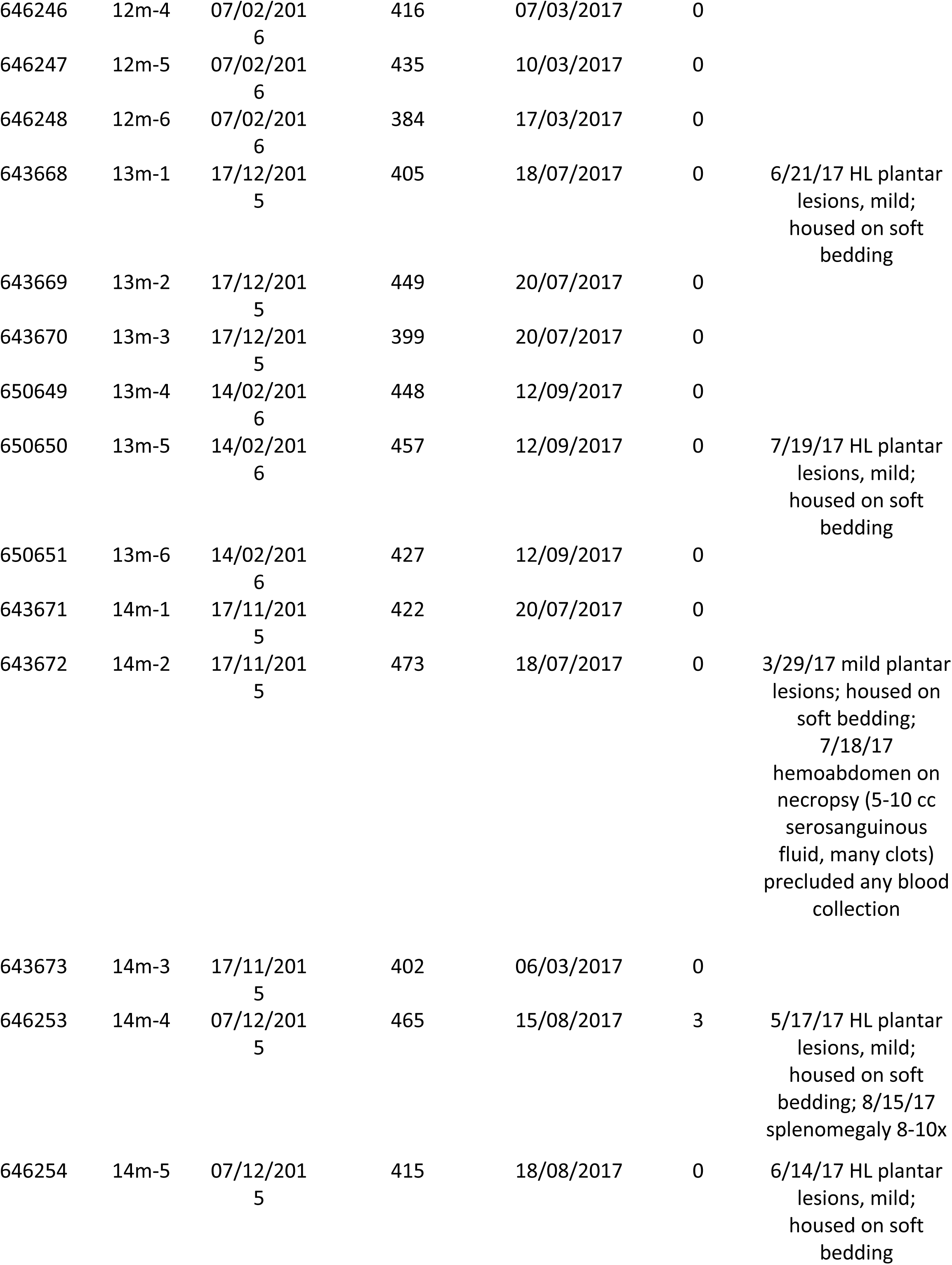
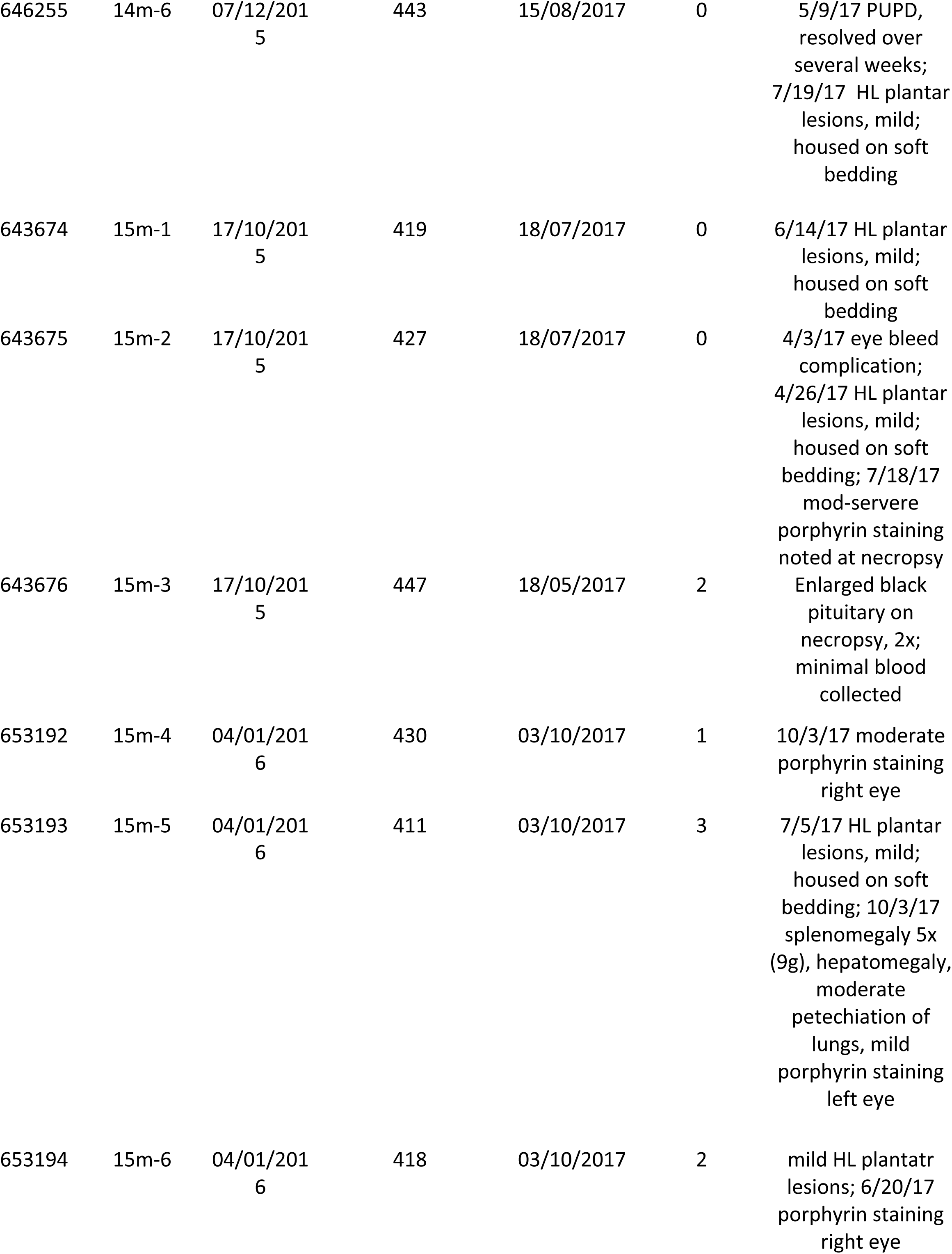
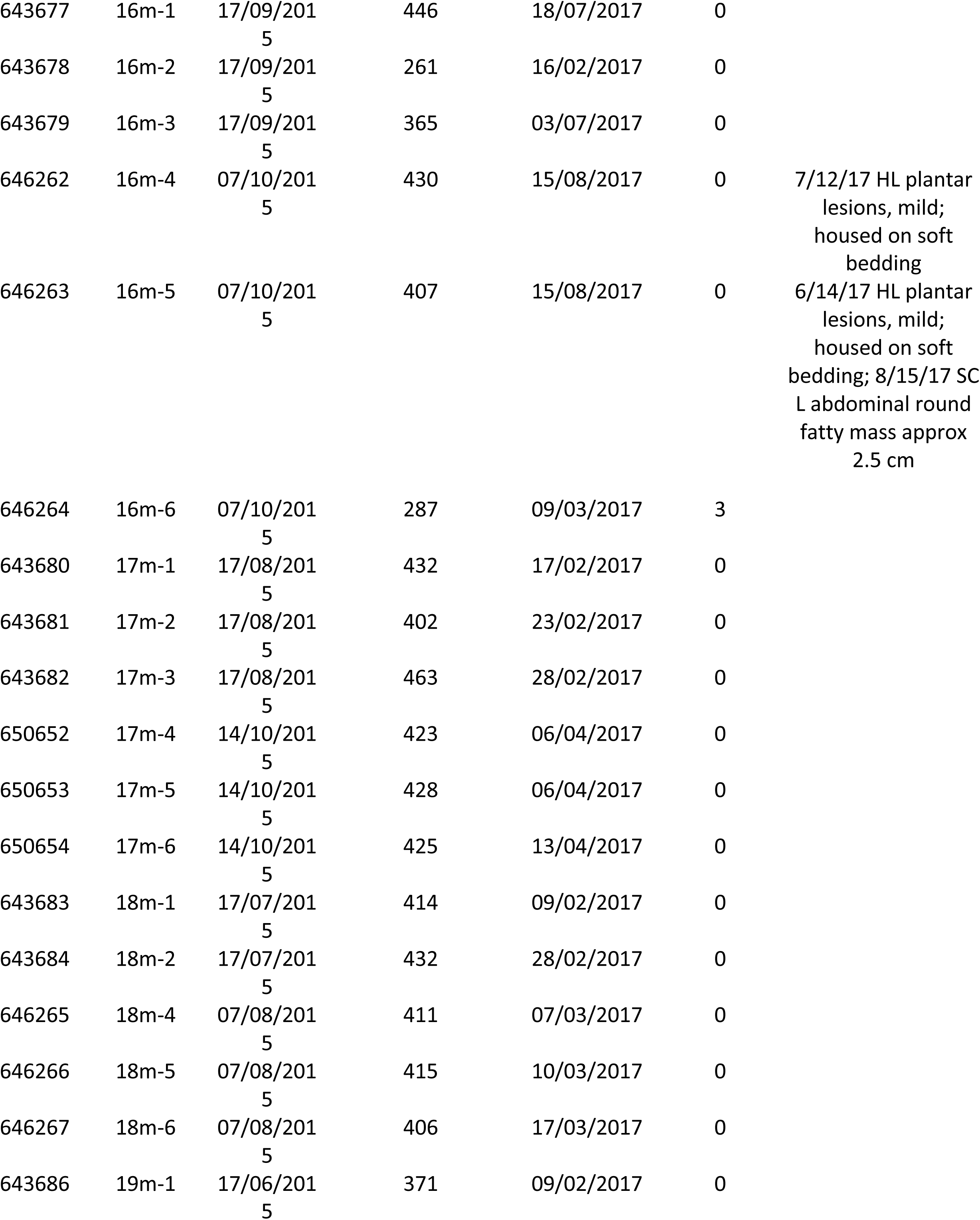
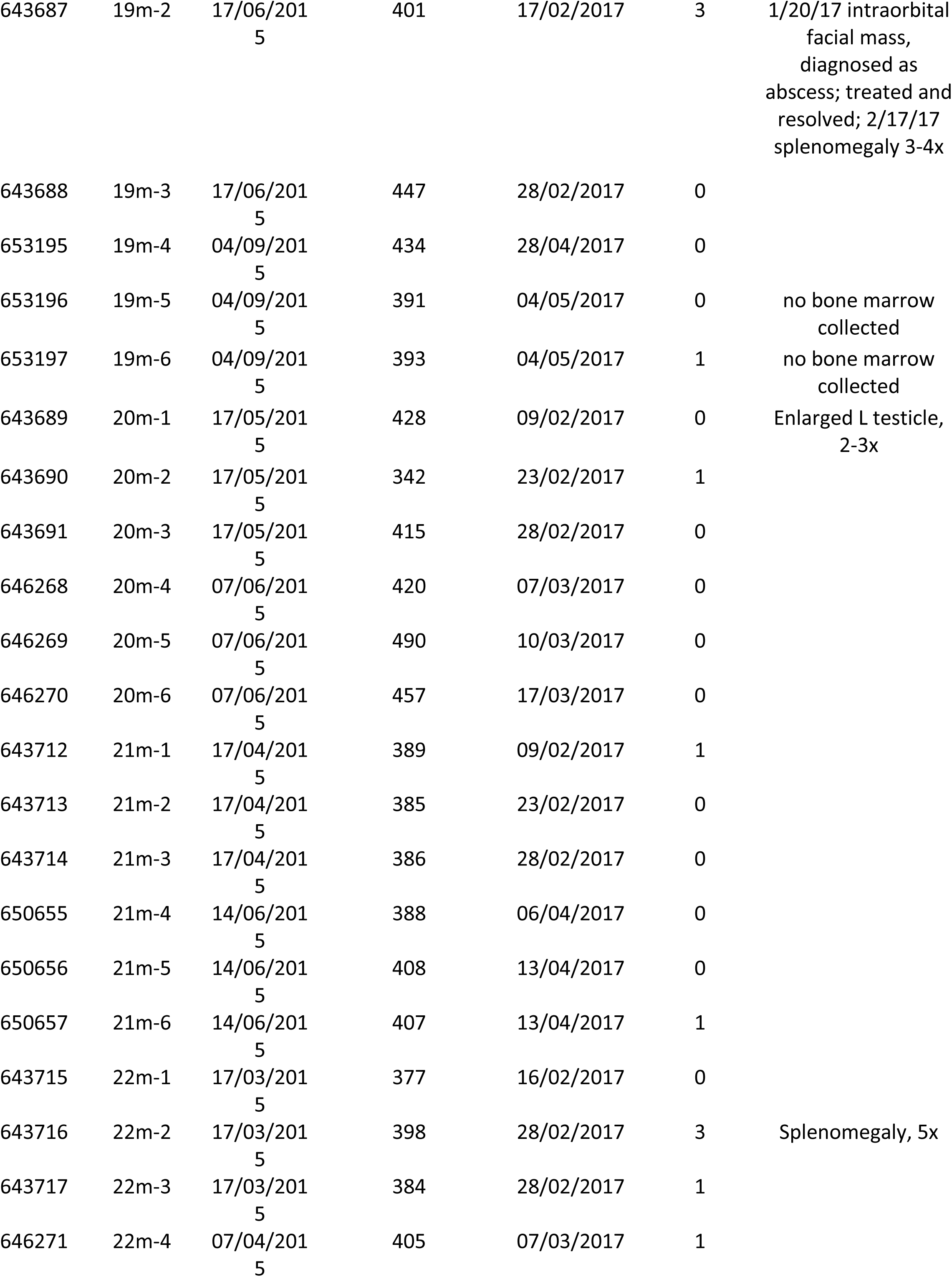
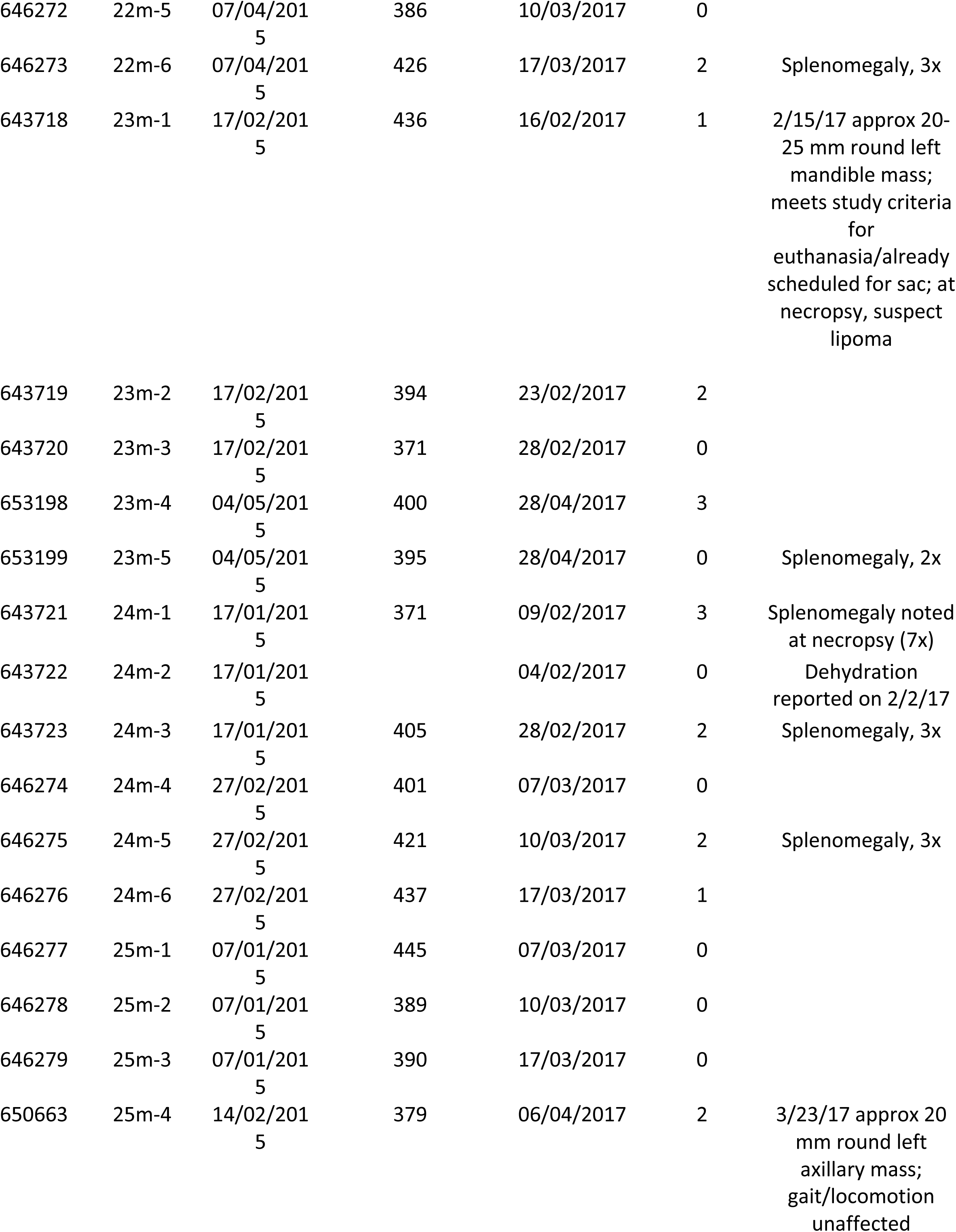
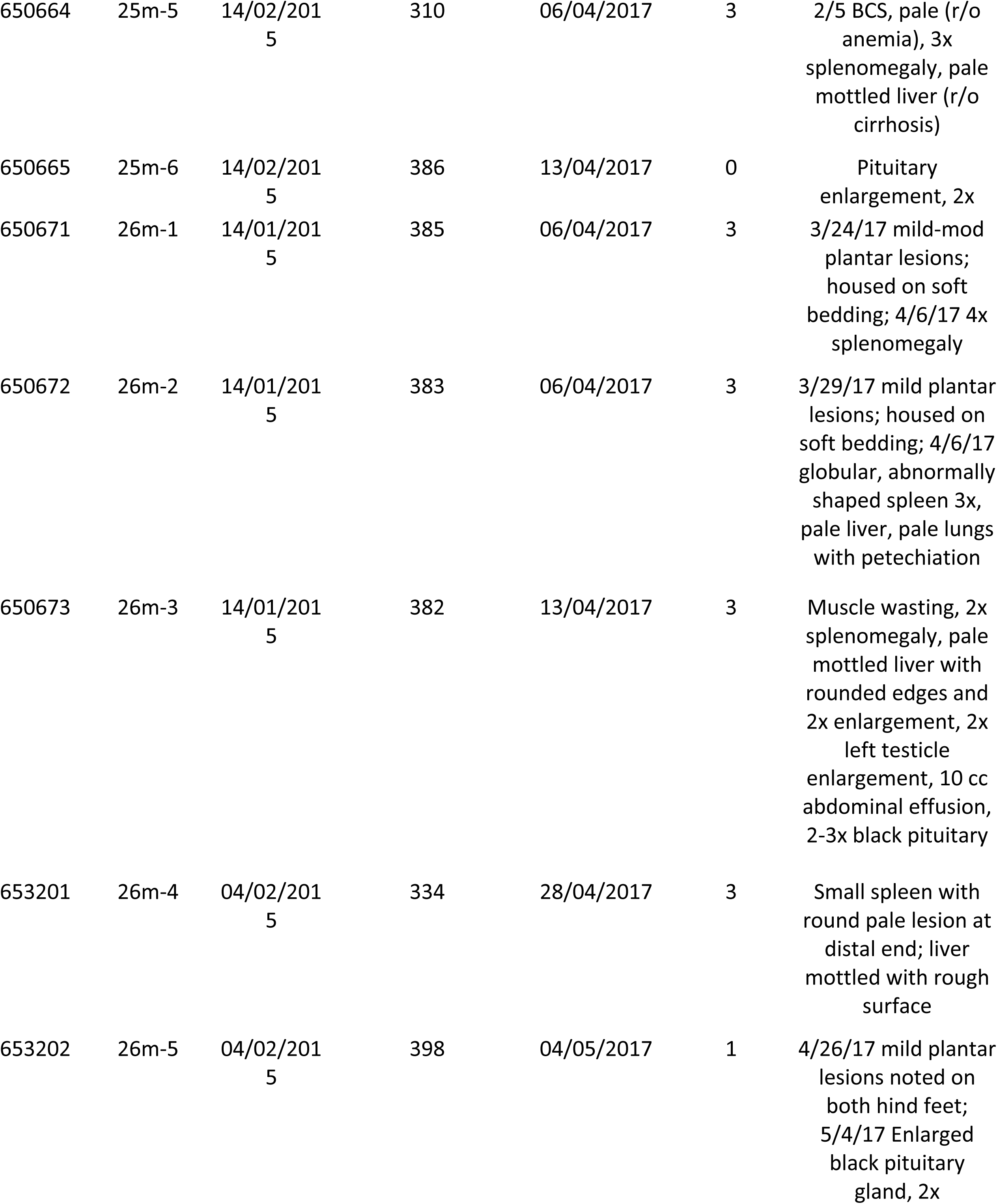
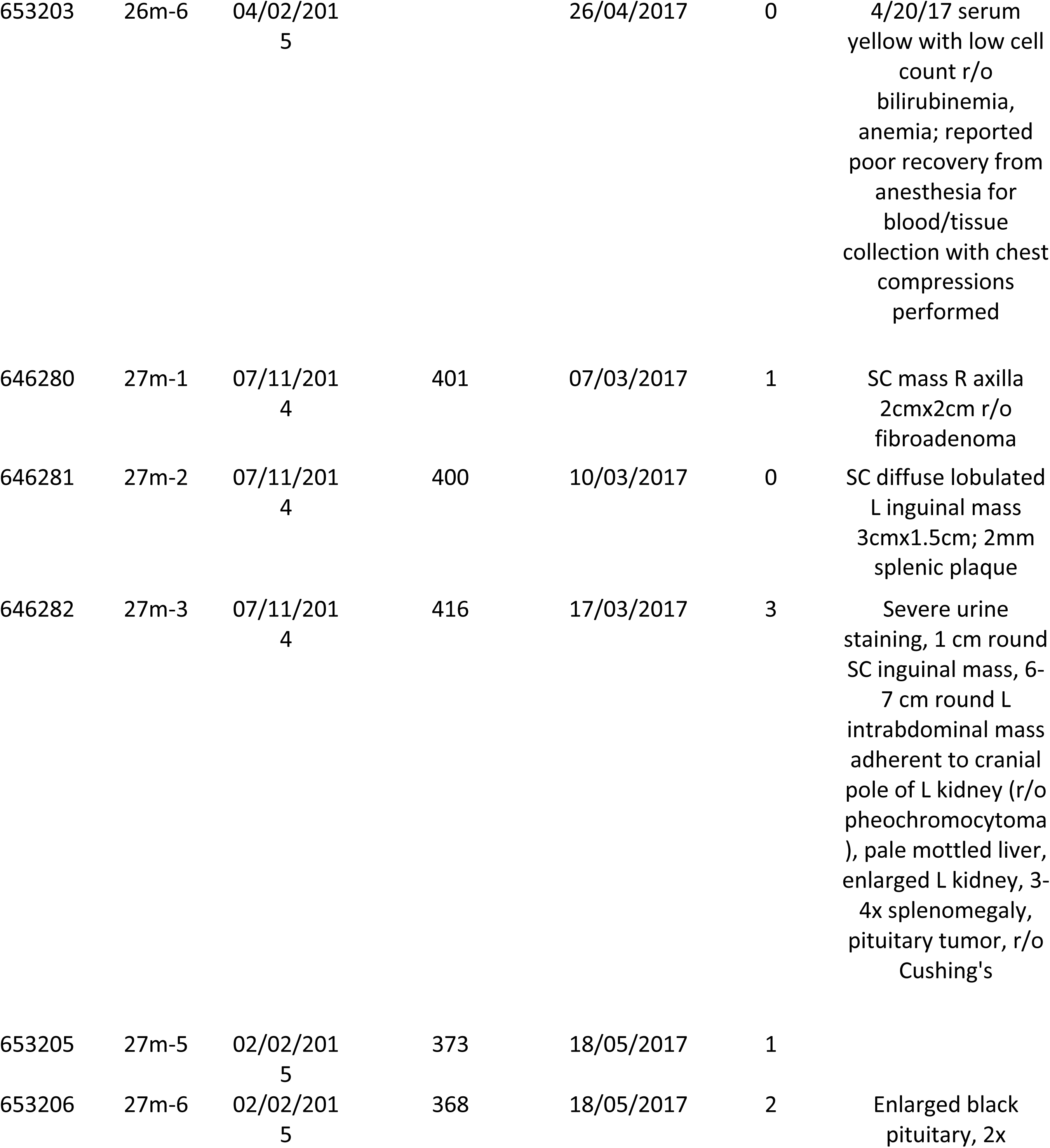
Animal Pathology

| Rat ID | Age (m) | BW (g) | T cells FoL | Neutrophils FoL | B cells FoL | NK cells FoL | monocytes FoL | Stimulated monocytes FoL | CD8+ Freq of T cells | CD4+ Freq of T cells | CD25 + Freq of CD4+ T cells | CD4+/CD25 + T cells FoL | CD4+/CD25- T cells FoL | CD8+ T cells FoL | Myeloid/lymphoid ratio | CD25 + Freq of T cells |
| --- | --- | --- | --- | --- | --- | --- | --- | --- | --- | --- | --- | --- | --- | --- | --- | --- |
| 3m-1 | 3.20 | 270 | 58.10 | 3.01 | 26.80 | 4.07 | 4.07 | 0.60 | 34.95 | 65.05 | 4.89 | 1.74 | 33.80 | 19.30 | 0.09 | 0.03 |
| 3m-2 | 3.50 | 303 | 63.20 | 0.98 | 26.40 | 3.71 | 3.11 | 0.40 | 37.01 | 62.99 | 6.40 | 2.44 | 35.70 | 22.70 | 0.05 | 0.04 |
| 3m-3 | 3.60 | 310 | 67.80 | 2.34 | 15.90 | 5.43 | 4.09 | 0.43 | 34.76 | 65.24 | 5.52 | 2.33 | 39.80 | 22.60 | 0.08 | 0.04 |
| 3m-4 | 3.20 | 294 | 61.30 | 2.39 | 28.70 | 2.51 | 2.23 | 0.39 | 35.20 | 64.80 | 5.94 | 2.21 | 35.00 | 20.40 | 0.05 | 0.04 |
| 3m-5 | 3.20 | 283 | 55.30 | 2.81 | 26.70 | 4.51 | 4.41 | 0.63 | 35.75 | 64.25 | 5.87 | 1.94 | 31.00 | 18.50 | 0.09 | 0.04 |
| 3m-6 | 3.50 | 311 | 59.60 | 2.21 | 28.30 | 2.83 | 2.97 | 0.42 | 36.84 | 63.16 | 7.21 | 2.58 | 33.20 | 21.10 | 0.06 | 0.05 |
| 4m-1 | 4.50 | 313 | 62.60 | 1.31 | 23.90 | 6.12 | 2.78 | 0.36 | 38.14 | 61.86 | 6.38 | 2.37 | 34.80 | 23.20 | 0.05 | 0.04 |
| 4m-2 | 4.50 | 314 | 63.50 | 2.45 | 23.20 | 4.48 | 2.73 | 0.80 | 39.00 | 61.00 | 7.52 | 2.81 | 34.50 | 24.20 | 0.07 | 0.05 |
| 4m-3 | 4.50 | 317 | 62.40 | 3.82 | 19.00 | 4.08 | 5.43 | 0.64 | 37.11 | 62.89 | 6.36 | 2.39 | 35.20 | 22.50 | 0.12 | 0.04 |
| 4m-4 | 4.10 | 269 | 59.40 | 2.68 | 29.60 | 2.17 | 2.32 | 0.81 | 37.79 | 62.21 | 6.49 | 2.25 | 32.40 | 21.30 | 0.06 | 0.04 |
| 4m-5 | 4.20 | 401 | 63.80 | 3.03 | 25.50 | 3.00 | 1.90 | 0.34 | 35.77 | 64.23 | 5.85 | 2.28 | 36.60 | 21.90 | 0.06 | 0.04 |
| 4m-6 | 4.50 | 288 | 70.40 | 1.85 | 19.30 | 2.63 | 2.83 | 0.27 | 34.62 | 65.38 | 5.76 | 2.37 | 38.80 | 22.10 | 0.05 | 0.04 |
| 5m-1 | 5.20 | 350 | 58.10 | 2.45 | 24.10 | 5.20 | 4.43 | 0.67 | 38.85 | 61.15 | 9.37 | 3.14 | 30.40 | 21.70 | 0.09 | 0.06 |
| 5m-2 | 5.50 | 340 | 65.20 | 2.19 | 21.70 | 3.42 | 3.64 | 0.76 | 38.75 | 61.25 | 8.74 | 3.36 | 35.00 | 24.70 | 0.07 | 0.05 |
| 5m-3 | 5.70 | 356 | 58.50 | 2.52 | 26.30 | 5.12 | 3.30 | 0.75 | 39.48 | 60.52 | 8.19 | 2.74 | 30.80 | 22.20 | 0.07 | 0.05 |
| 5m-4 | 5.30 | 337 | 62.80 | 4.50 | 14.70 | 5.49 | 6.04 | 0.71 | 35.68 | 64.32 | 7.76 | 2.85 | 33.90 | 20.70 | 0.14 | 0.05 |
| 5m-5 | 5.50 | 369 | 58.70 | 3.12 | 24.00 | 3.86 | 4.71 | 0.56 | 37.01 | 62.99 | 7.56 | 2.61 | 31.80 | 20.50 | 0.10 | 0.05 |
| 5m-6 | 5.50 | 334 | 57.90 | 5.04 | 20.60 | 5.82 | 5.30 | 0.74 | 34.93 | 65.07 | 5.38 | 1.89 | 33.20 | 19.10 | 0.13 | 0.04 |
| 6m-1 | 6.30 | 385 | 58.00 | 3.06 | 24.40 | 4.91 | 4.82 | 0.62 | 37.59 | 62.41 | 7.97 | 2.76 | 31.80 | 21.20 | 0.10 | 0.05 |
| 6m-2 | 6.50 | 360 | 59.20 | 2.79 | 27.60 | 3.73 | 3.00 | 0.87 | 37.55 | 62.45 | 8.80 | 3.13 | 32.50 | 21.80 | 0.07 | 0.05 |
| 6m-3 | 6.70 | 376 | 59.00 | 3.32 | 24.00 | 3.98 | 4.72 | 0.59 | 36.72 | 63.28 | 9.16 | 3.24 | 32.10 | 20.90 | 0.10 | 0.06 |
| 6m-4 | 6.20 | 370 | 63.30 | 3.76 | 23.80 | 4.25 | 1.55 | 0.39 | 35.96 | 64.04 | 7.14 | 2.69 | 34.90 | 21.40 | 0.06 | 0.05 |
| 6m-5 | 6.30 | 373 | 60.50 | 3.42 | 22.60 | 5.33 | 3.46 | 0.40 | 34.24 | 65.76 | 7.26 | 2.72 | 34.80 | 19.80 | 0.08 | 0.05 |
| 6m-6 | 6.50 | 312 | 60.90 | 4.46 | 15.20 | 6.22 | 6.76 | 0.85 | 33.59 | 66.41 | 5.53 | 2.01 | 34.40 | 18.70 | 0.15 | 0.04 |
| 7m-1 | 7.20 | 363 | 54.10 | 2.85 | 26.80 | 4.71 | 6.14 | 0.87 | 37.10 | 62.90 | 8.30 | 2.68 | 29.60 | 19.40 | 0.12 | 0.05 |
| 7m-2 | 7.50 | 385 | 62.30 | 1.31 | 25.90 | 4.28 | 3.17 | 0.49 | 37.25 | 62.75 | 10.59 | 3.92 | 33.00 | 22.30 | 0.05 | 0.07 |
| 7m-3 | 7.70 | 351 | 55.70 | 2.88 | 25.20 | 6.70 | 5.03 | 0.58 | 37.19 | 62.81 | 9.84 | 3.28 | 30.10 | 20.00 | 0.10 | 0.06 |
| 7m-4 | 7.20 | 365 | 63.20 | 2.13 | 22.90 | 4.47 | 2.82 | 0.44 | 35.20 | 64.80 | 10.37 | 4.04 | 34.90 | 21.50 | 0.06 | 0.07 |
| 7m-5 | 7.50 | 365 | 60.60 | 4.69 | 18.70 | 4.94 | 5.48 | 0.63 | 37.13 | 62.87 | 9.70 | 3.47 | 32.30 | 21.50 | 0.13 | 0.06 |
| 7m-6 | 7.50 | 390 | 57.40 | 1.60 | 29.70 | 3.63 | 3.33 | 0.28 | 39.44 | 60.56 | 9.92 | 3.25 | 29.50 | 21.70 | 0.06 | 0.06 |
| 8m-1 | 8.30 | 408 | 48.10 | 2.94 | 27.50 | 6.41 | 8.11 | 1.37 | 37.93 | 62.07 | 9.69 | 2.71 | 25.30 | 17.50 | 0.15 | 0.06 |
| 8m-2 | 8.60 | 419 | 57.00 | 3.38 | 29.20 | 2.66 | 3.51 | 0.69 | 41.64 | 58.36 | 11.52 | 3.67 | 28.20 | 23.20 | 0.09 | 0.07 |
| 8m-3 | 8.70 | 426 | 57.50 | 4.35 | 23.10 | 4.79 | 4.46 | 0.65 | 36.48 | 63.52 | 9.44 | 3.19 | 30.60 | 19.80 | 0.11 | 0.06 |
| 8m-4 | 8.20 | 352 | 56.50 | 5.88 | 26.40 | 2.56 | 3.84 | 0.84 | 39.16 | 60.84 | 9.54 | 2.99 | 28.40 | 20.60 | 0.12 | 0.06 |
| 8m-5 | 8.30 | 348 | 52.70 | 5.23 | 18.20 | 9.90 | 6.36 | 1.15 | 35.84 | 64.16 | 7.74 | 2.39 | 28.50 | 17.50 | 0.16 | 0.05 |
| 8m-6 | 8.50 | 360 | 61.70 | 5.77 | 7.36 | 9.07 | 8.27 | 0.72 | 35.11 | 64.89 | 5.59 | 1.86 | 31.40 | 18.30 | 0.19 | 0.04 |
| 9m-1 | 9.50 | 411 | 65.00 | 0.38 | 25.80 | 5.49 | 1.37 | 0.08 | 38.33 | 61.67 | 10.15 | 3.86 | 34.20 | 24.10 | 0.02 | 0.06 |
| 9m-2 | 9.50 | 400 | 55.30 | 4.13 | 27.80 | 4.17 | 3.65 | 0.97 | 39.77 | 60.23 | 11.52 | 3.65 | 28.10 | 21.40 | 0.10 | 0.07 |
| 9m-3 | 9.70 | 378 | 48.90 | 4.84 | 24.60 | 6.74 | 7.28 | 0.91 | 36.88 | 63.13 | 14.23 | 4.13 | 24.80 | 17.30 | 0.16 | 0.09 |
| 9m-4 | 9.30 | 415 | 54.10 | 5.47 | 21.90 | 6.61 | 4.14 | 0.73 | 37.19 | 62.81 | 16.12 | 4.70 | 24.50 | 17.70 | 0.13 | 0.10 |
| 9m-5 | 9.50 | 413 | 53.10 | 3.14 | 23.40 | 8.79 | 5.58 | 0.52 | 32.59 | 67.41 | 14.04 | 4.15 | 25.50 | 14.70 | 0.11 | 0.09 |
| 9m-6 | 9.50 | 409 | 46.80 | 4.91 | 26.40 | 5.62 | 7.35 | 0.65 | 35.15 | 64.85 | 16.24 | 4.40 | 22.70 | 15.10 | 0.16 | 0.11 |
| 10m-1 | 10.50 | 404 | 57.00 | 4.15 | 21.90 | 6.23 | 5.56 | 0.74 | 37.13 | 62.87 | 12.05 | 4.09 | 29.80 | 20.40 | 0.12 | 0.08 |
| 10m-2 | 10.60 | 413 | 55.20 | 3.51 | 24.50 | 4.74 | 7.18 | 0.82 | 39.50 | 60.50 | 12.65 | 4.00 | 27.60 | 21.10 | 0.14 | 0.08 |
| 10m-3 | 10.60 | 407 | 65.90 | 2.96 | 13.80 | 8.12 | 4.54 | 0.36 | 30.83 | 69.17 | 10.16 | 3.89 | 34.40 | 17.30 | 0.09 | 0.07 |
| 10m-4 | 10.20 | 382 | 52.50 | 2.63 | 26.30 | 6.89 | 6.32 | 0.57 | 35.63 | 64.37 | 12.90 | 3.94 | 26.70 | 17.30 | 0.11 | 0.08 |
| 10m-5 | 10.30 | 374 | 53.50 | 4.97 | 21.80 | 5.88 | 5.76 | 0.93 | 36.31 | 63.69 | 9.80 | 3.04 | 28.00 | 18.00 | 0.14 | 0.06 |
| 10m-6 | 10.50 | 423 | 55.30 | 2.69 | 26.40 | 5.67 | 4.97 | 0.60 | 35.66 | 64.34 | 10.73 | 3.49 | 29.00 | 18.40 | 0.09 | 0.07 |
| 11m-1 | 11.70 | 431 | 54.60 | 5.65 | 24.90 | 3.22 | 5.68 | 0.54 | 38.39 | 61.61 | 15.30 | 4.80 | 26.50 | 20.00 | 0.14 | 0.09 |
| 11m-2 | 11.50 | 440 | 48.90 | 3.47 | 29.50 | 3.83 | 8.15 | 0.79 | 37.87 | 62.13 | 15.49 | 4.44 | 24.20 | 18.00 | 0.15 | 0.10 |
| 11m-3 | 11.70 | 409 | 56.20 | 6.56 | 16.50 | 8.82 | 5.42 | 1.18 | 33.23 | 66.77 | 10.50 | 3.69 | 31.50 | 17.90 | 0.16 | 0.07 |
| 11m-4 | 11.20 | 410 | 53.50 | 3.68 | 25.90 | 4.39 | 4.78 | 0.47 | 38.61 | 61.39 | 15.34 | 4.45 | 24.60 | 18.70 | 0.11 | 0.09 |
| 11m-5 | 11.20 | 412 | 49.50 | 3.27 | 27.60 | 6.21 | 5.51 | 0.53 | 38.40 | 61.60 | 11.81 | 3.40 | 25.40 | 18.30 | 0.11 | 0.07 |
| 11m-6 | 11.50 | 430 | 58.80 | 3.46 | 27.40 | 5.47 | 1.85 | 0.36 | 35.67 | 64.33 | 12.77 | 4.59 | 31.30 | 20.30 | 0.06 | 0.08 |
| 12m-1 | 12.50 | 430 | 59.30 | 2.49 | 27.00 | 2.68 | 4.65 | 0.57 | 36.52 | 63.48 | 14.87 | 5.29 | 30.40 | 21.00 | 0.09 | 0.09 |
| 12m-2 | 12.50 | 407 | 44.80 | 6.01 | 30.30 | 5.04 | 6.74 | 1.13 | 39.57 | 60.43 | 16.84 | 4.31 | 21.20 | 17.10 | 0.17 | 0.10 |
| 12m-3 | 12.70 | 429 | 48.60 | 3.65 | 28.10 | 8.14 | 5.30 | 0.84 | 36.07 | 63.93 | 12.42 | 3.65 | 25.70 | 16.90 | 0.12 | 0.08 |
| 12m-4 | 12.10 | 422 | 59.50 | 3.54 | 23.90 | 4.29 | 4.93 | 0.56 | 31.97 | 68.03 | 12.68 | 4.81 | 33.00 | 18.10 | 0.10 | 0.09 |
| 12m-5 | 12.20 | 443 | 57.70 | 3.26 | 24.10 | 4.31 | 5.23 | 0.56 | 39.16 | 60.84 | 12.97 | 4.26 | 28.60 | 21.60 | 0.11 | 0.08 |
| 12m-6 | 12.50 | 390 | 51.00 | 4.11 | 26.70 | 4.42 | 7.68 | 0.64 | 36.01 | 63.99 | 12.99 | 3.93 | 26.30 | 17.40 | 0.15 | 0.08 |
| 13m-1 | 13.30 | 373 | 49.60 | 4.35 | 22.40 | 7.95 | 7.72 | 1.07 | 36.52 | 63.48 | 15.21 | 4.28 | 23.90 | 16.70 | 0.16 | 0.10 |
| 13m-2 | 13.60 | 414 | 53.50 | 4.36 | 29.20 | 2.30 | 5.77 | 0.87 | 36.47 | 63.53 | 16.03 | 5.08 | 26.60 | 18.70 | 0.13 | 0.10 |
| 13m-3 | 13.70 | 415 | 53.00 | 3.32 | 23.90 | 7.34 | 6.64 | 0.90 | 38.09 | 61.91 | 13.79 | 4.10 | 25.60 | 18.80 | 0.13 | 0.09 |
| 13m-4 | 13.30 | 413 | 55.20 | 5.75 | 18.30 | 7.00 | 6.14 | 0.76 | 39.49 | 60.51 | 12.26 | 3.64 | 26.10 | 19.80 | 0.16 | 0.07 |
| 13m-5 | 13.30 | 417 | 58.90 | 3.60 | 25.00 | 3.32 | 4.73 | 0.41 | 35.83 | 64.17 | 10.12 | 3.31 | 29.40 | 18.60 | 0.10 | 0.06 |
| 13m-6 | 13.50 | 406 | 52.00 | 4.35 | 27.70 | 4.94 | 5.23 | 0.52 | 35.36 | 64.64 | 8.63 | 2.72 | 28.80 | 17.50 | 0.12 | 0.06 |
| 14m-1 | 14.50 | 389 | 51.70 | 2.78 | 26.00 | 6.38 | 6.59 | 1.25 | 36.81 | 63.19 | 15.49 | 4.67 | 25.50 | 18.00 | 0.13 | 0.10 |
| 14m-2 | 14.50 | 436 | 46.40 | 2.86 | 33.90 | 3.61 | 7.71 | 0.63 | 39.47 | 60.53 | 19.92 | 5.12 | 20.60 | 17.40 | 0.13 | 0.12 |
| 14m-3 | 14.70 | 420 | 46.70 | 7.35 | 12.90 | 14.60 | 8.82 | 1.94 | 33.26 | 66.74 | 13.55 | 3.82 | 24.40 | 14.50 | 0.24 | 0.09 |
| 14m-4 | 14.20 | 456 | 42.70 | 6.06 | 29.10 | 4.69 | 8.15 | 1.49 | 34.57 | 65.43 | 13.09 | 3.25 | 21.60 | 13.50 | 0.21 | 0.09 |
| 14m-5 | 14.30 | 426 | 43.00 | 5.73 | 22.50 | 8.12 | 9.78 | 0.98 | 36.62 | 63.38 | 16.99 | 4.09 | 20.00 | 14.30 | 0.22 | 0.11 |
| 14m-6 | 14.50 | 450 | 38.30 | 8.59 | 12.80 | 15.70 | 14.10 | 1.36 | 38.09 | 61.91 | 16.77 | 3.51 | 17.50 | 13.30 | 0.36 | 0.10 |
| 15m-1 | 15.30 | 401 | 45.80 | 4.45 | 29.90 | 4.69 | 8.57 | 1.06 | 35.61 | 64.39 | 18.14 | 4.95 | 22.30 | 15.50 | 0.18 | 0.12 |
| 15m-2 | 15.60 | 430 | 47.90 | 3.01 | 31.70 | 4.58 | 7.56 | 0.70 | 35.13 | 64.87 | 17.75 | 5.16 | 23.90 | 16.30 | 0.13 | 0.12 |
| 15m-3 | 15.70 | 438 | 48.00 | 3.79 | 31.40 | 3.92 | 5.91 | 1.12 | 40.10 | 59.90 | 16.70 | 4.49 | 22.50 | 18.50 | 0.13 | 0.10 |
| 15m-4 | 15.20 | 436 | 53.90 | 3.93 | 21.10 | 5.24 | 7.29 | 0.85 | 38.12 | 61.88 | 16.02 | 4.80 | 25.30 | 19.00 | 0.15 | 0.10 |
| 15m-5 | 15.50 | 418 | 46.50 | 7.83 | 18.30 | 7.58 | 10.10 | 0.73 | 35.37 | 64.63 | 18.14 | 4.93 | 22.30 | 15.40 | 0.26 | 0.12 |
| 15m-6 | 15.50 | 421 | 46.30 | 6.34 | 22.90 | 7.52 | 8.21 | 0.97 | 39.67 | 60.33 | 20.74 | 5.40 | 20.70 | 17.70 | 0.20 | 0.13 |
| 16m-1 | 16.30 | 436 | 43.90 | 3.93 | 33.40 | 4.78 | 7.43 | 1.02 | 34.71 | 65.29 | 20.37 | 5.34 | 20.80 | 14.40 | 0.15 | 0.13 |
| 16m-2 | 16.50 | 388 | 51.20 | 4.73 | 19.30 | 5.89 | 5.97 | 4.20 | 35.51 | 64.49 | 15.72 | 4.88 | 26.20 | 17.60 | 0.20 | 0.10 |
| 16m-3 | 16.70 | 448 | 49.80 | 3.44 | 27.30 | 5.30 | 7.35 | 0.81 | 36.19 | 63.81 | 17.57 | 5.21 | 24.40 | 17.20 | 0.14 | 0.11 |
| 16m-4 | 16.20 | 434 | 55.00 | 4.01 | 22.10 | 3.67 | 7.86 | 1.10 | 34.24 | 65.76 | 16.15 | 5.23 | 27.20 | 17.40 | 0.16 | 0.11 |
| 16m-5 | 16.30 | 431 | 54.70 | 4.10 | 20.10 | 5.19 | 7.86 | 1.12 | 32.98 | 67.02 | 18.00 | 6.00 | 27.30 | 16.90 | 0.16 | 0.12 |
| 17m-1 | 17.50 | 452 | 52.30 | 4.98 | 21.90 | 5.89 | 6.18 | 1.70 | 37.31 | 62.69 | 19.65 | 5.86 | 23.90 | 18.30 | 0.16 | 0.12 |
| 17m-2 | 17.60 | 414 | 47.50 | 6.37 | 19.60 | 6.96 | 8.97 | 2.44 | 34.67 | 65.33 | 17.75 | 5.06 | 23.50 | 15.60 | 0.24 | 0.12 |
| 17m-3 | 17.60 | 453 | 52.00 | 5.79 | 11.00 | 10.90 | 10.90 | 1.26 | 32.46 | 67.54 | 14.49 | 4.75 | 27.90 | 16.10 | 0.24 | 0.10 |
| 17m-4 | 17.30 | 441 | 56.40 | 7.17 | 15.10 | 4.90 | 7.45 | 0.86 | 35.10 | 64.90 | 19.50 | 6.00 | 24.80 | 17.20 | 0.20 | 0.13 |
| 17m-5 | 17.30 | 452 | 49.20 | 6.89 | 17.20 | 6.56 | 8.92 | 1.88 | 38.03 | 61.97 | 20.37 | 5.41 | 21.20 | 16.90 | 0.24 | 0.13 |
| 17m-6 | 17.50 | 447 | 47.80 | 4.82 | 15.50 | 13.60 | 9.57 | 0.82 | 35.75 | 64.25 | 19.67 | 5.32 | 21.70 | 15.60 | 0.20 | 0.13 |
| 18m-1 | 18.30 | 399 | 38.30 | 8.37 | 21.20 | 8.70 | 15.00 | 0.96 | 29.06 | 70.94 | 18.03 | 4.47 | 20.30 | 10.50 | 0.36 | 0.13 |
| 18m-2 | 18.60 | 474 | 51.80 | 6.09 | 18.40 | 6.07 | 8.53 | 1.76 | 36.86 | 63.14 | 19.44 | 5.86 | 24.30 | 18.30 | 0.21 | 0.12 |
| 18m-4 | 18.20 | 418 | 46.80 | 6.00 | 23.80 | 6.07 | 8.46 | 0.65 | 40.04 | 59.96 | 16.26 | 4.13 | 21.20 | 17.40 | 0.20 | 0.10 |
| 18m-5 | 18.30 | 418 | 48.80 | 6.64 | 26.10 | 4.61 | 6.35 | 0.83 | 36.67 | 63.33 | 18.16 | 4.67 | 21.00 | 15.40 | 0.17 | 0.12 |
| 18m-6 | 18.60 | 413 | 52.00 | 3.41 | 31.00 | 4.76 | 4.49 | 0.61 | 35.67 | 64.33 | 16.77 | 5.13 | 25.50 | 17.50 | 0.10 | 0.11 |
| 19m-1 | 19.30 | 367 | 52.80 | 7.12 | 18.40 | 4.17 | 7.48 | 1.59 | 34.79 | 65.21 | 19.40 | 6.00 | 25.00 | 17.20 | 0.21 | 0.13 |
| 19m-2 | 19.50 | 409 | 43.30 | 5.04 | 12.90 | 18.00 | 11.80 | 1.57 | 41.46 | 58.54 | 16.60 | 3.65 | 18.30 | 16.20 | 0.25 | 0.10 |
| 19m-3 | 19.60 | 456 | 51.50 | 4.29 | 5.56 | 13.70 | 13.50 | 0.80 | 28.50 | 71.50 | 17.05 | 5.73 | 27.90 | 13.80 | 0.26 | 0.12 |
| 19m-4 | 19.20 | 440 | 50.50 | 4.23 | 18.40 | 8.47 | 8.46 | 1.08 | 34.87 | 65.13 | 21.38 | 6.18 | 22.70 | 16.00 | 0.18 | 0.14 |
| 19m-5 | 19.50 | 409 | 49.90 | 7.73 | 8.09 | 12.20 | 10.60 | 1.26 | 32.92 | 67.08 | 18.53 | 5.33 | 23.50 | 14.70 | 0.28 | 0.12 |
| 19m-6 | 19.50 | 413 | 44.60 | 7.08 | 21.00 | 4.90 | 11.50 | 1.49 | 31.93 | 68.07 | 18.58 | 5.20 | 22.70 | 13.50 | 0.28 | 0.13 |
| 20m-1 | 20.30 | 431 | 45.70 | 9.72 | 11.60 | 4.38 | 14.10 | 3.26 | 36.25 | 63.75 | 14.93 | 3.92 | 22.30 | 15.40 | 0.44 | 0.10 |
| 20m-2 | 20.60 | 375 | 49.20 | 4.87 | 9.66 | 13.50 | 13.50 | 2.47 | 30.04 | 69.96 | 18.61 | 5.90 | 25.80 | 14.10 | 0.29 | 0.13 |
| 20m-3 | 20.60 | 426 | 54.30 | 4.52 | 7.70 | 11.50 | 11.80 | 1.56 | 32.04 | 67.96 | 16.89 | 5.74 | 28.30 | 16.60 | 0.24 | 0.11 |
| 20m-4 | 20.20 | 424 | 57.50 | 3.92 | 18.00 | 3.74 | 6.93 | 2.11 | 30.56 | 69.44 | 12.11 | 4.48 | 32.50 | 16.70 | 0.16 | 0.08 |
| 20m-5 | 20.30 | 491 | 61.60 | 7.35 | 8.49 | 4.91 | 7.78 | 1.51 | 33.21 | 66.79 | 15.21 | 5.12 | 28.60 | 17.20 | 0.22 | 0.10 |
| 20m-6 | 20.60 | 472 | 44.20 | 7.80 | 10.50 | 9.92 | 13.10 | 1.85 | 35.52 | 64.48 | 15.05 | 3.73 | 21.10 | 14.10 | 0.35 | 0.10 |
| 21m-1 | 21.30 | 398 | 55.00 | 7.46 | 9.25 | 4.44 | 9.99 | 2.08 | 34.95 | 65.05 | 18.90 | 5.93 | 25.50 | 17.50 | 0.28 | 0.12 |
| 21m-2 | 21.60 | 407 | 59.30 | 3.69 | 17.00 | 3.65 | 7.94 | 1.43 | 36.50 | 63.50 | 21.45 | 7.32 | 26.80 | 20.30 | 0.16 | 0.14 |
| 21m-3 | 21.60 | 405 | 71.50 | 4.90 | 11.30 | 3.73 | 3.28 | 1.32 | 29.63 | 70.37 | 16.49 | 7.53 | 38.10 | 19.80 | 0.11 | 0.12 |
| 21m-4 | 21.30 | 411 | 55.50 | 7.32 | 7.76 | 7.30 | 11.00 | 1.45 | 31.38 | 68.62 | 22.10 | 6.66 | 23.40 | 14.30 | 0.28 | 0.15 |
| 21m-5 | 21.50 | 418 | 52.20 | 7.68 | 12.40 | 7.93 | 10.00 | 1.28 | 28.25 | 71.75 | 22.30 | 7.21 | 25.00 | 13.10 | 0.26 | 0.16 |
| 21m-6 | 21.50 | 427 | 55.10 | 7.91 | 5.15 | 8.08 | 12.80 | 2.04 | 28.71 | 71.29 | 13.46 | 4.31 | 27.70 | 13.20 | 0.33 | 0.10 |
| 22m-1 | 22.60 | 374 | 53.40 | 7.40 | 9.08 | 6.70 | 9.82 | 3.13 | 31.73 | 68.27 | 19.50 | 6.57 | 27.10 | 16.20 | 0.29 | 0.13 |
| 22m-2 | 22.60 | 414 | 39.20 | 6.28 | 9.75 | 20.80 | 14.50 | 1.78 | 34.99 | 65.01 | 21.44 | 4.83 | 17.70 | 12.70 | 0.32 | 0.14 |
| 22m-3 | 22.60 | 410 | 48.90 | 5.94 | 12.40 | 5.58 | 14.00 | 2.65 | 31.07 | 68.93 | 15.50 | 4.83 | 26.40 | 14.50 | 0.34 | 0.11 |
| 22m-4 | 22.20 | 406 | 49.90 | 9.41 | 8.84 | 7.33 | 11.40 | 1.81 | 33.48 | 66.52 | 18.67 | 5.35 | 23.20 | 14.90 | 0.34 | 0.12 |
| 22m-5 | 22.30 | 392 | 48.70 | 4.24 | 16.70 | 11.80 | 9.15 | 1.25 | 31.73 | 68.27 | 18.84 | 5.52 | 23.80 | 14.10 | 0.19 | 0.13 |
| 22m-6 | 22.60 | 451 | 37.50 | 8.37 | 9.82 | 14.30 | 22.30 | 1.28 | 39.77 | 60.23 | 15.42 | 2.85 | 15.70 | 12.80 | 0.52 | 0.09 |
| 23m-1 | 23.50 | 458 | 49.60 | 6.53 | 7.53 | 12.40 | 12.00 | 2.08 | 27.19 | 72.81 | 16.41 | 5.37 | 27.30 | 12.60 | 0.30 | 0.12 |
| 23m-2 | 23.50 | 397 | 66.10 | 5.71 | 8.36 | 3.74 | 6.75 | 2.36 | 24.39 | 75.61 | 13.23 | 6.10 | 39.90 | 15.20 | 0.19 | 0.10 |
| 23m-3 | 23.60 | 388 | 54.70 | 4.39 | 9.70 | 7.22 | 14.10 | 1.94 | 27.98 | 72.02 | 17.62 | 6.29 | 29.50 | 14.40 | 0.29 | 0.13 |
| 23m-4 | 23.30 | 413 | 42.10 | 5.87 | 11.90 | 11.60 | 14.40 | 1.09 | 33.68 | 66.32 | 24.10 | 5.58 | 17.60 | 12.20 | 0.33 | 0.16 |
| 23m-5 | 23.30 | 407 | 57.20 | 4.94 | 12.80 | 4.72 | 10.20 | 1.20 | 33.80 | 66.20 | 18.01 | 6.06 | 27.60 | 17.80 | 0.22 | 0.12 |
| 24m-1 | 24.20 | 402 | 29.40 | 8.76 | 2.82 | 2.04 | 21.40 | 7.36 | 41.80 | 58.20 | 14.65 | 2.14 | 12.50 | 11.10 | 1.10 | 0.09 |
| 24m-2 | 24.50 | 326 | 27.40 | 0.35 | 38.80 | 27.60 | 7.02 | 0.22 | 33.48 | 66.52 | 11.02 | 1.82 | 14.80 | 8.60 | 0.08 | 0.07 |
| 24m-3 | 24.60 | 420 | 54.10 | 10.20 | 10.40 | 1.71 | 13.20 | 1.52 | 31.72 | 68.28 | 17.01 | 5.75 | 28.00 | 16.30 | 0.38 | 0.12 |
| 24m-4 | 23.50 | 409 | 53.50 | 7.82 | 5.47 | 9.46 | 12.50 | 1.22 | 26.83 | 73.17 | 17.15 | 5.92 | 28.70 | 13.10 | 0.31 | 0.13 |
| 24m-5 | 23.60 | 424 | 45.20 | 8.15 | 6.14 | 3.43 | 22.20 | 1.32 | 30.14 | 69.86 | 16.95 | 4.32 | 21.20 | 11.50 | 0.58 | 0.12 |
| 24m-6 | 23.80 | 390 | 52.40 | 7.32 | 11.30 | 7.49 | 12.00 | 0.73 | 31.57 | 68.43 | 15.76 | 4.97 | 26.60 | 15.00 | 0.28 | 0.11 |
| 25m-1 | 25.20 | 460 | 46.70 | 7.55 | 7.75 | 8.70 | 14.70 | 1.17 | 25.32 | 74.68 | 18.67 | 5.93 | 25.80 | 11.20 | 0.37 | 0.14 |
| 25m-2 | 25.30 | 428 | 55.60 | 3.49 | 8.82 | 8.20 | 12.60 | 2.63 | 30.99 | 69.01 | 17.36 | 5.56 | 26.50 | 15.00 | 0.26 | 0.12 |
| 25m-3 | 25.50 | 404 | 51.40 | 12.10 | 2.77 | 14.30 | 10.50 | 1.44 | 17.22 | 82.78 | 18.21 | 7.07 | 31.70 | 8.49 | 0.35 | 0.15 |
| 25m-4 | 25.30 | 402 | 46.20 | 9.00 | 7.11 | 8.81 | 14.60 | 2.99 | 24.11 | 75.89 | 30.20 | 9.73 | 22.50 | 10.60 | 0.43 | 0.23 |
| 25m-5 | 25.30 | 356 | 4.64 | 11.50 | 0.05 | 62.70 | 5.45 | 2.67 | 38.12 | 61.88 | 27.91 | 0.35 | 0.91 | 0.83 | 0.29 | 0.17 |
| 25m-6 | 25.50 | 402 | 47.40 | 5.46 | 10.20 | 8.39 | 17.00 | 2.63 | 25.81 | 74.19 | 22.18 | 7.19 | 25.30 | 11.70 | 0.38 | 0.16 |
| 26m-1 | 26.30 | 420 | 43.50 | 8.07 | 3.33 | 19.50 | 16.50 | 1.20 | 52.64 | 47.36 | 14.48 | 2.39 | 14.10 | 19.10 | 0.39 | 0.07 |
| 26m-2 | 26.30 | 405 | 18.30 | 11.40 | 3.05 | 7.99 | 36.80 | 2.61 | 35.85 | 64.15 | 27.06 | 2.24 | 6.04 | 4.95 | 1.73 | 0.17 |
| 26m-3 | 26.50 | 399 | 37.40 | 9.82 | 13.70 | 8.01 | 14.00 | 3.95 | 32.00 | 68.00 | 21.52 | 4.66 | 17.00 | 10.80 | 0.47 | 0.15 |
| 26m-4 | 26.20 | 378 | 38.20 | 10.00 | 10.00 | 9.71 | 18.40 | 2.78 | 32.63 | 67.37 | 21.45 | 5.00 | 18.30 | 11.90 | 0.54 | 0.14 |
| 26m-5 | 26.50 | 424 | 44.00 | 7.75 | 10.30 | 14.30 | 10.30 | 2.05 | 26.50 | 73.50 | 19.83 | 5.88 | 23.70 | 11.10 | 0.29 | 0.15 |
| 26m-6 | 26.50 | 336 | 8.04 | 5.97 | 0.80 | 77.50 | 4.46 | 0.33 | 31.89 | 68.11 | 18.61 | 0.80 | 3.48 | 2.07 | 0.12 | 0.13 |
| 27m-1 | 27.20 | 427 | 43.90 | 8.88 | 19.10 | 2.61 | 12.10 | 2.80 | 26.23 | 73.77 | 14.18 | 4.03 | 24.30 | 10.50 | 0.36 | 0.10 |
| 27m-2 | 27.30 | 437 | 48.20 | 7.01 | 8.73 | 4.57 | 16.00 | 3.18 | 30.78 | 69.22 | 11.51 | 3.10 | 23.70 | 12.40 | 0.43 | 0.08 |
| 27m-3 | 27.50 | 434 | 37.90 | 13.30 | 0.47 | 19.50 | 25.20 | 0.69 | 9.85 | 90.15 | 66.05 | 19.70 | 10.10 | 3.54 | 0.68 | 0.60 |
| 27m-5 | 27.00 | 393 | 53.90 | 7.62 | 8.70 | 3.81 | 9.91 | 1.33 | 24.20 | 75.80 | 10.99 | 3.95 | 32.00 | 11.80 | 0.28 | 0.08 |
| 27m-6 | 27.00 | 372 | 37.40 | 11.90 | 8.44 | 4.93 | 20.40 | 2.17 | 31.58 | 68.42 | 20.27 | 4.61 | 18.20 | 11.00 | 0.68 | 0.14 |

**Suppl. Table 3.**
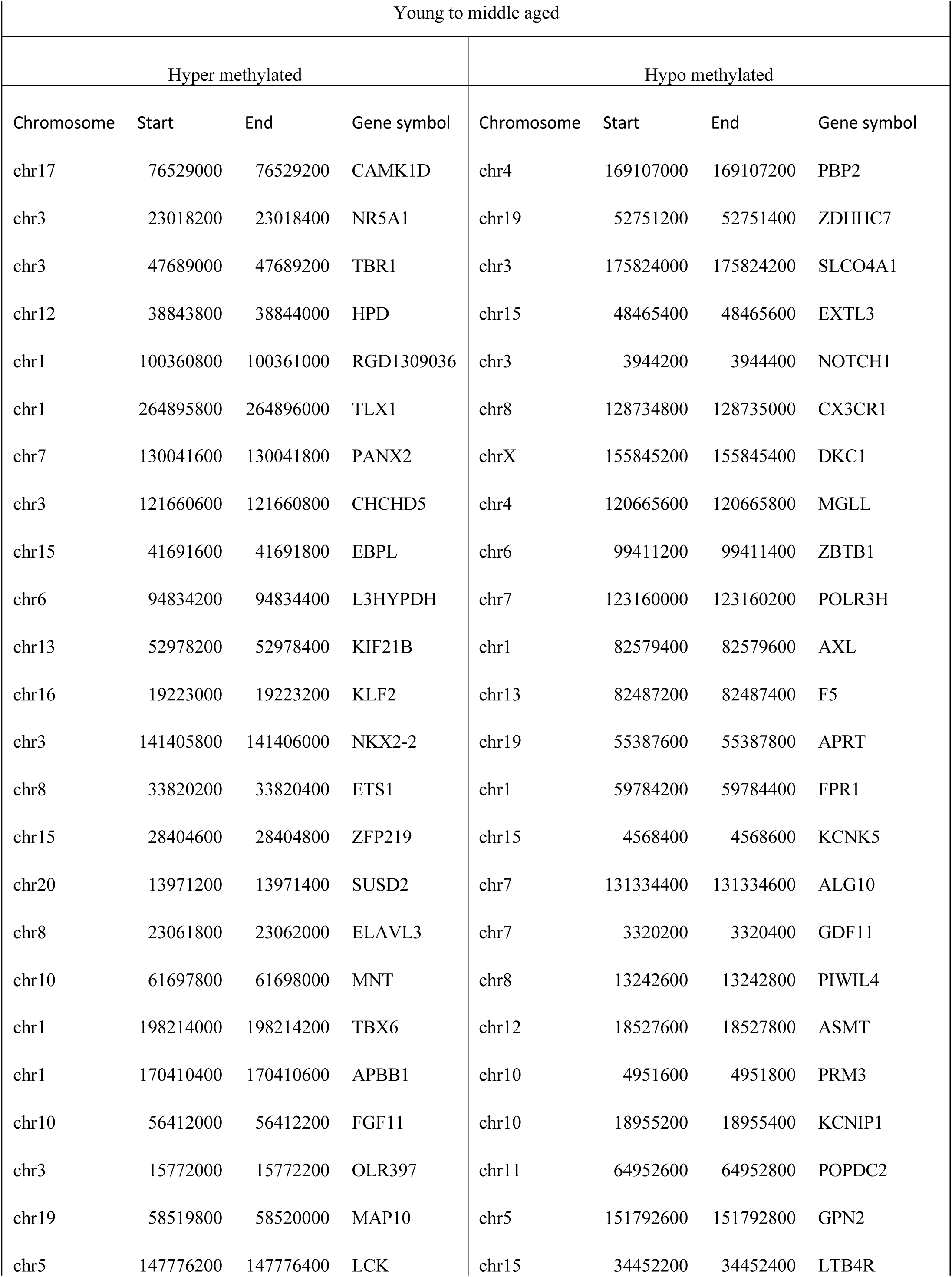
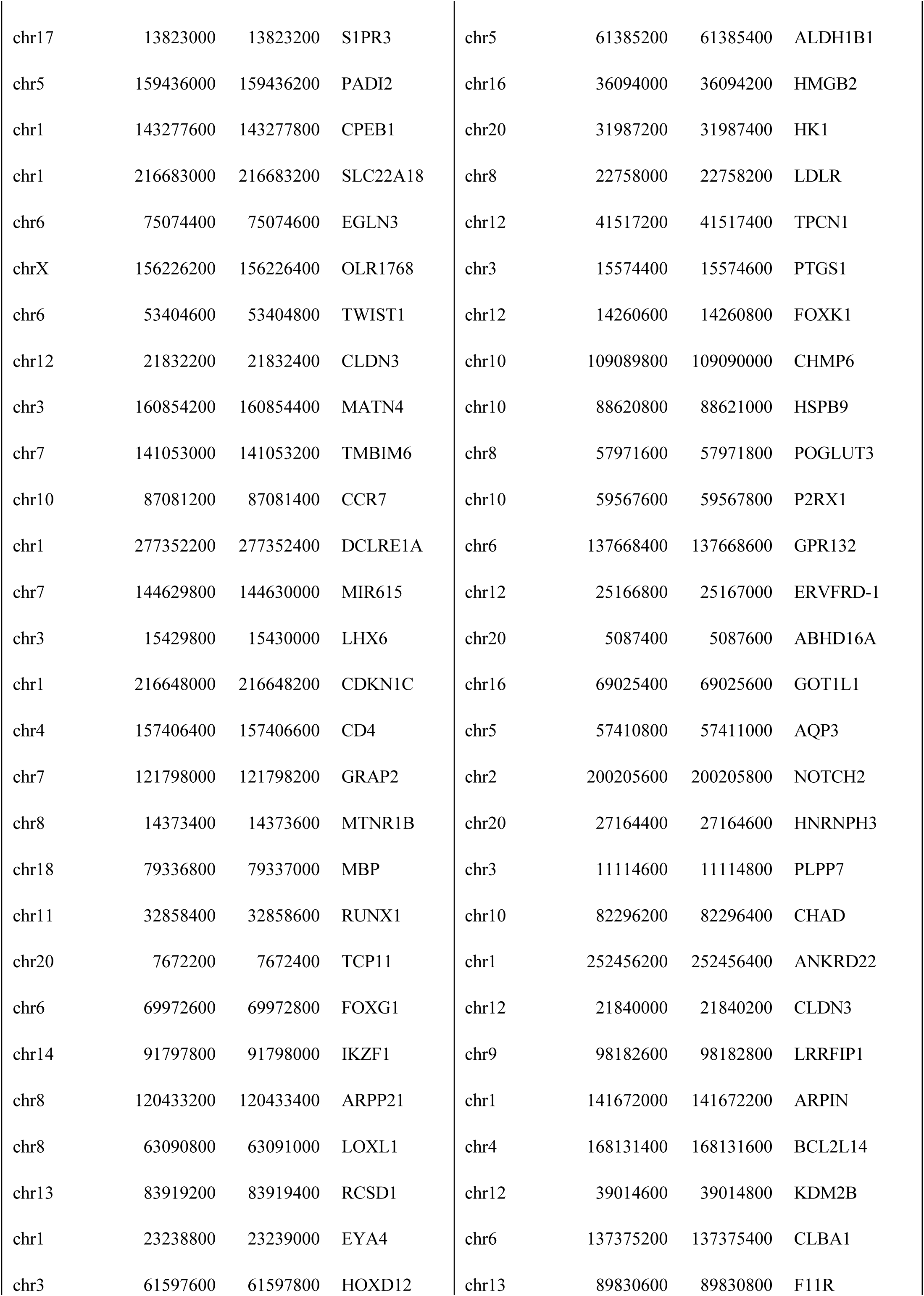
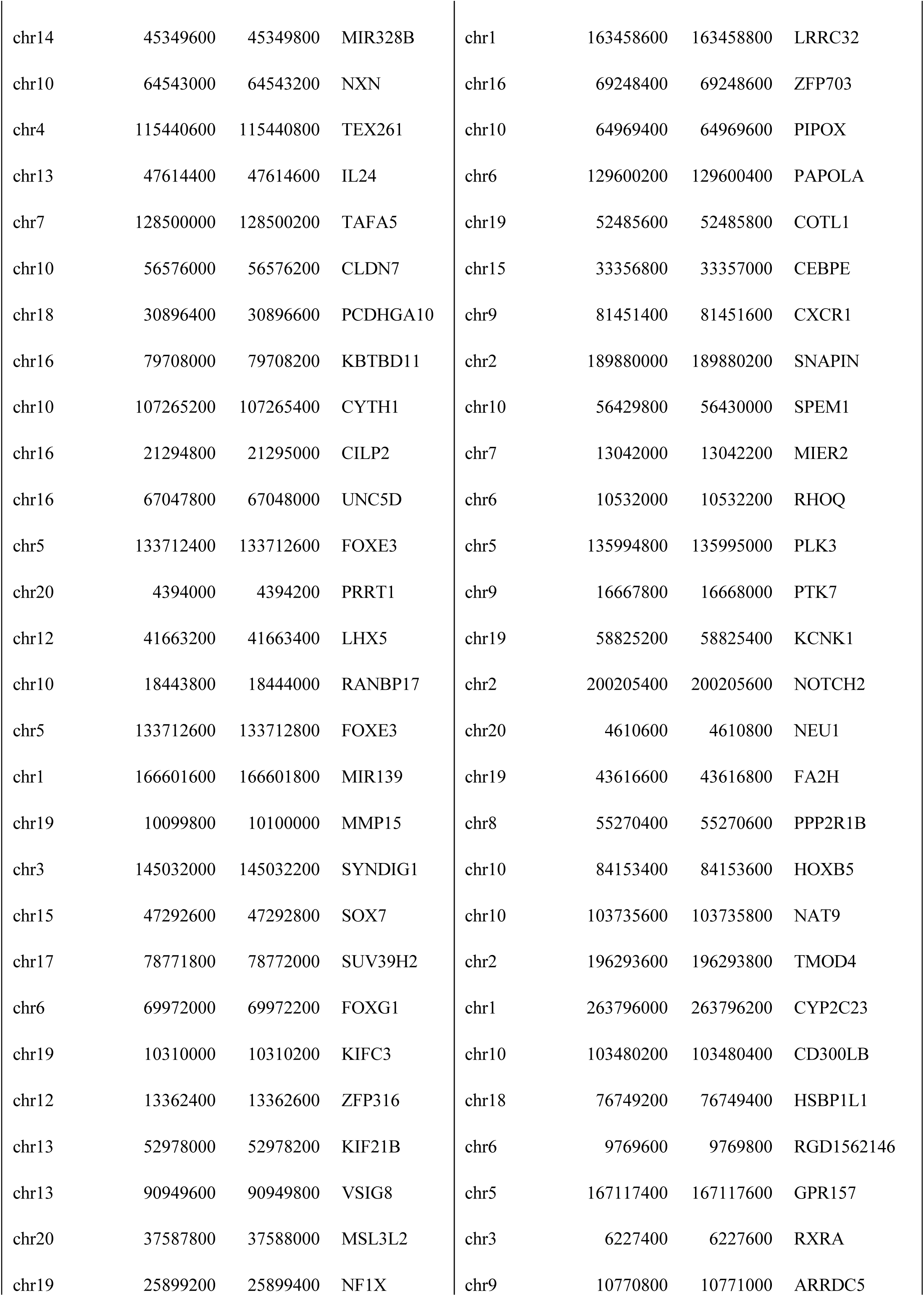
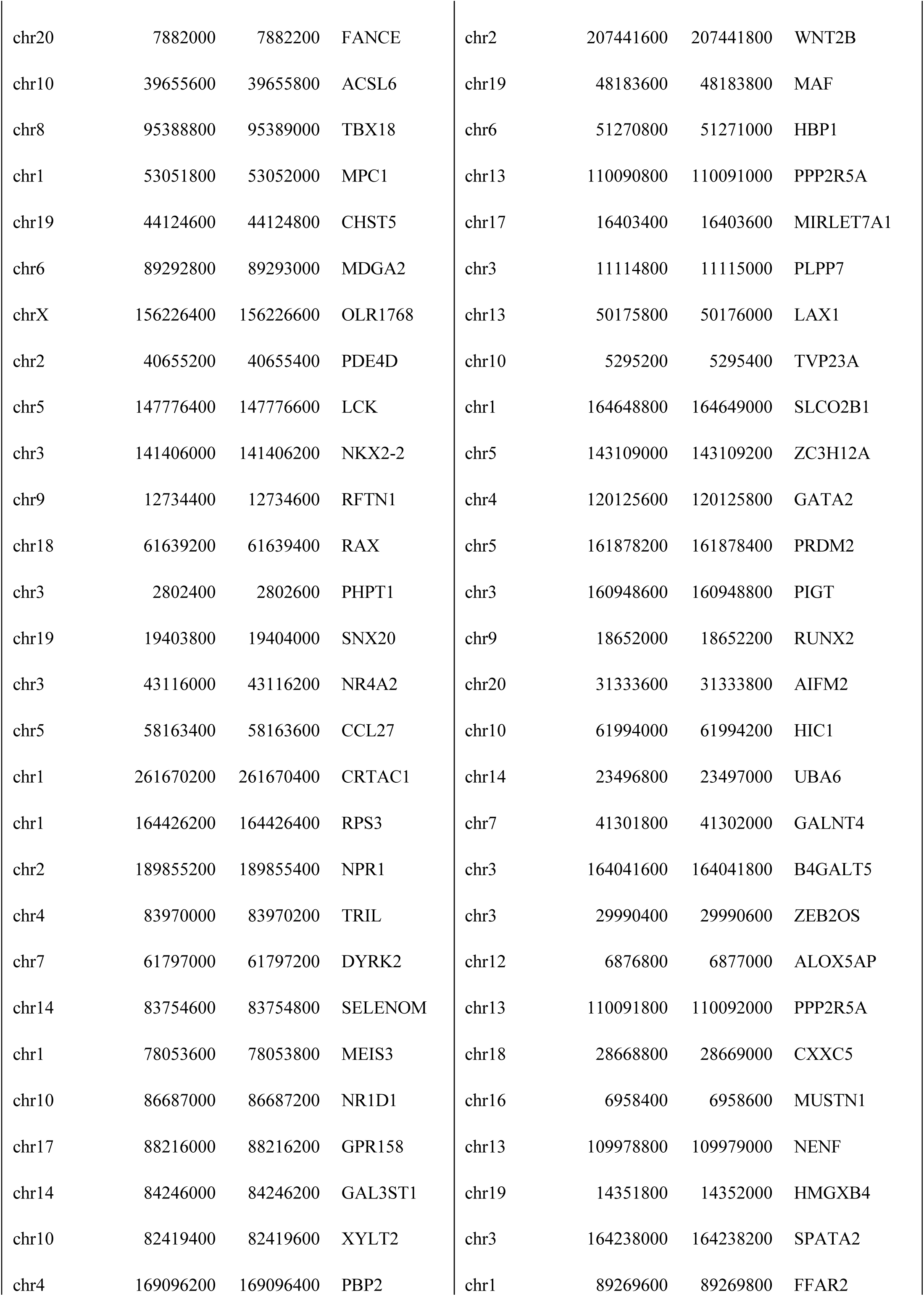
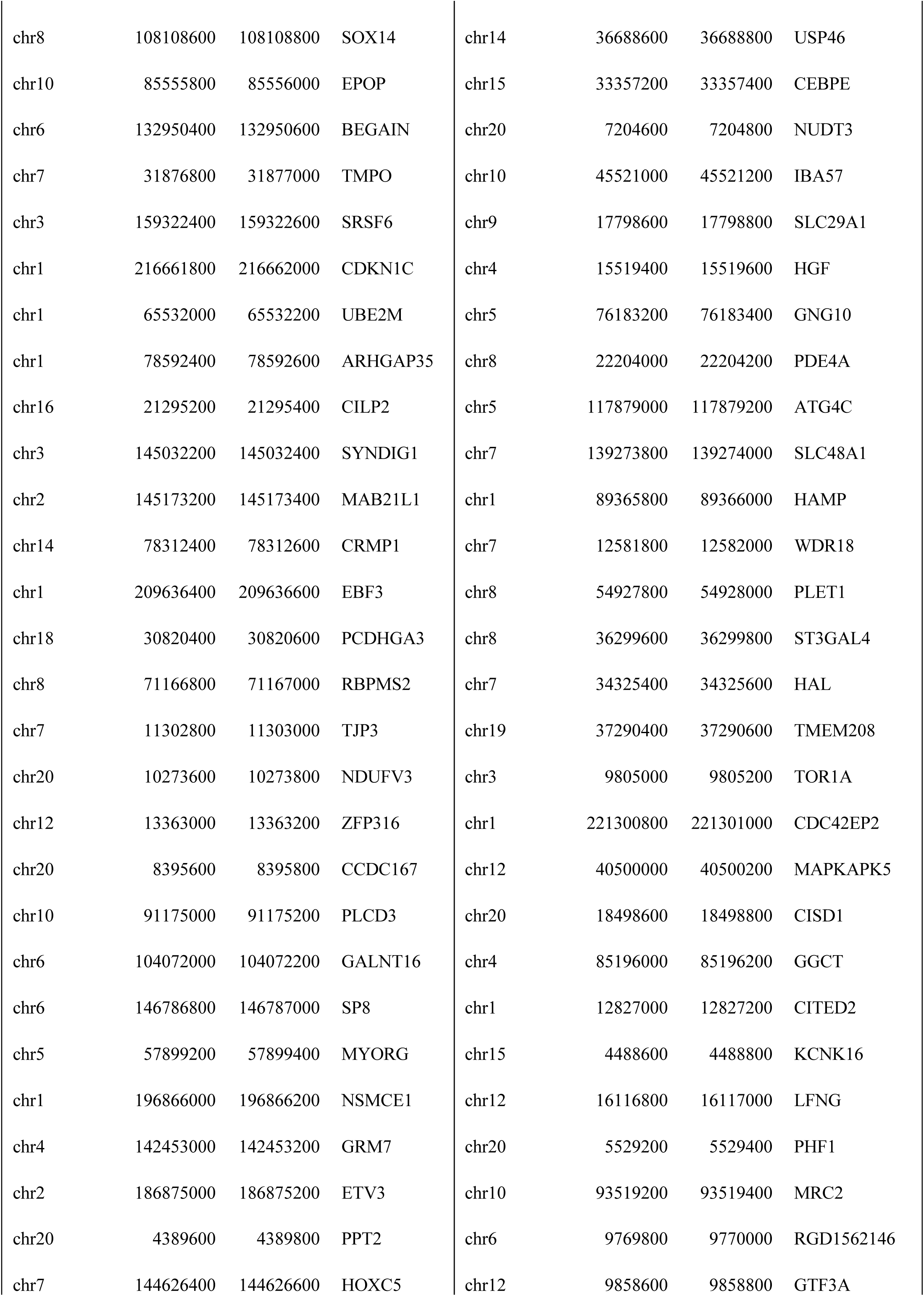
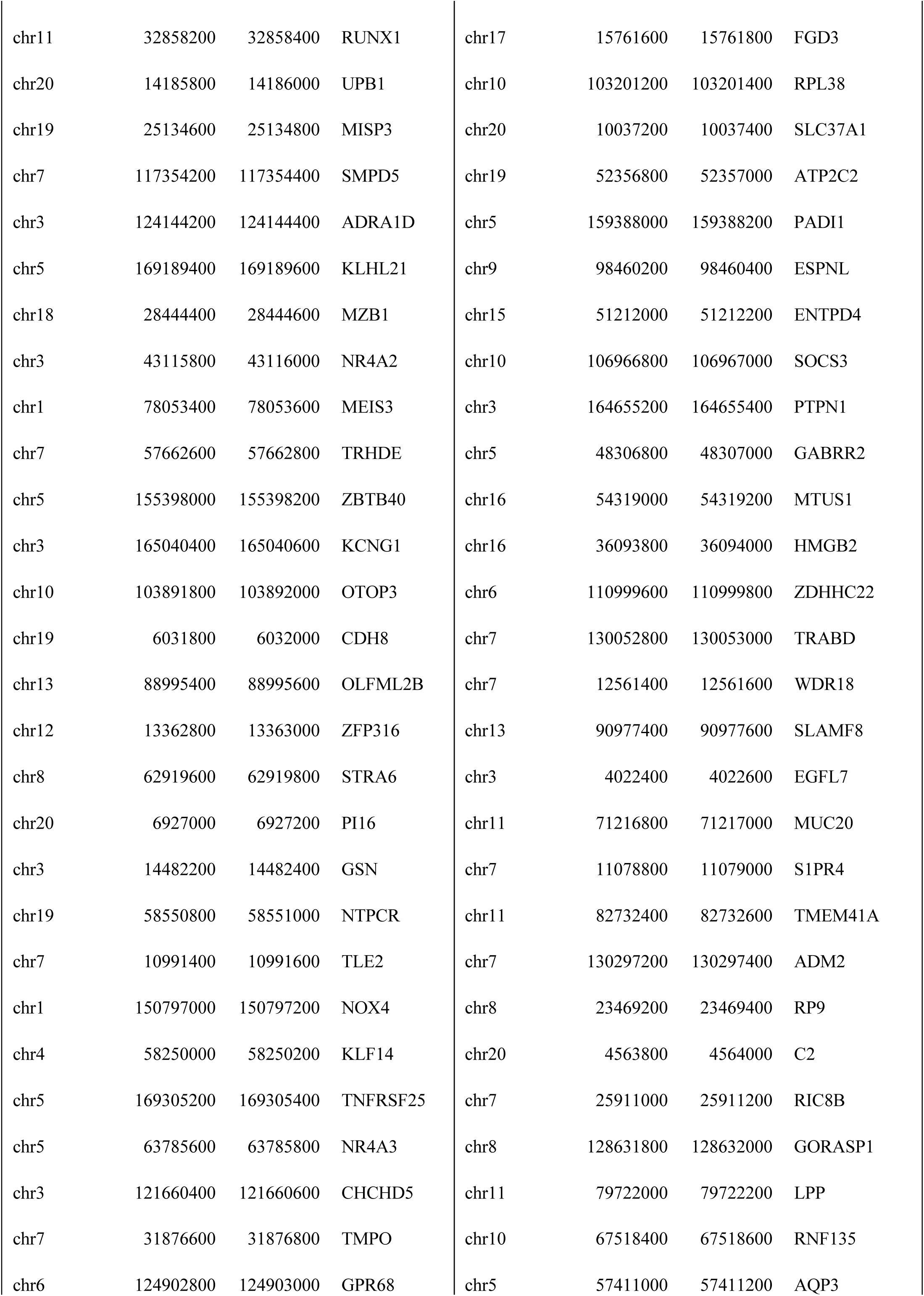
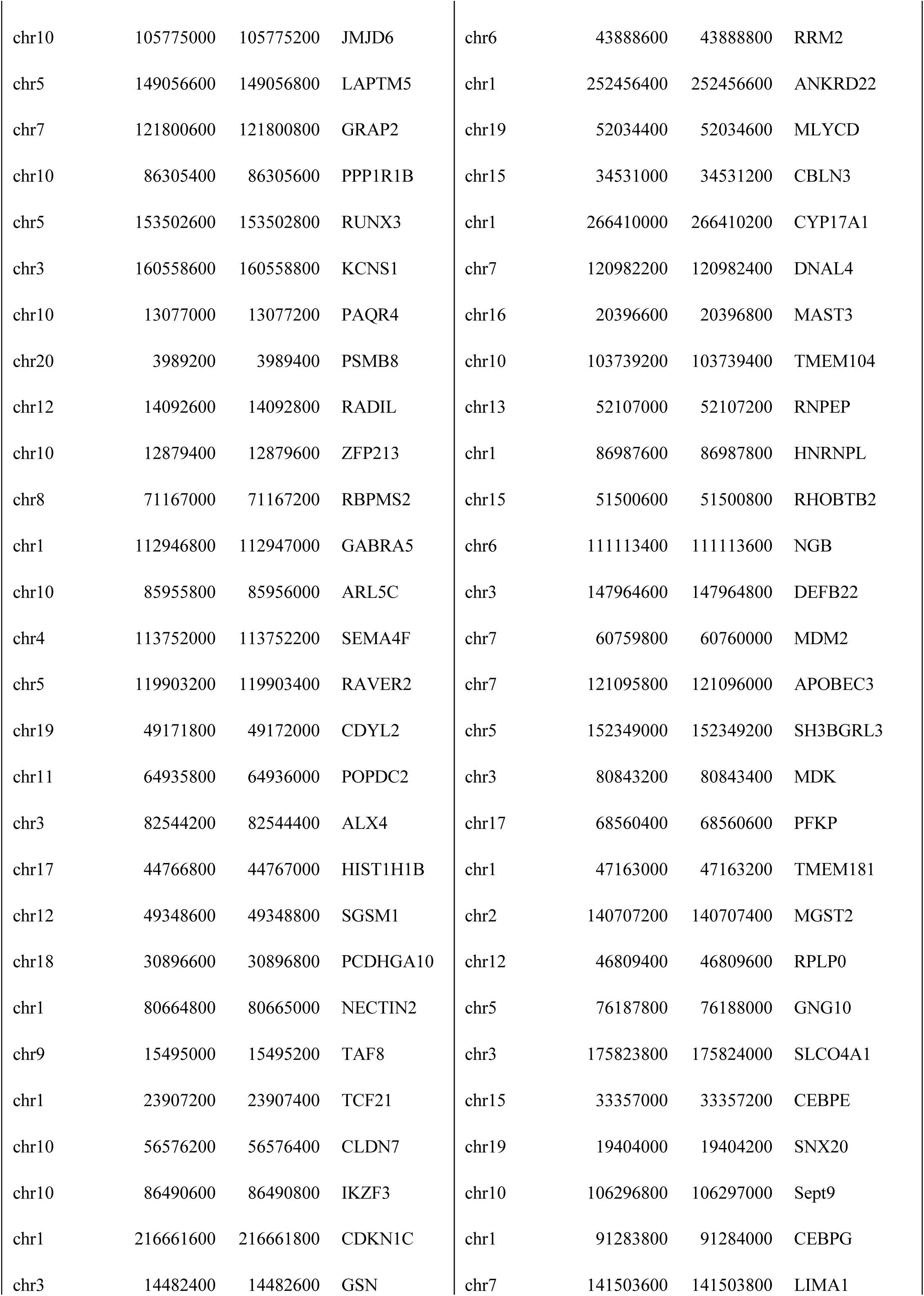
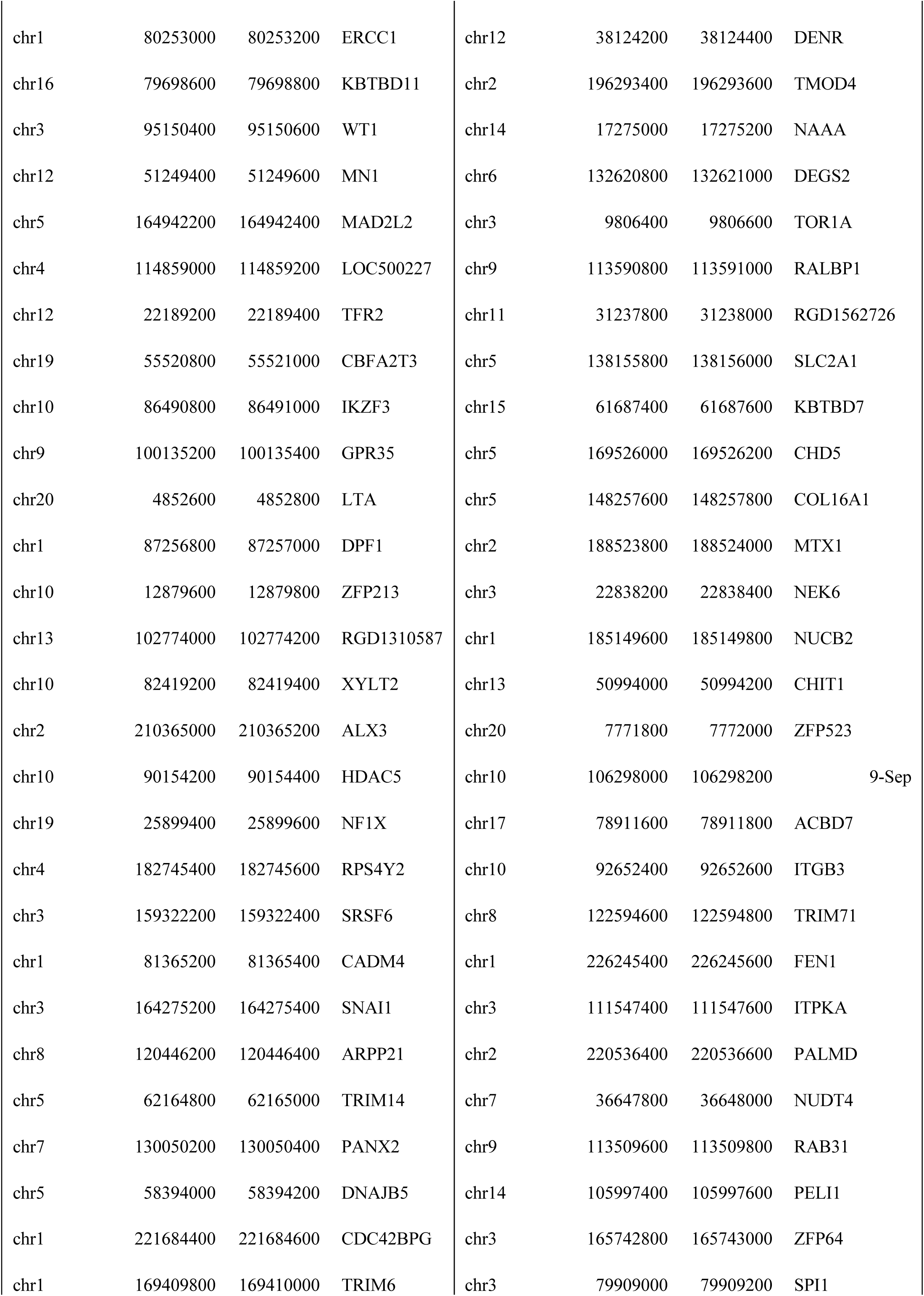
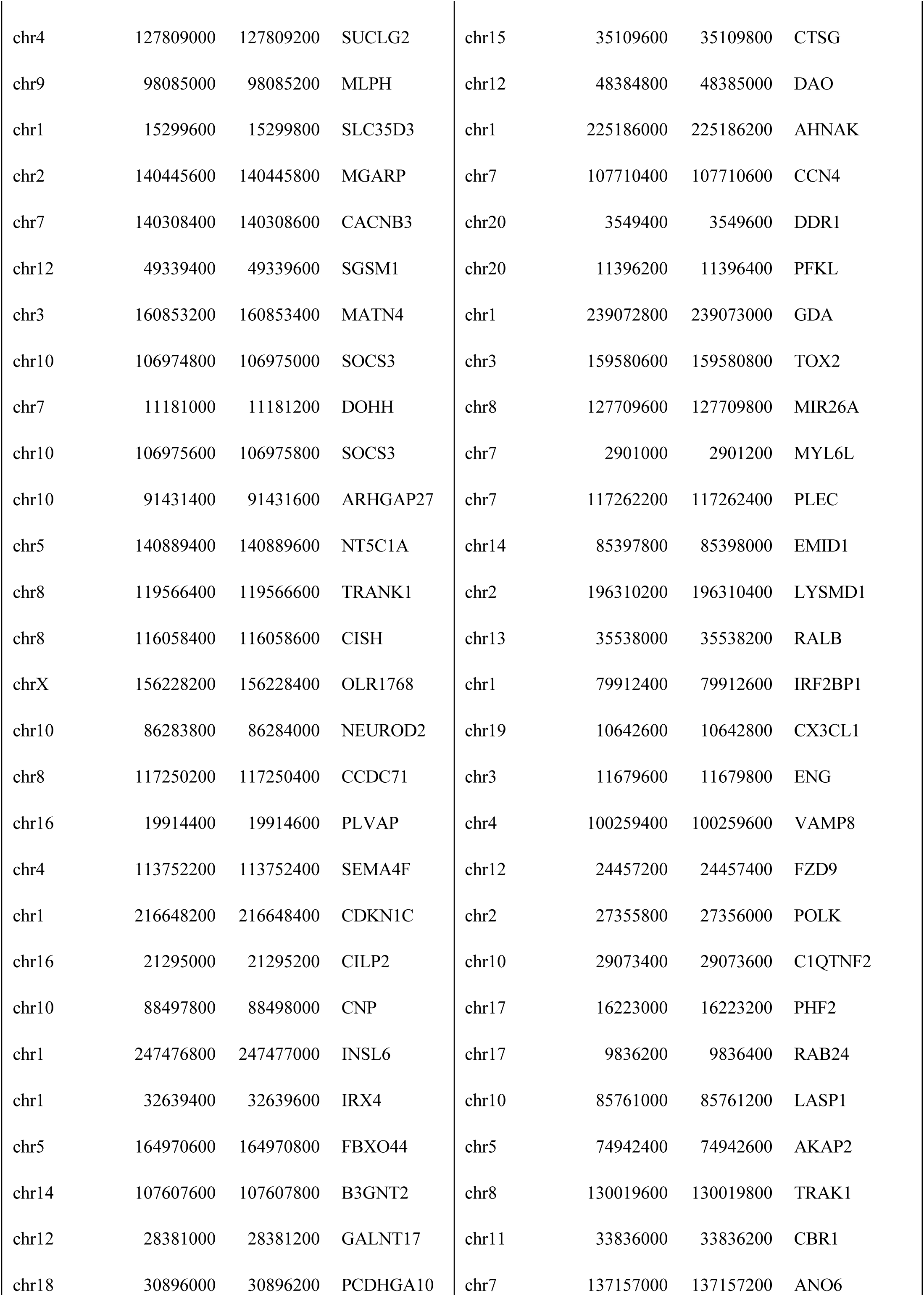
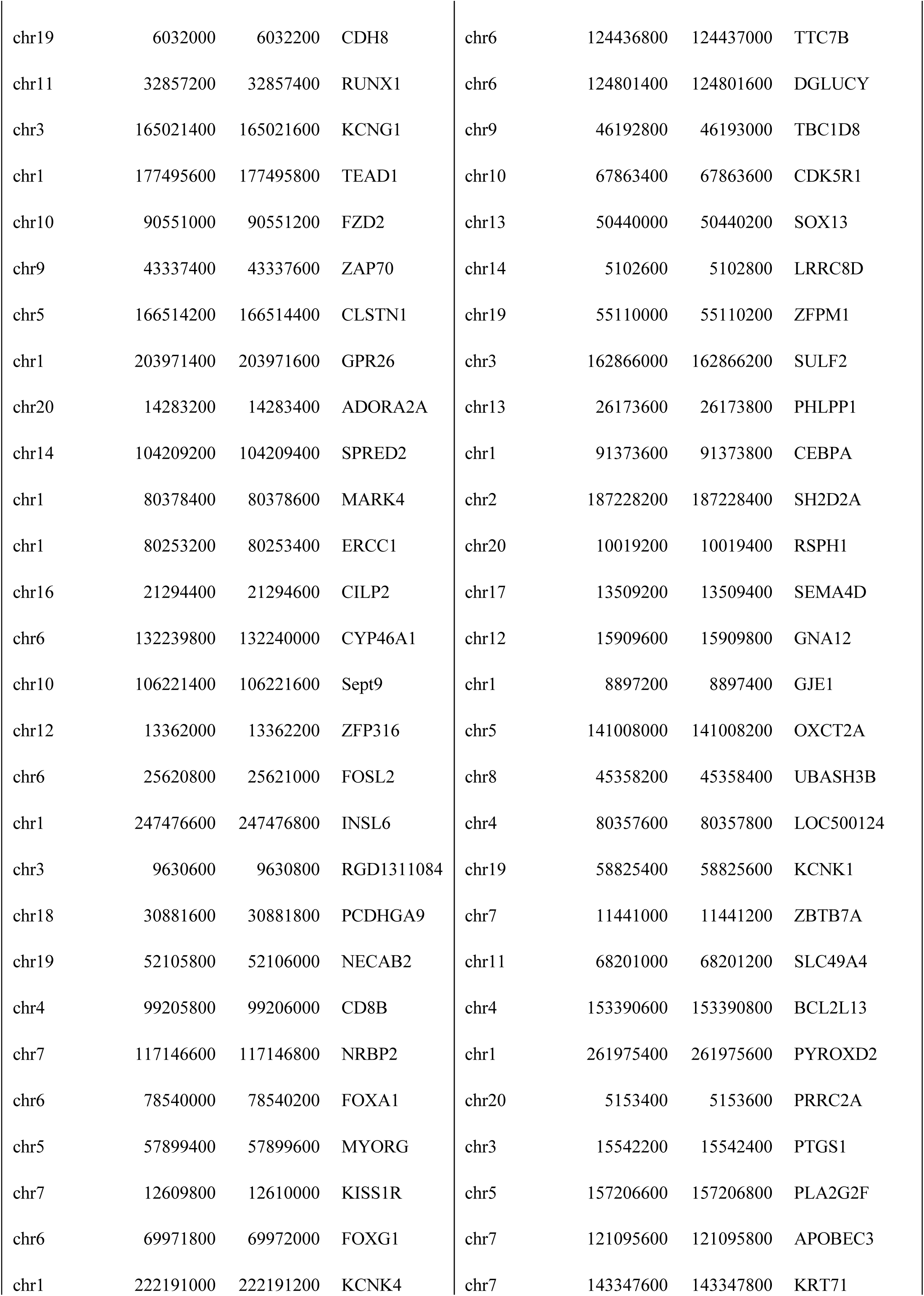
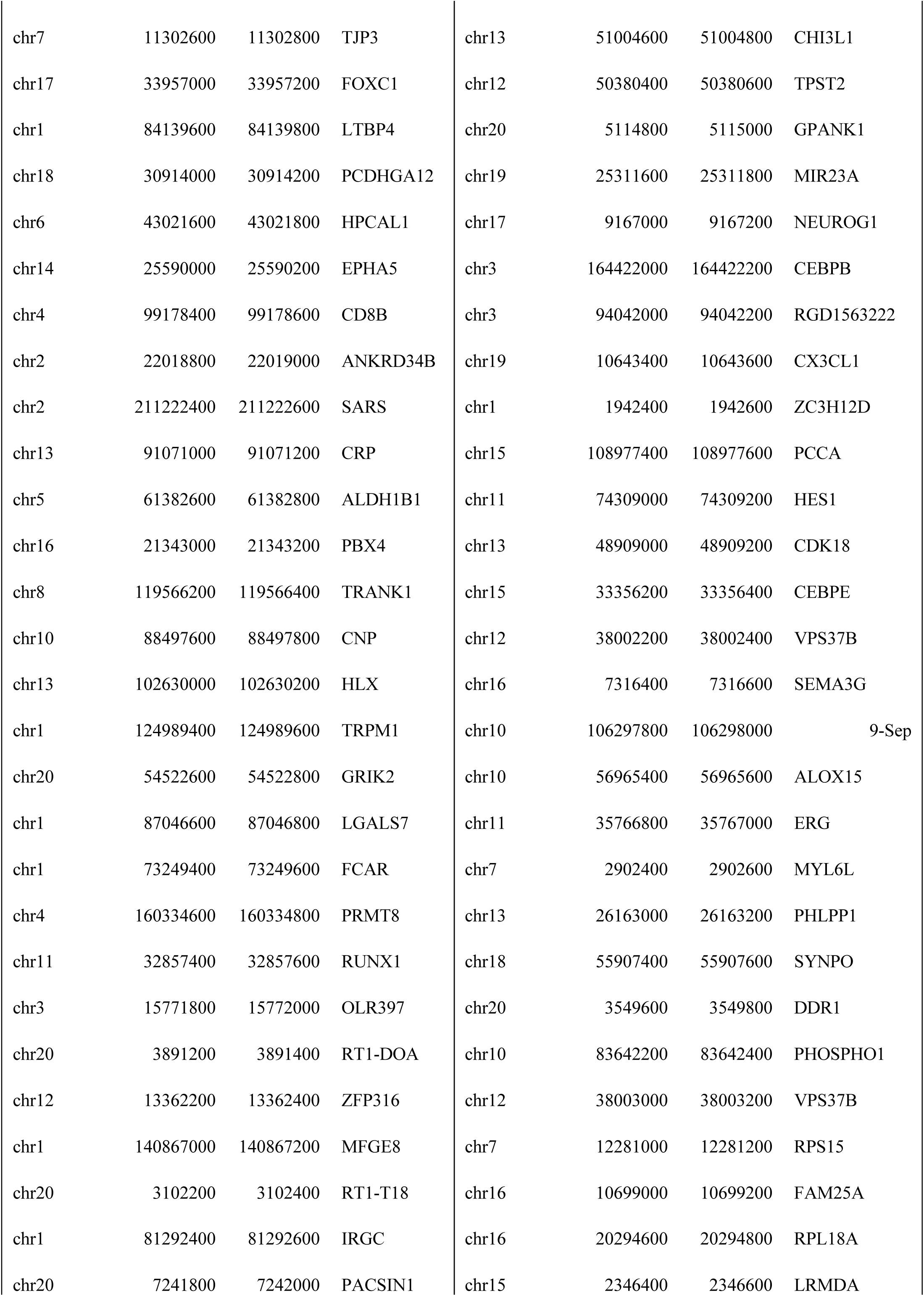
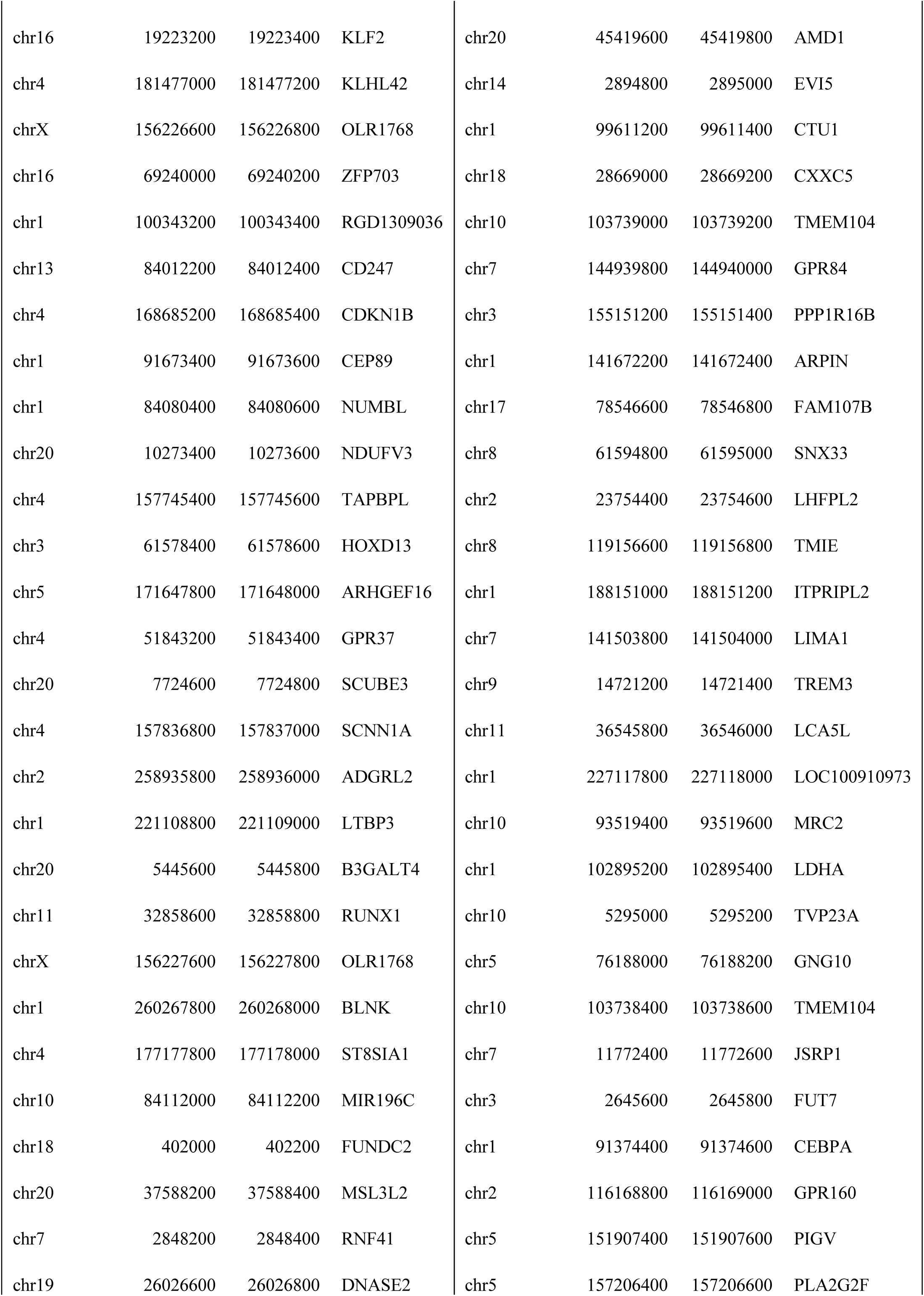
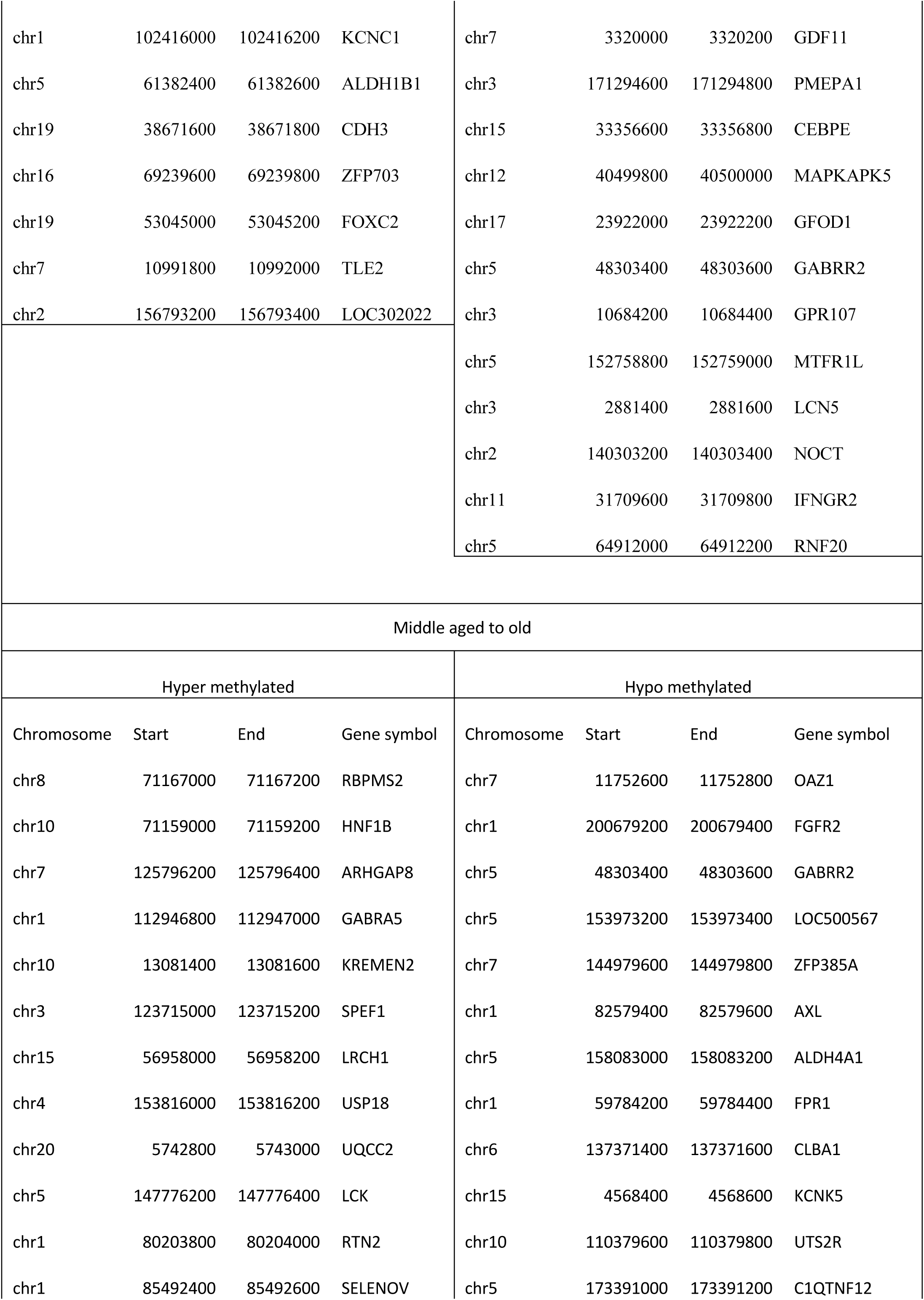
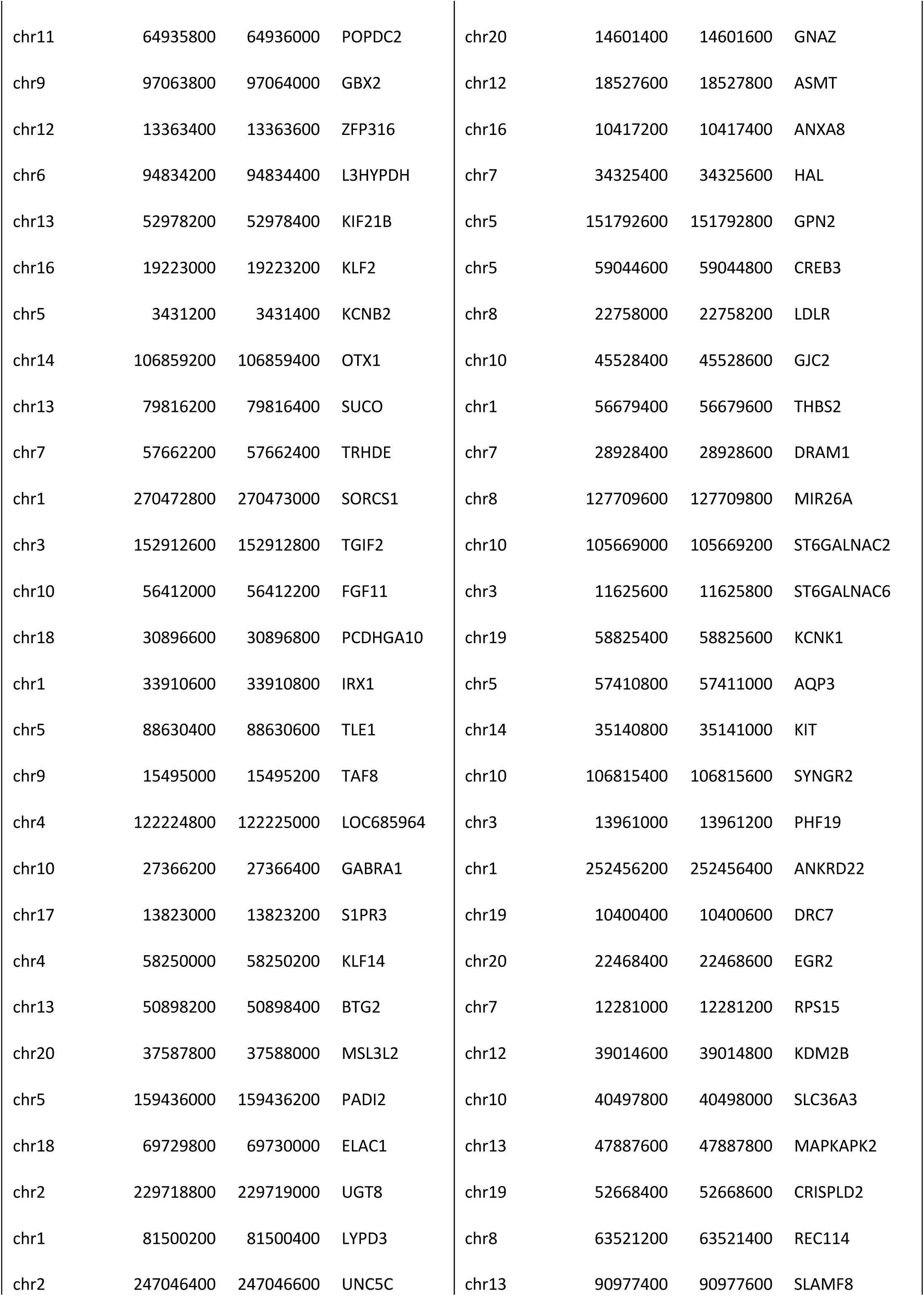
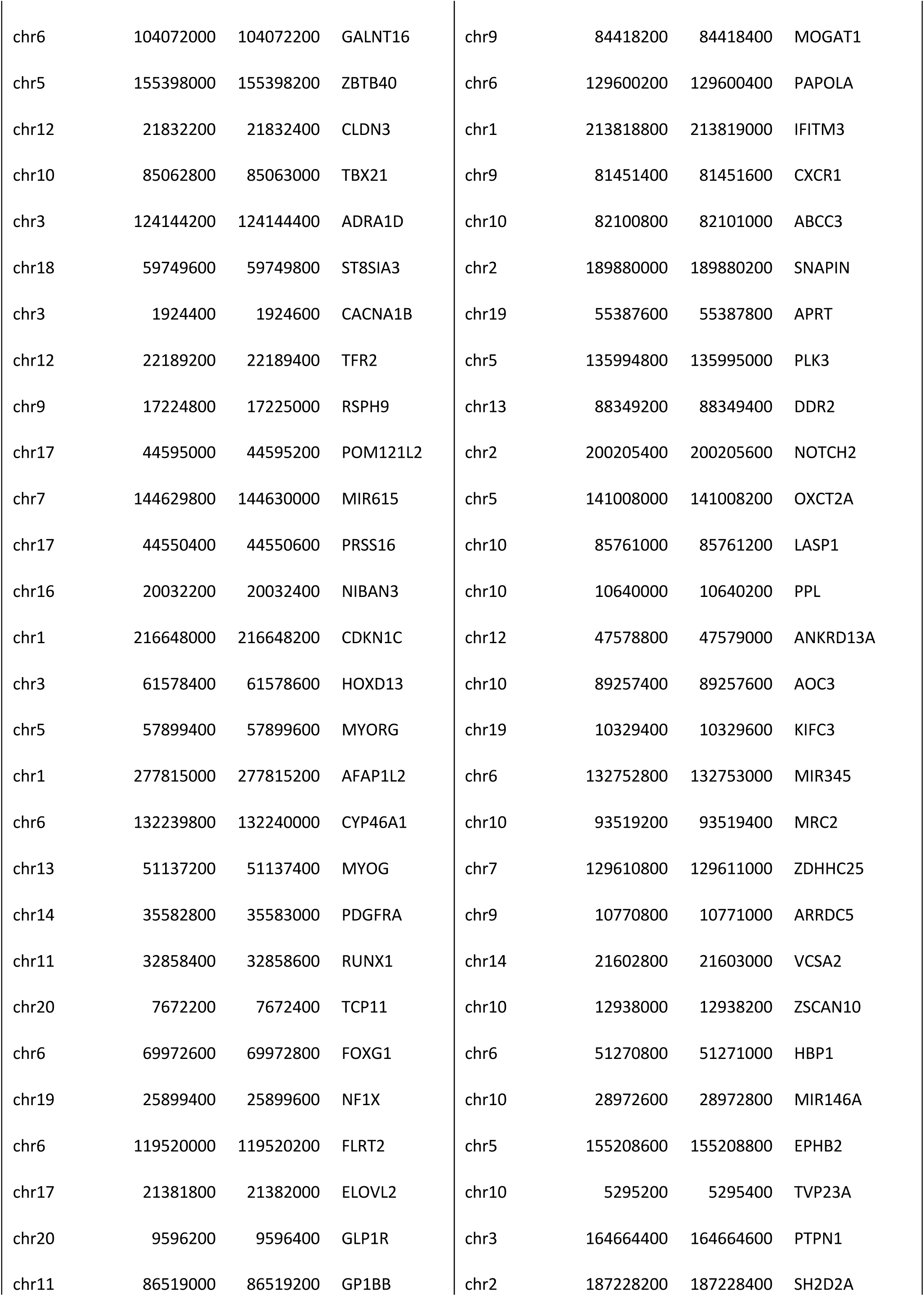
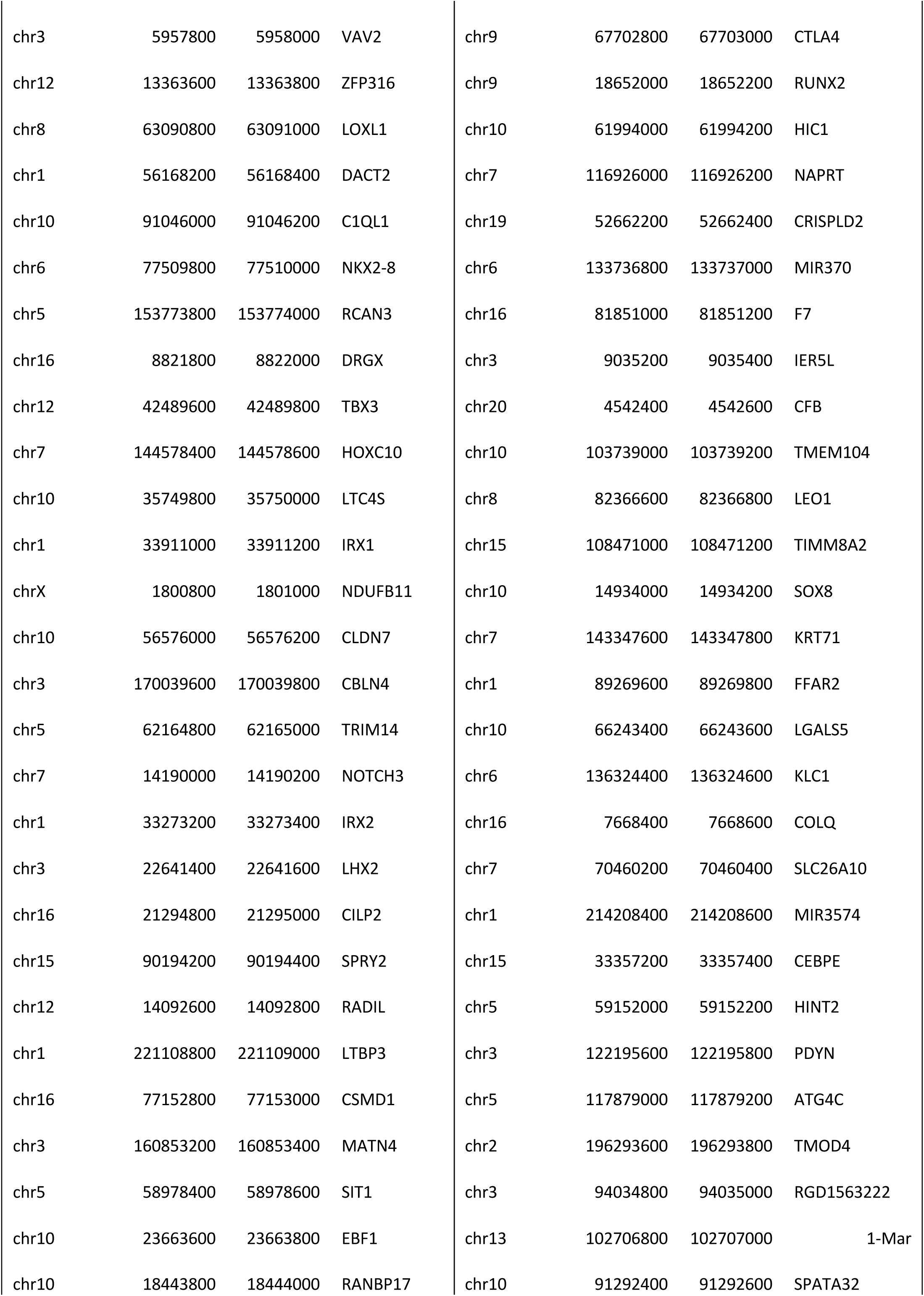
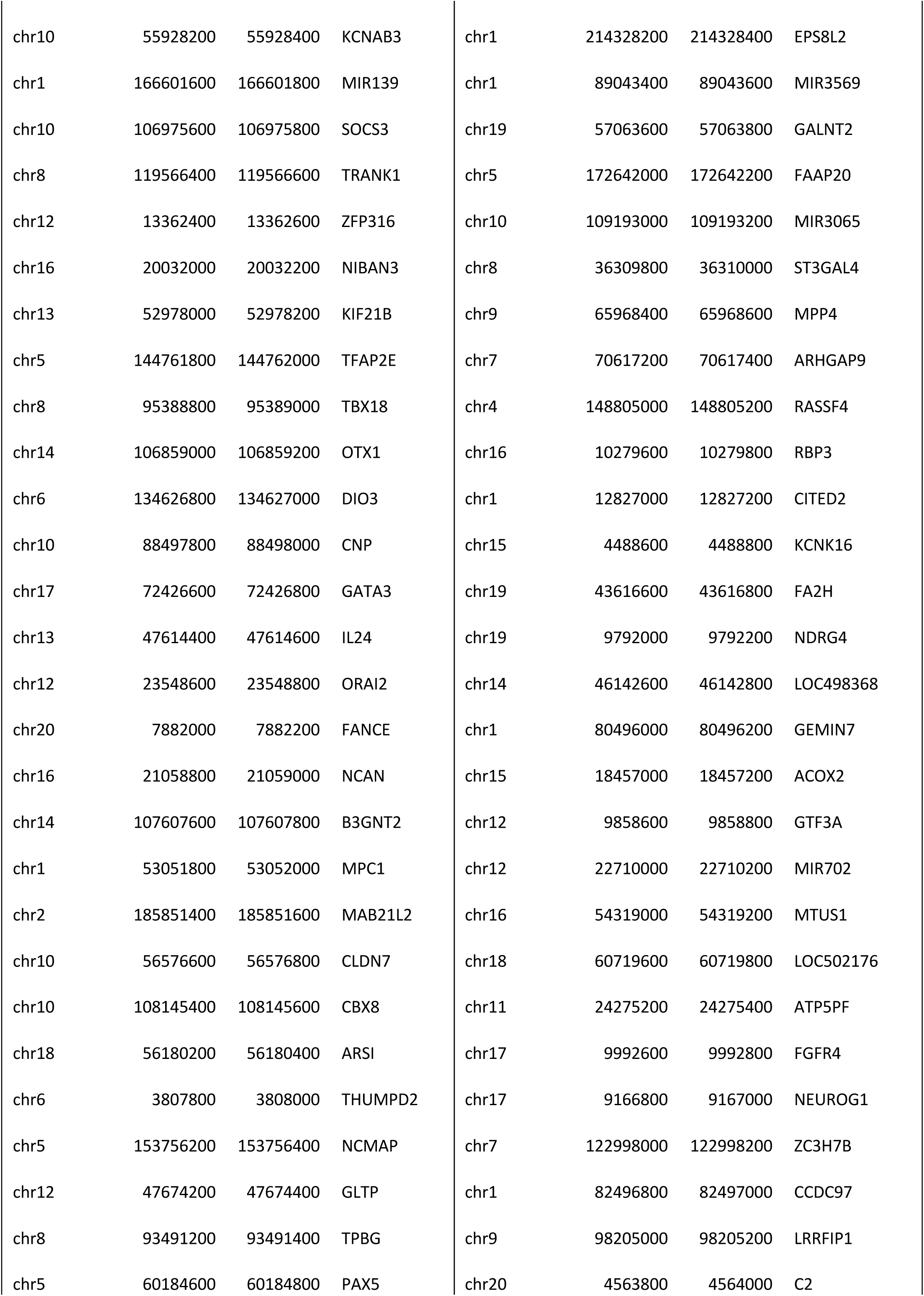
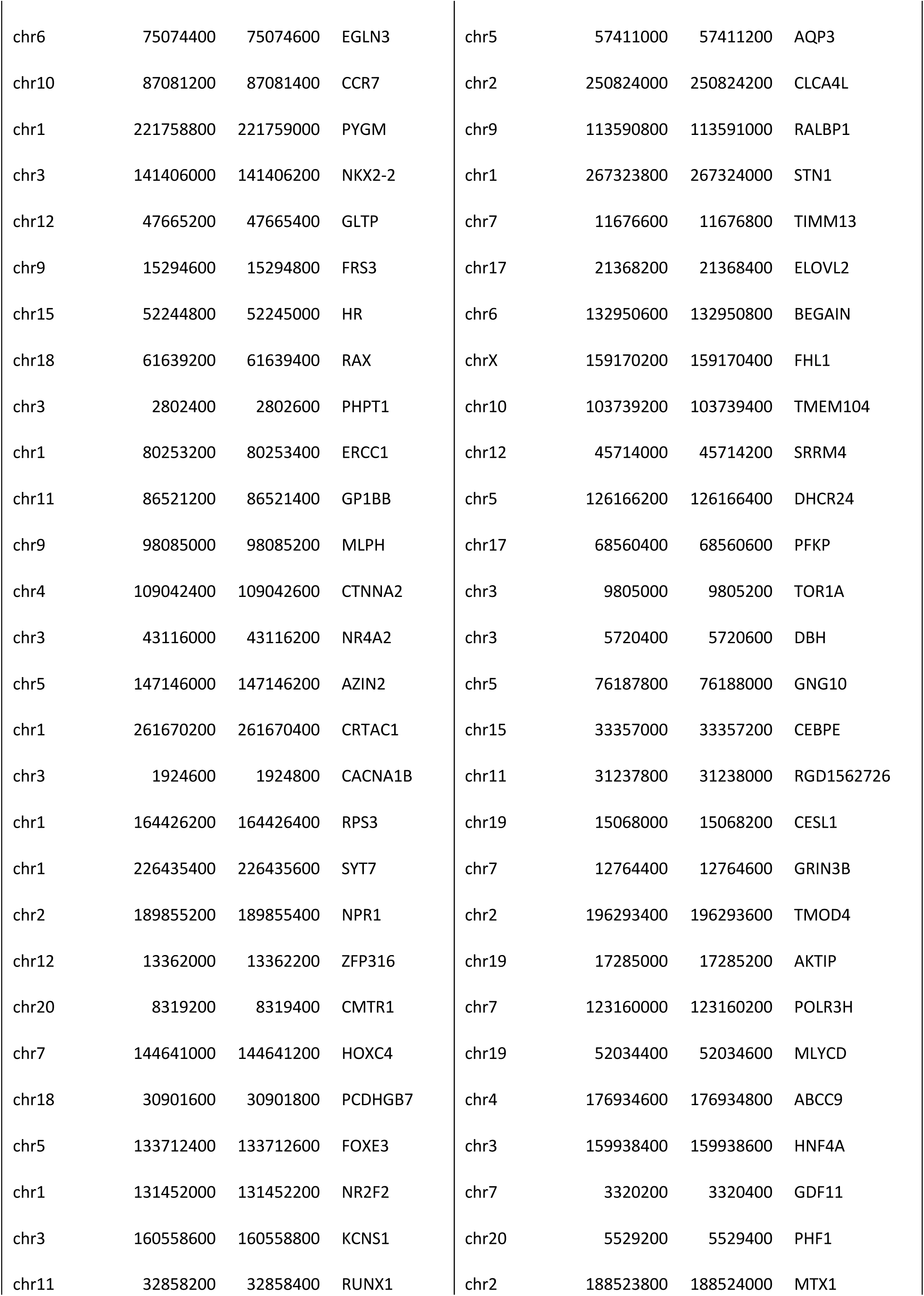
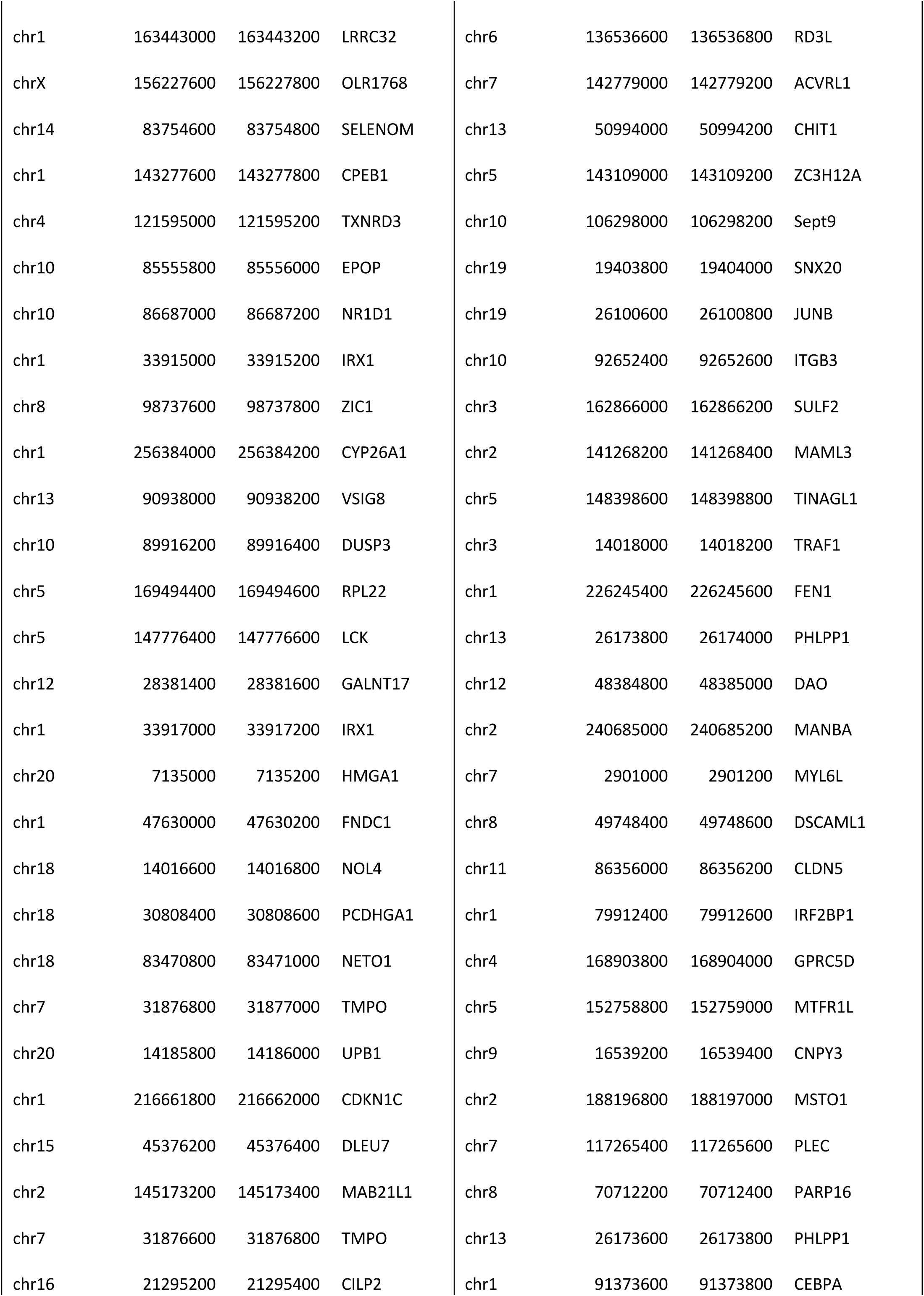
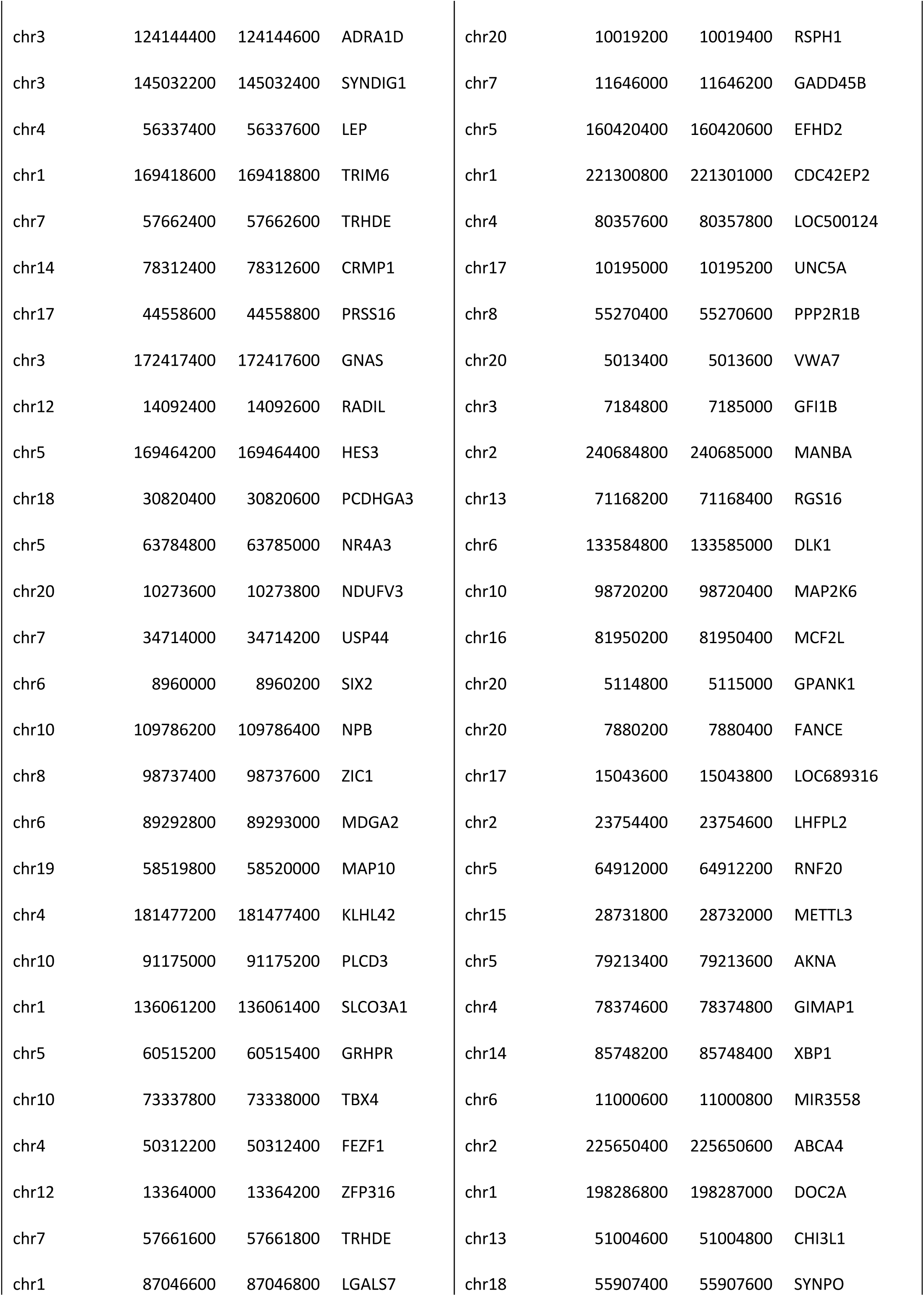
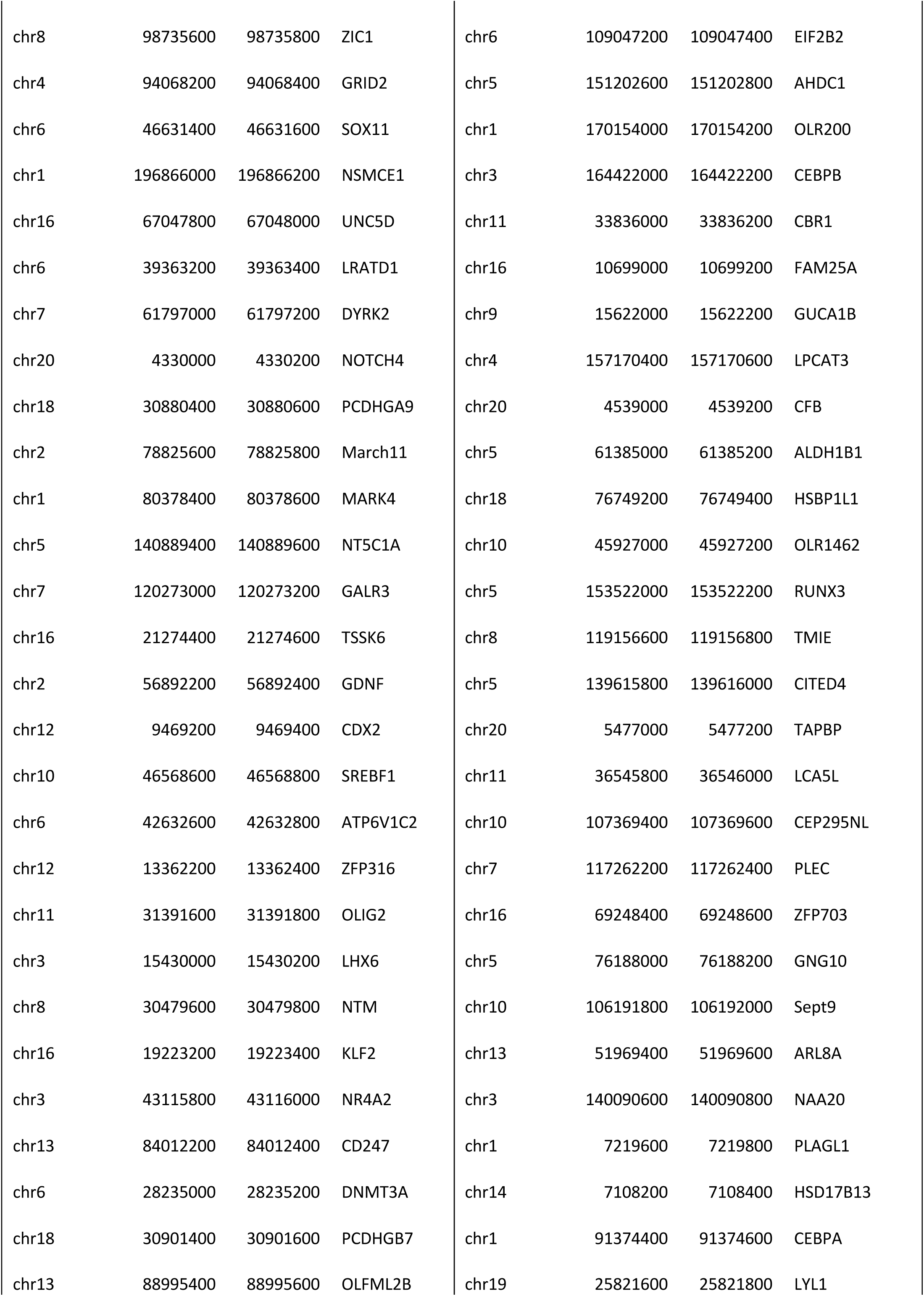
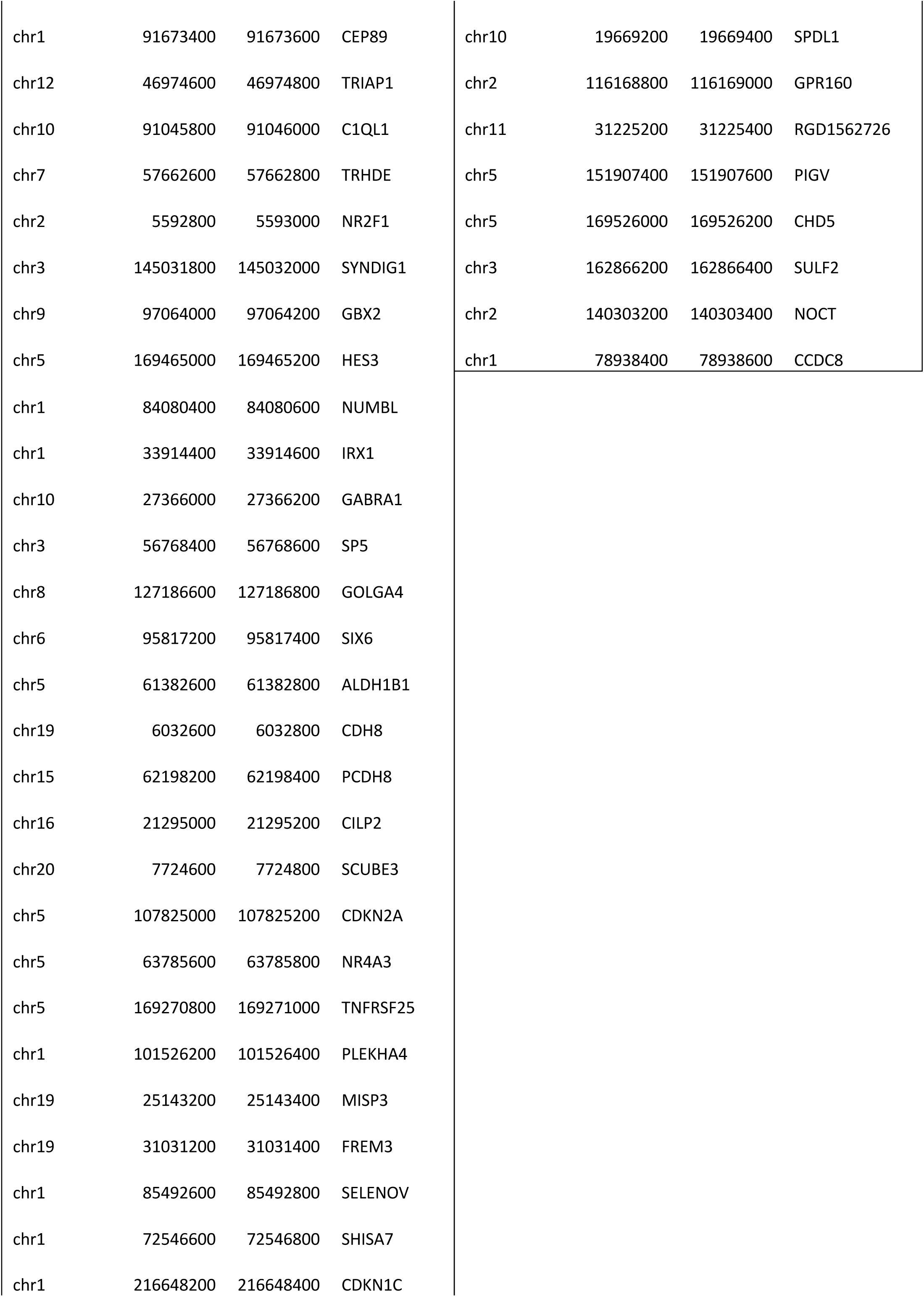
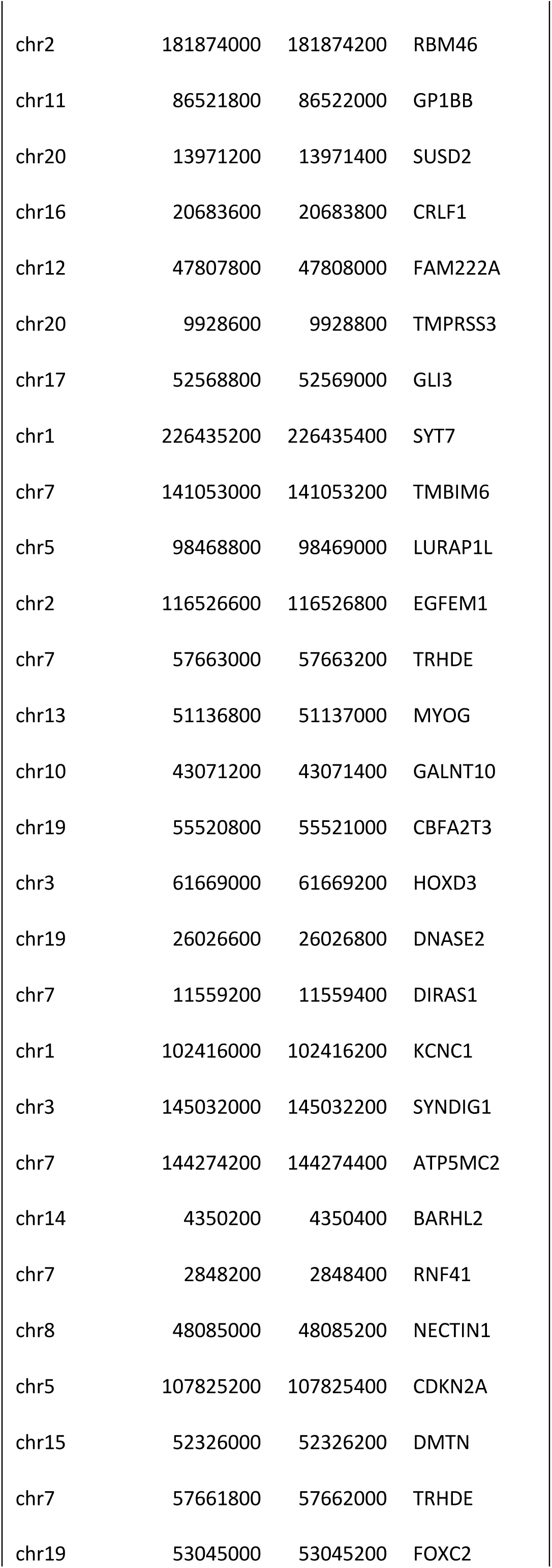
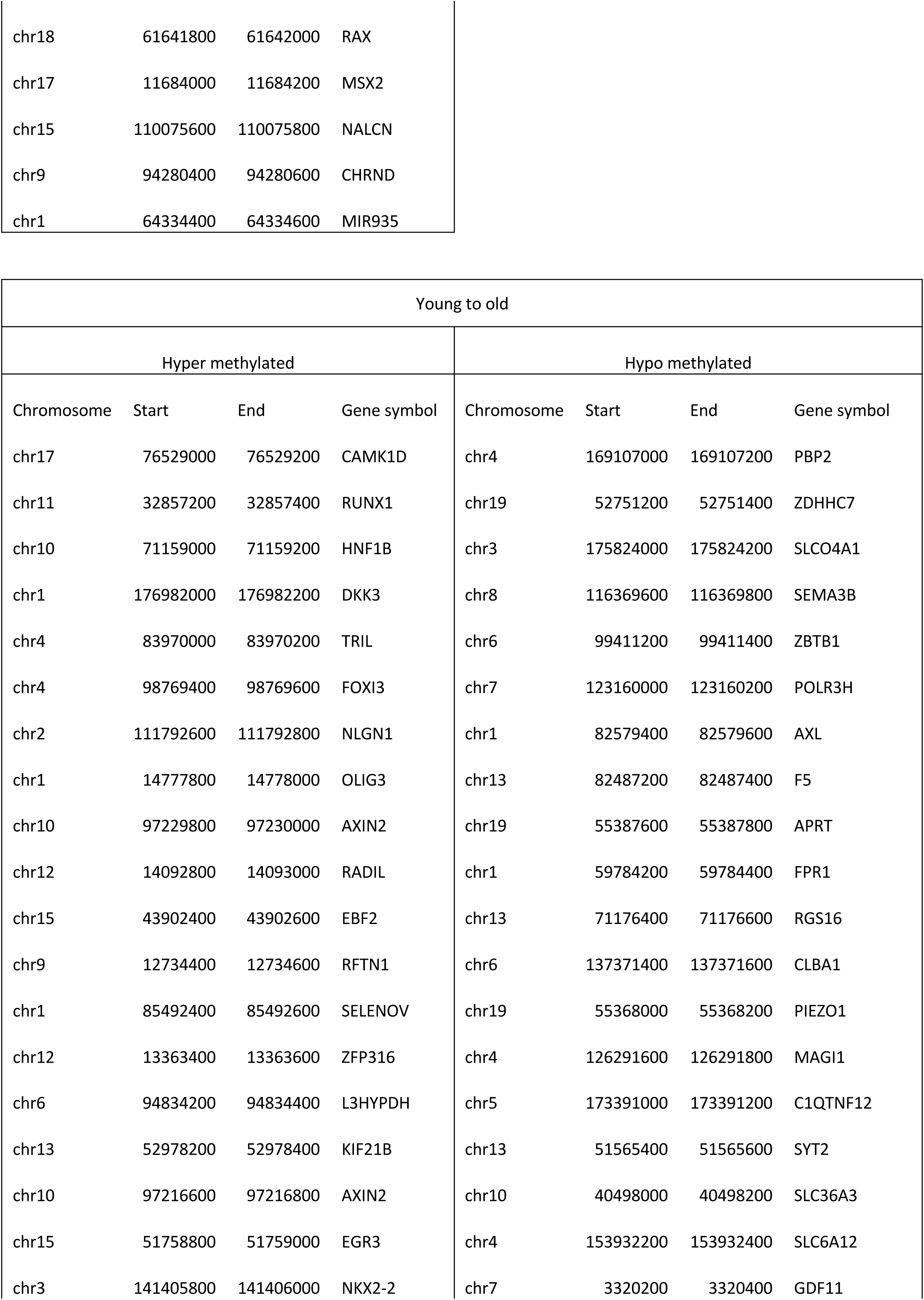
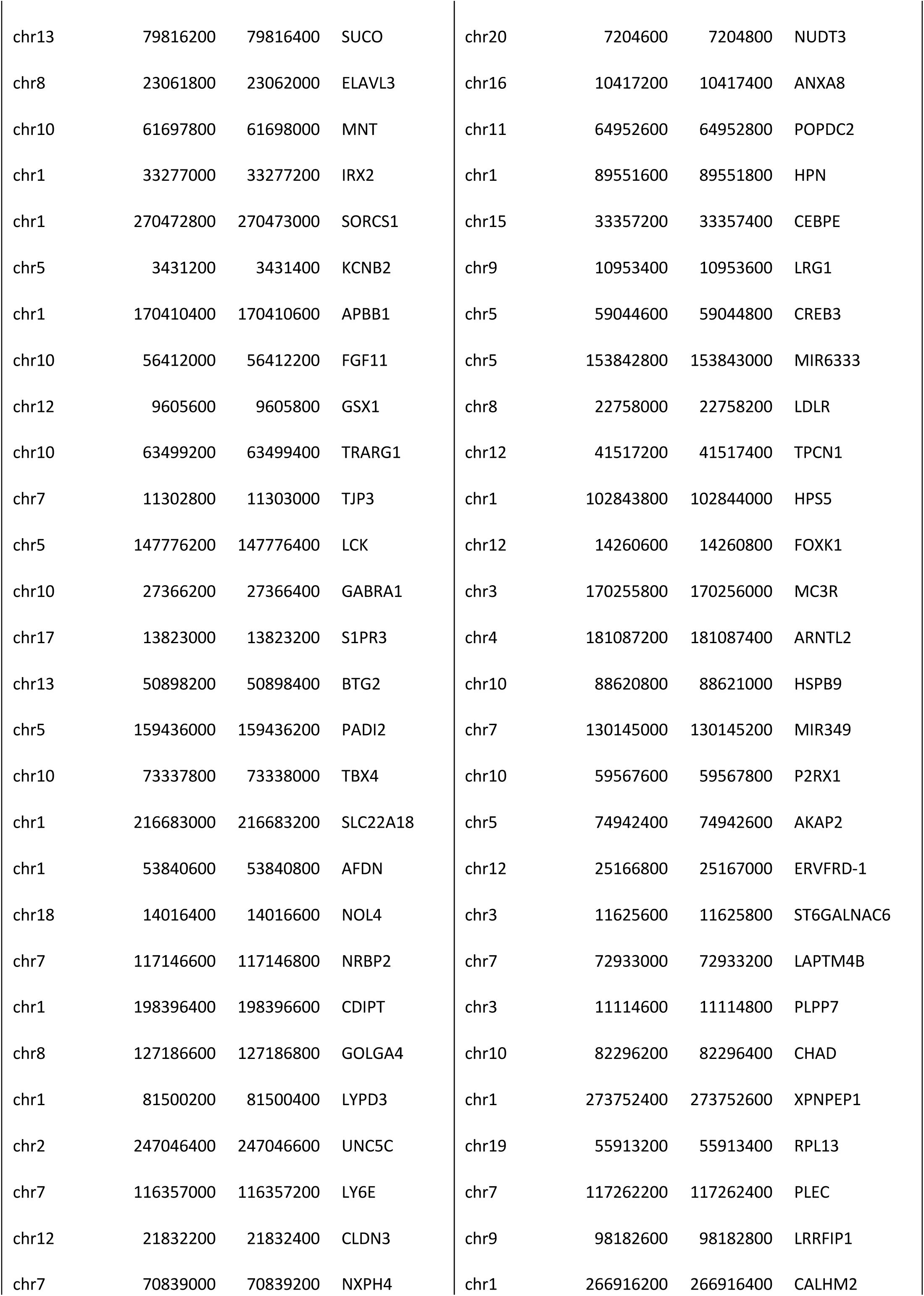
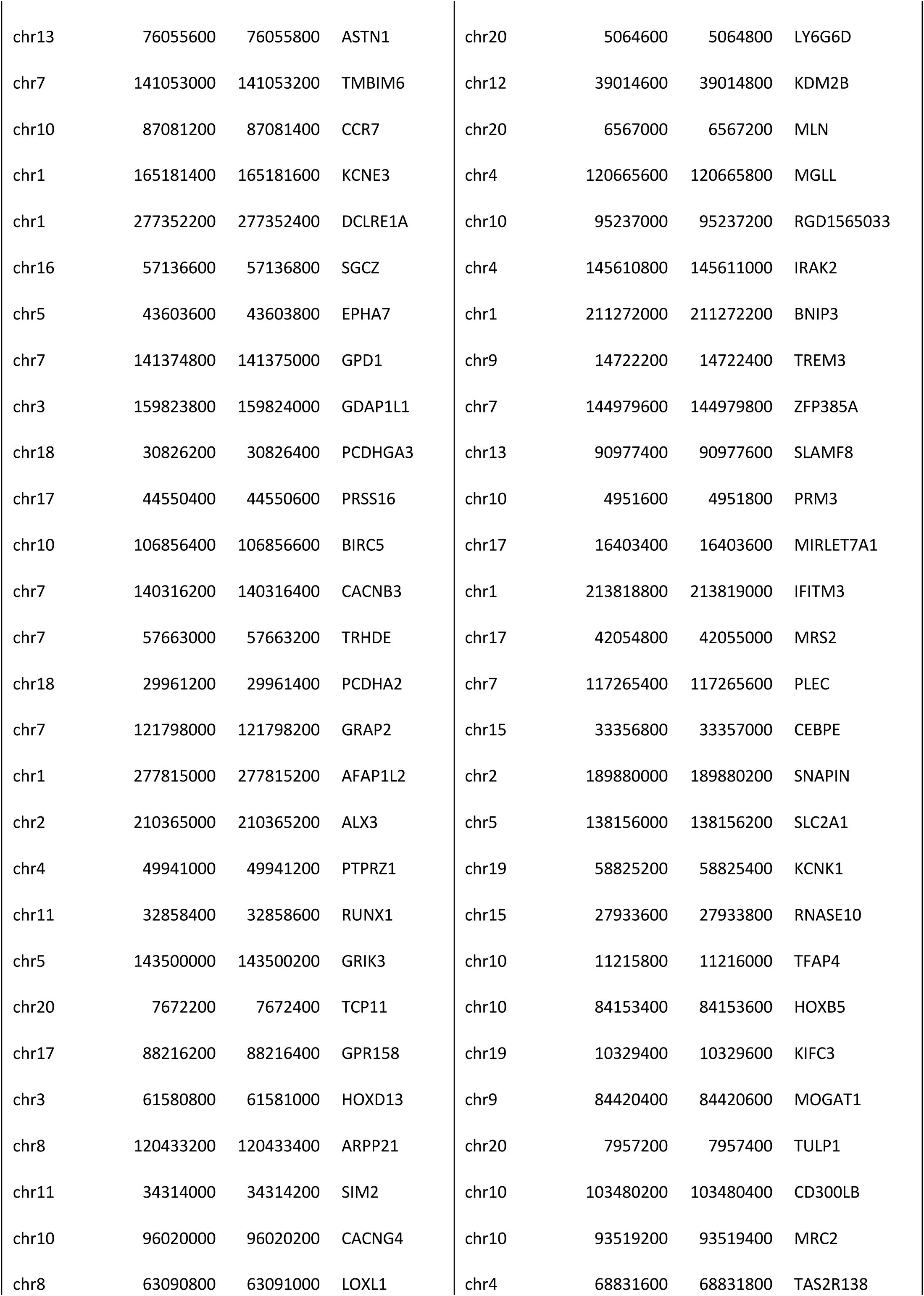
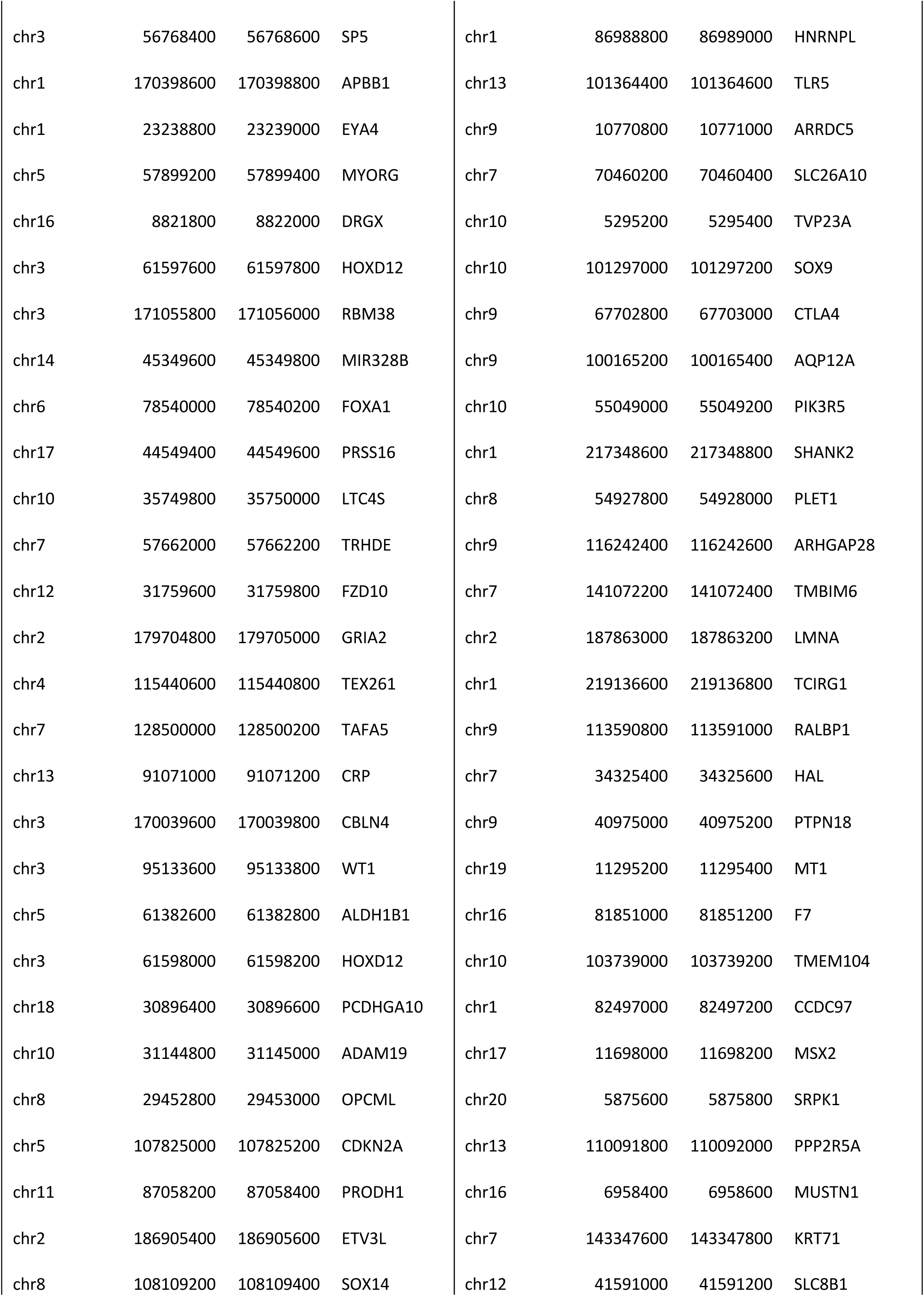
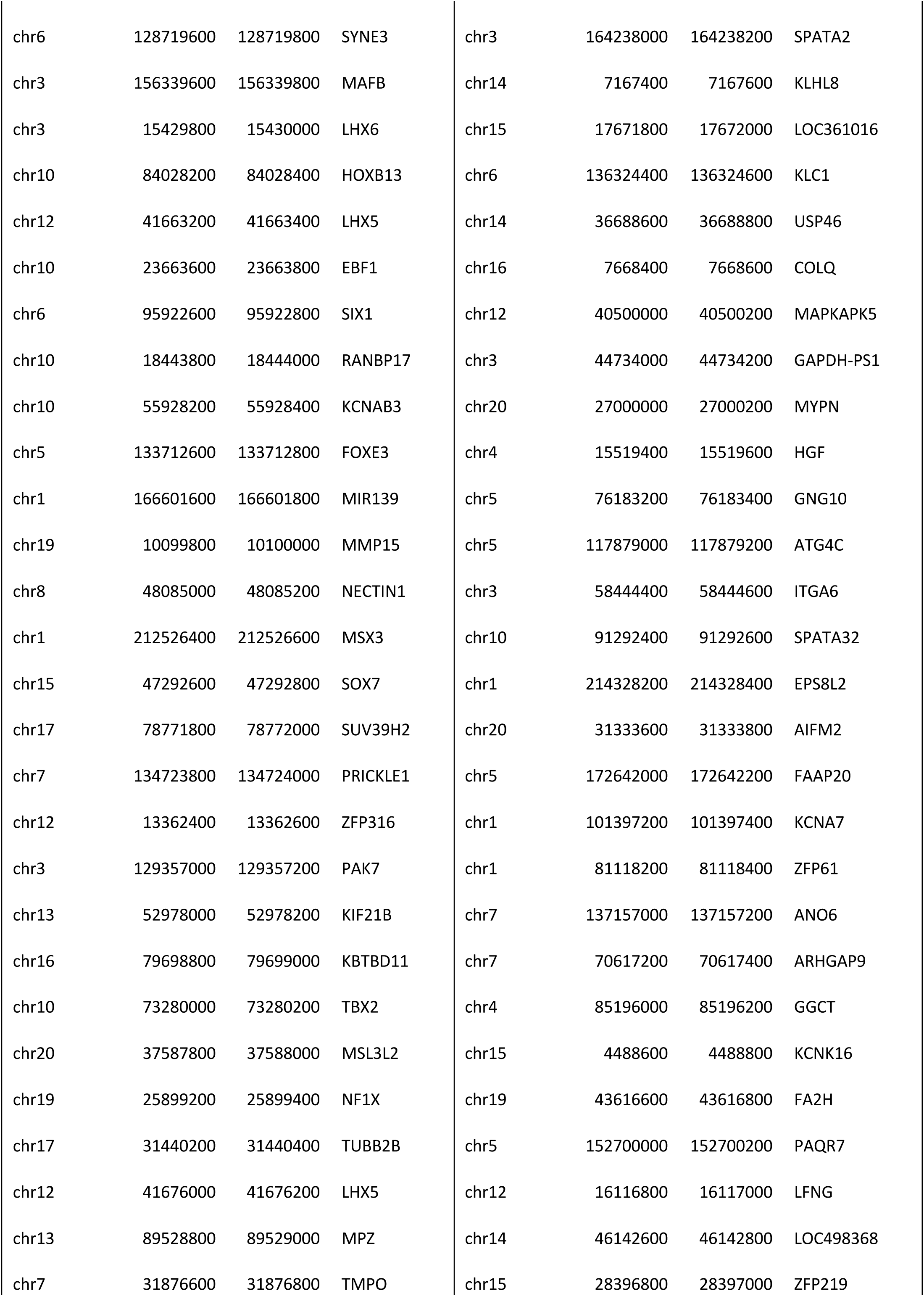
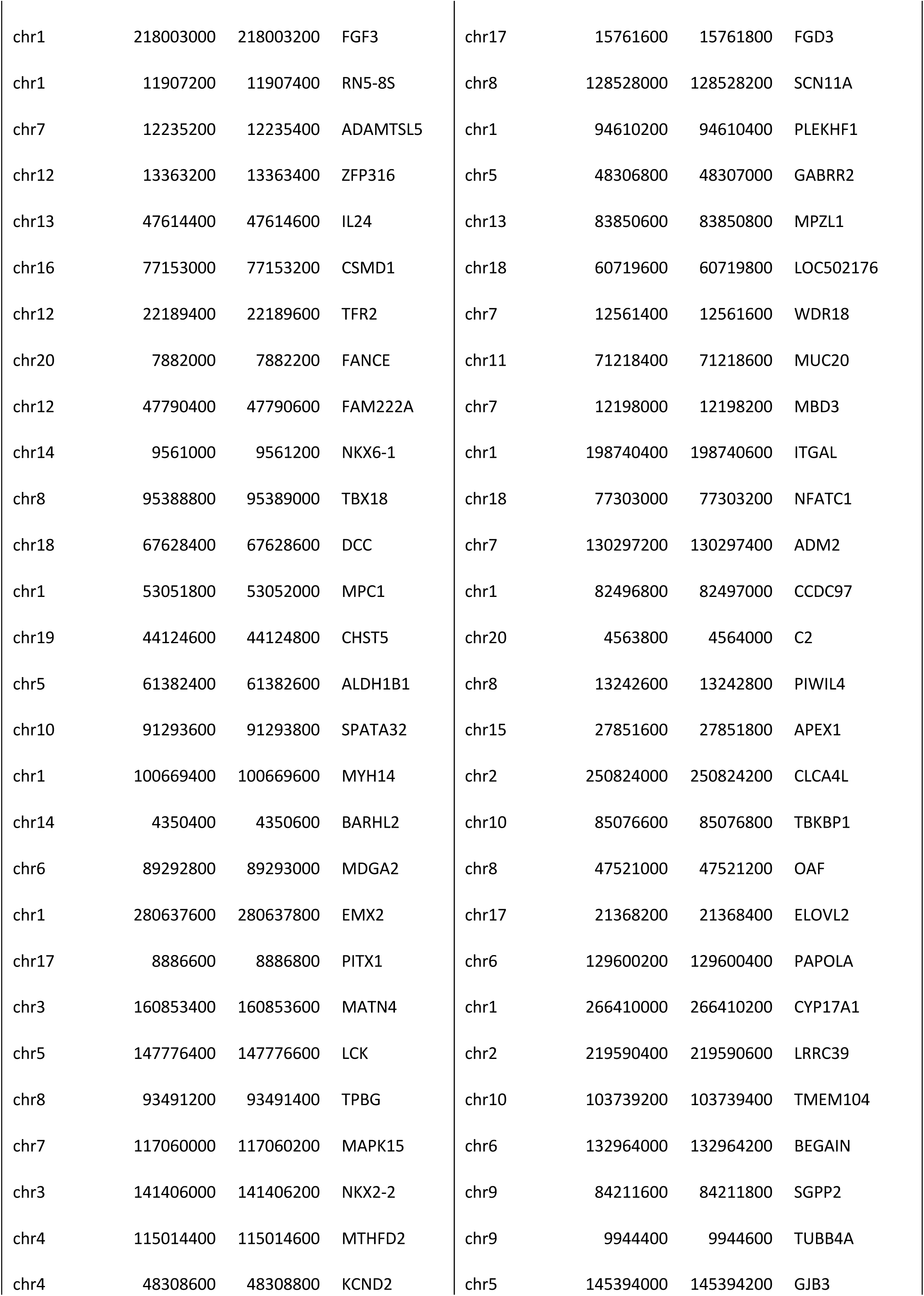
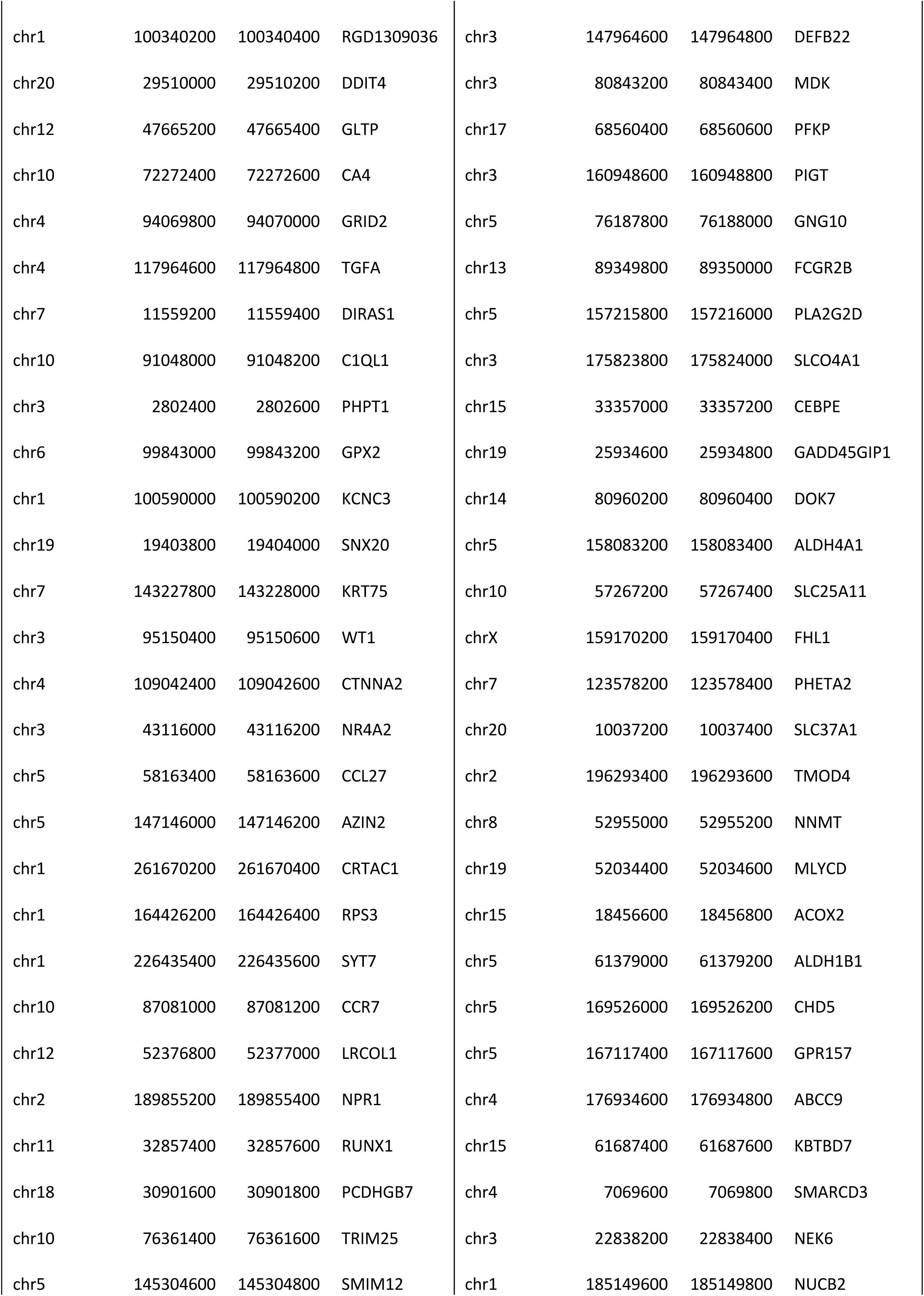
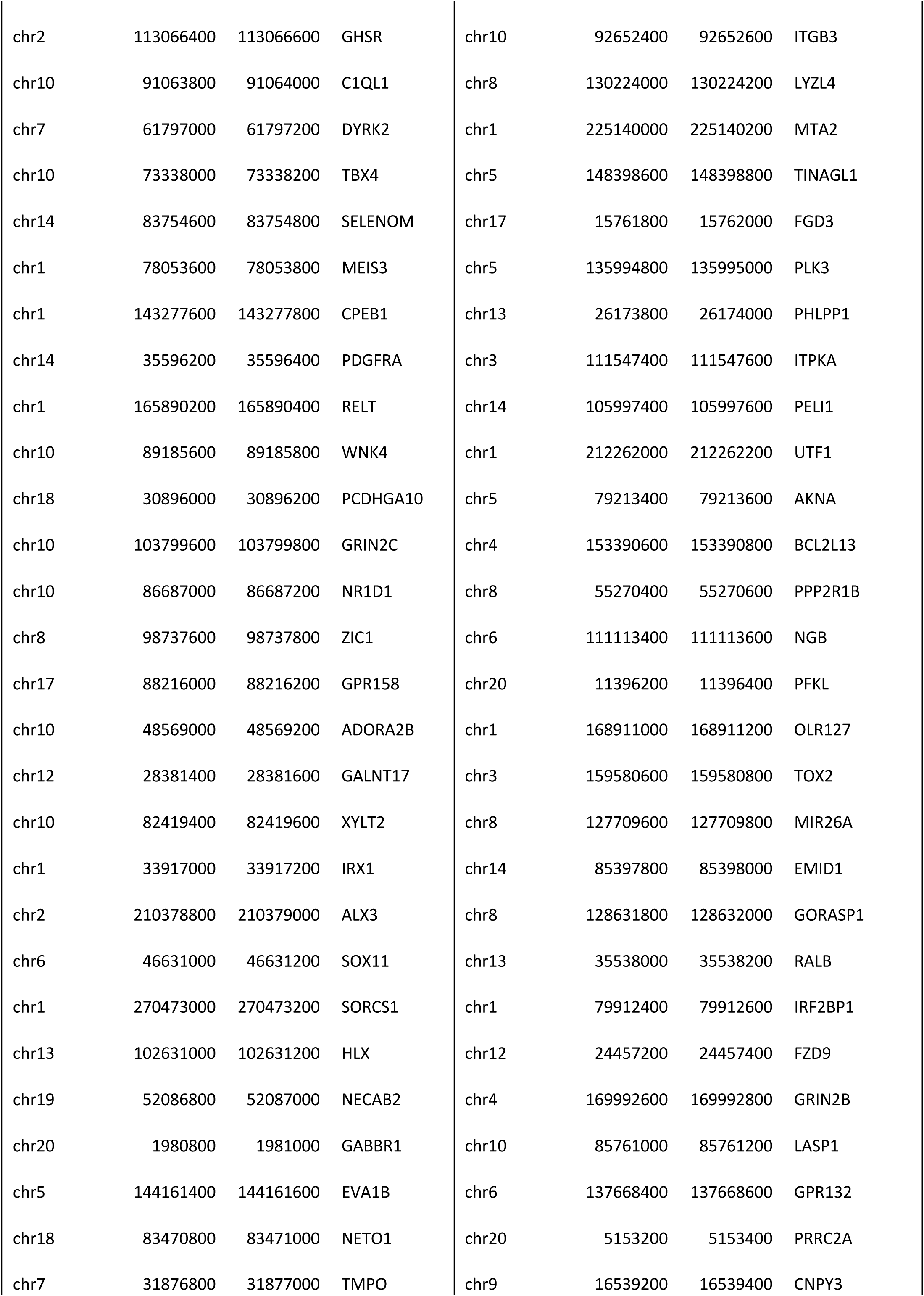
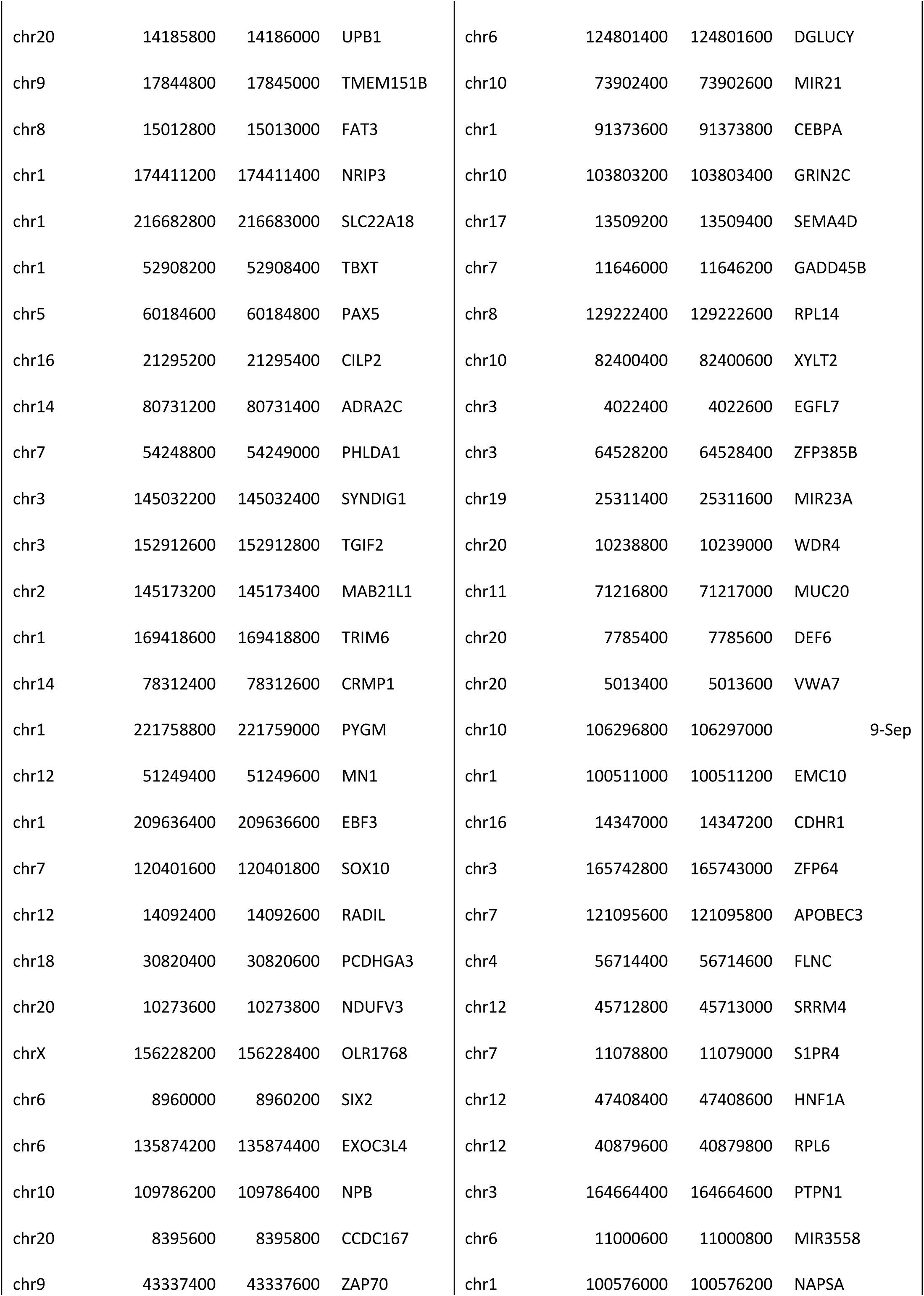
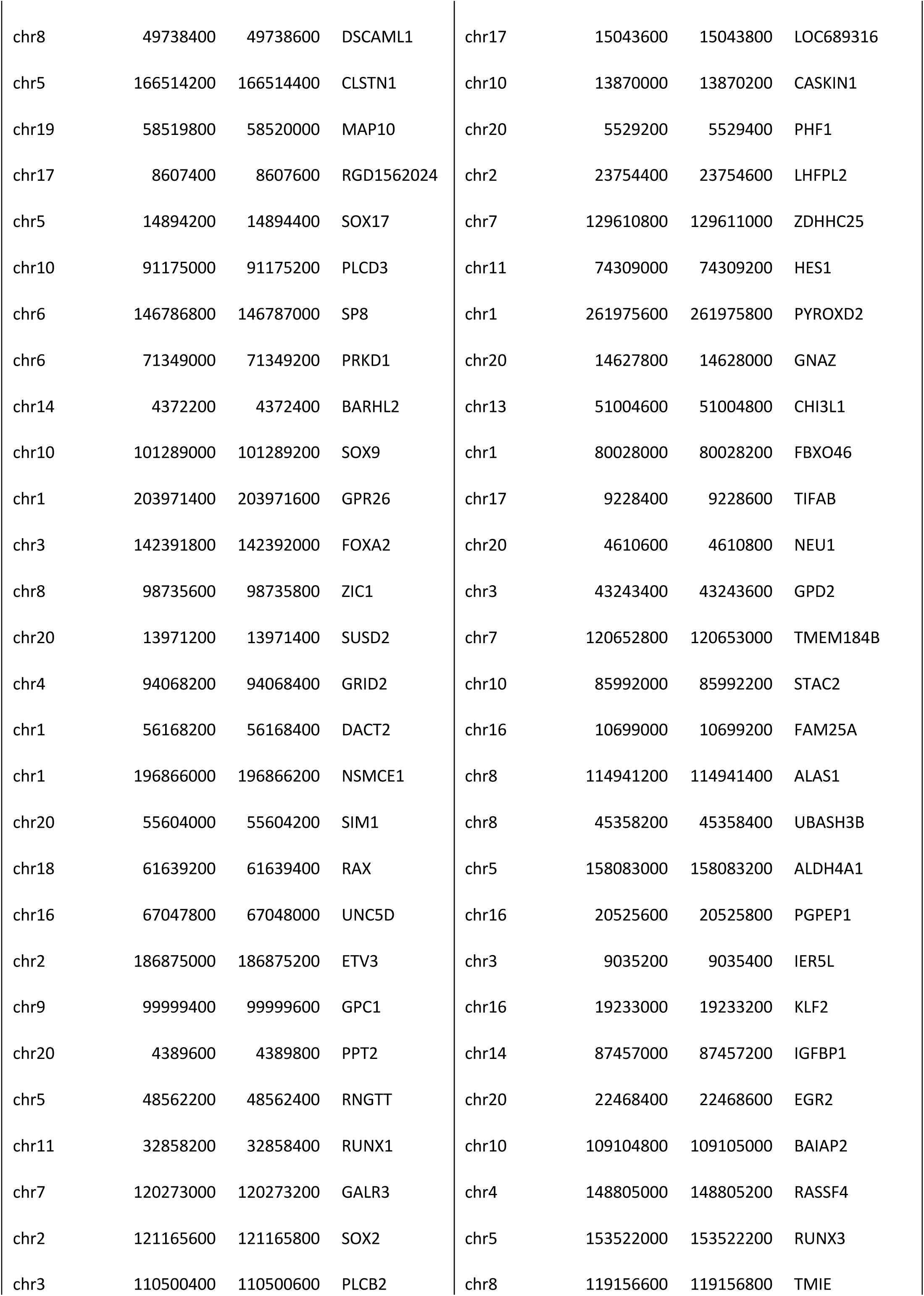
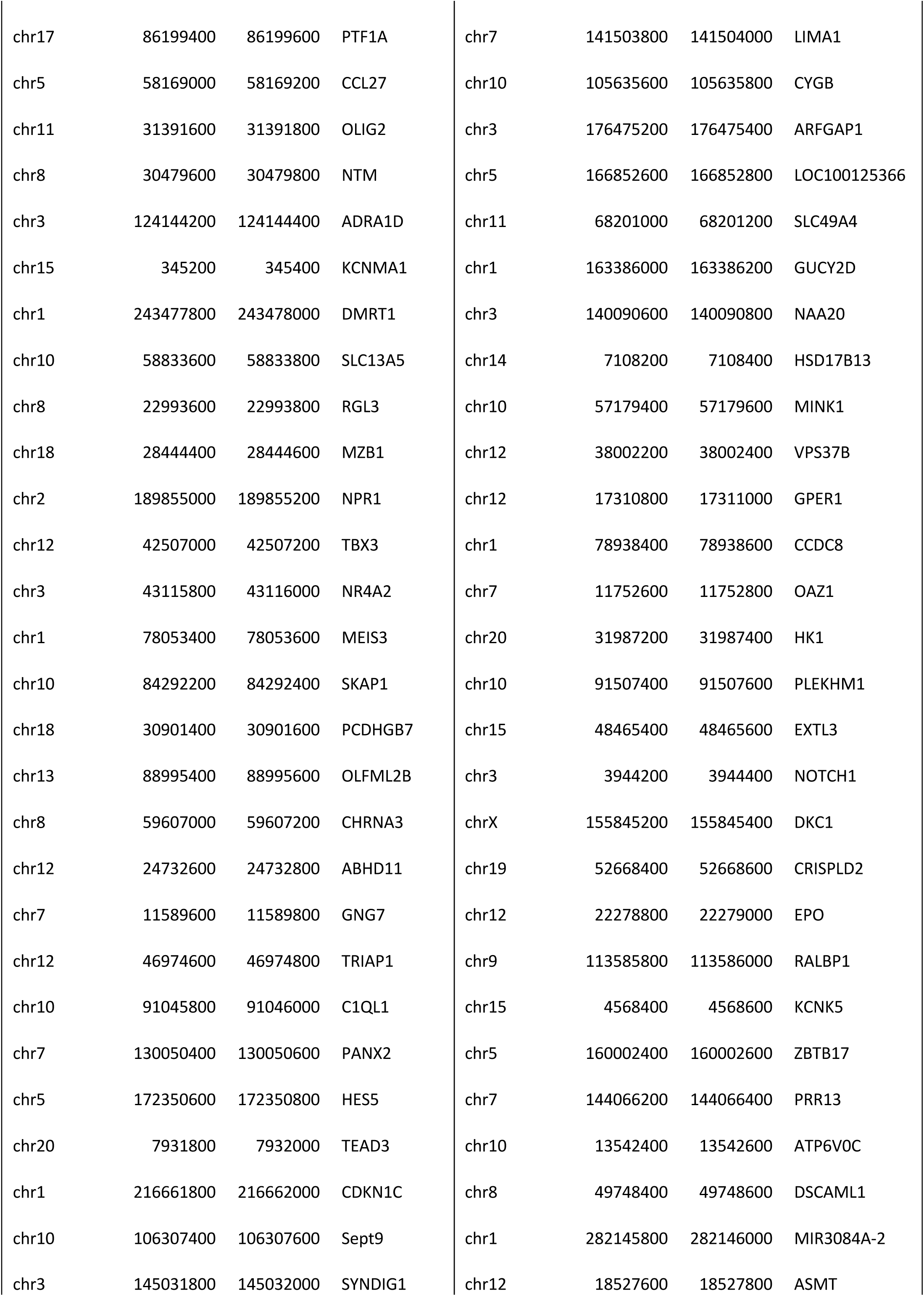
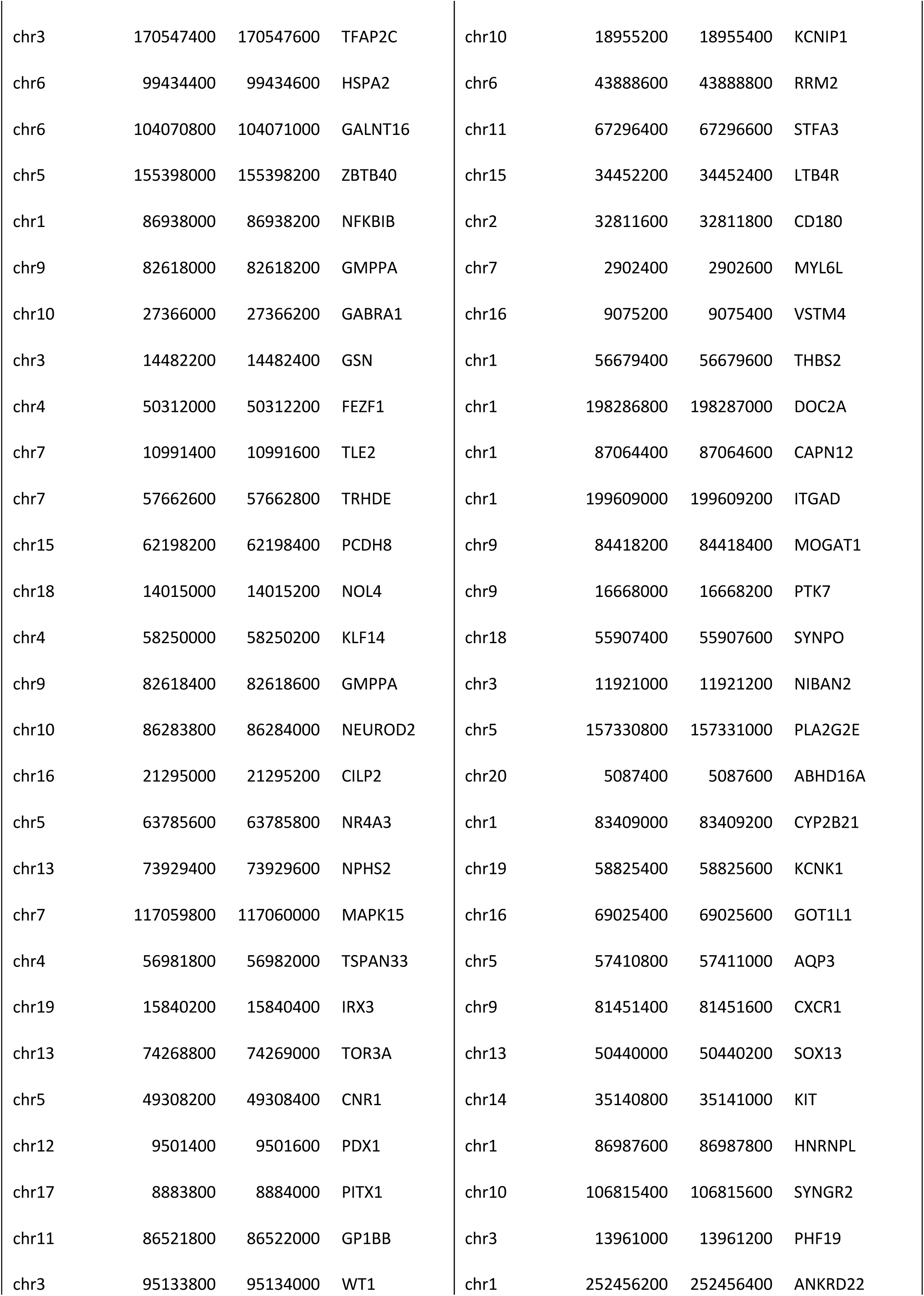
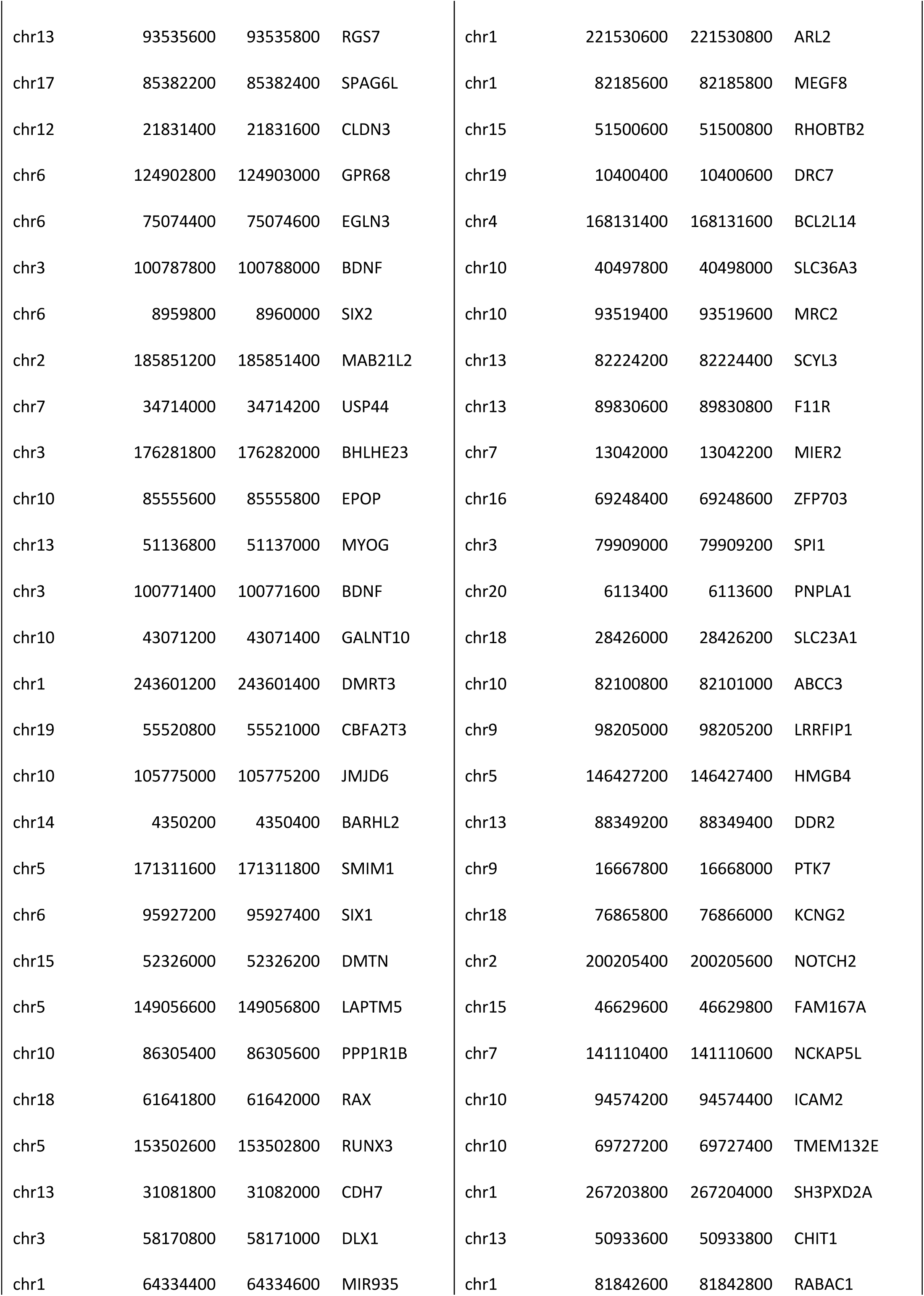
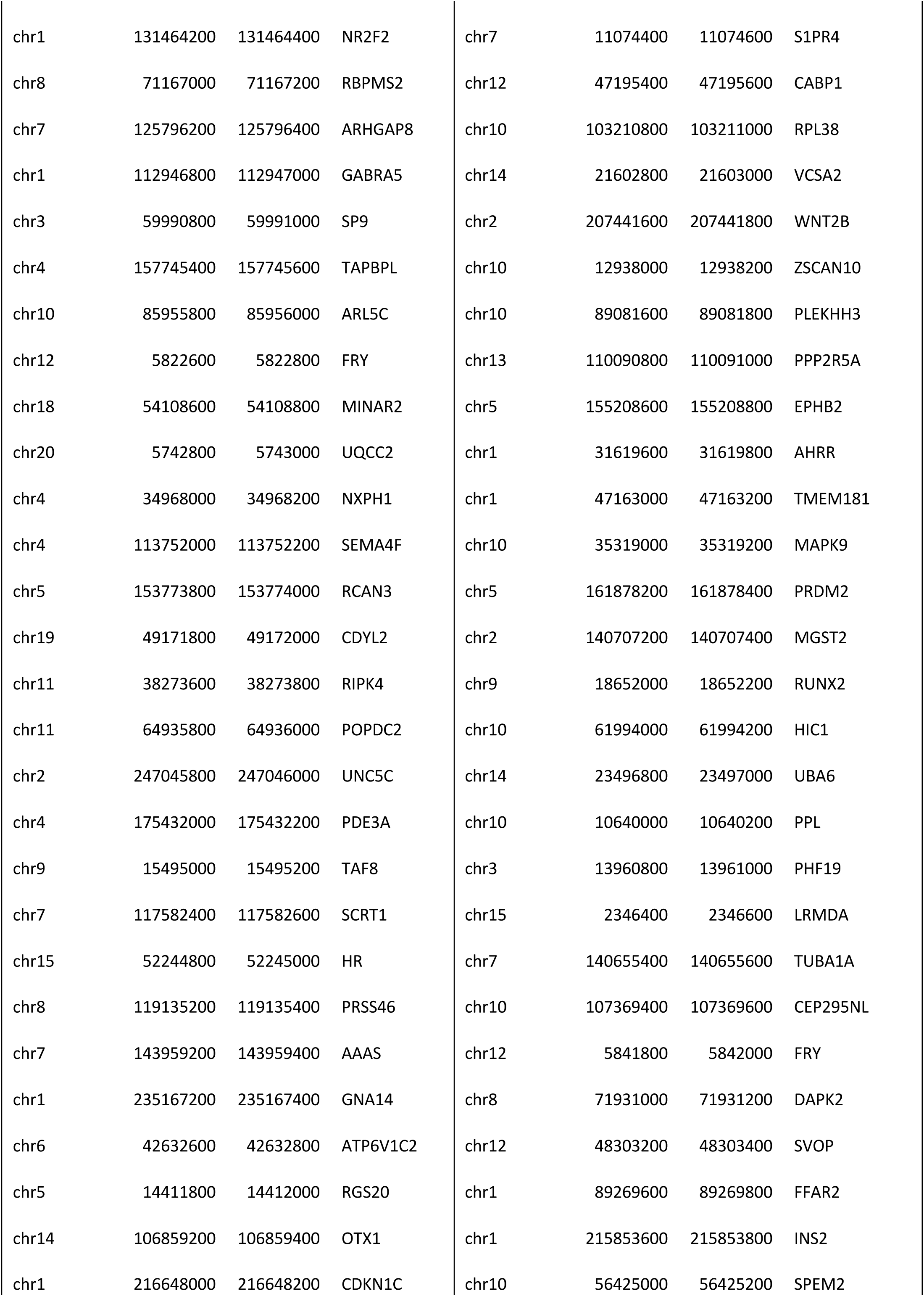
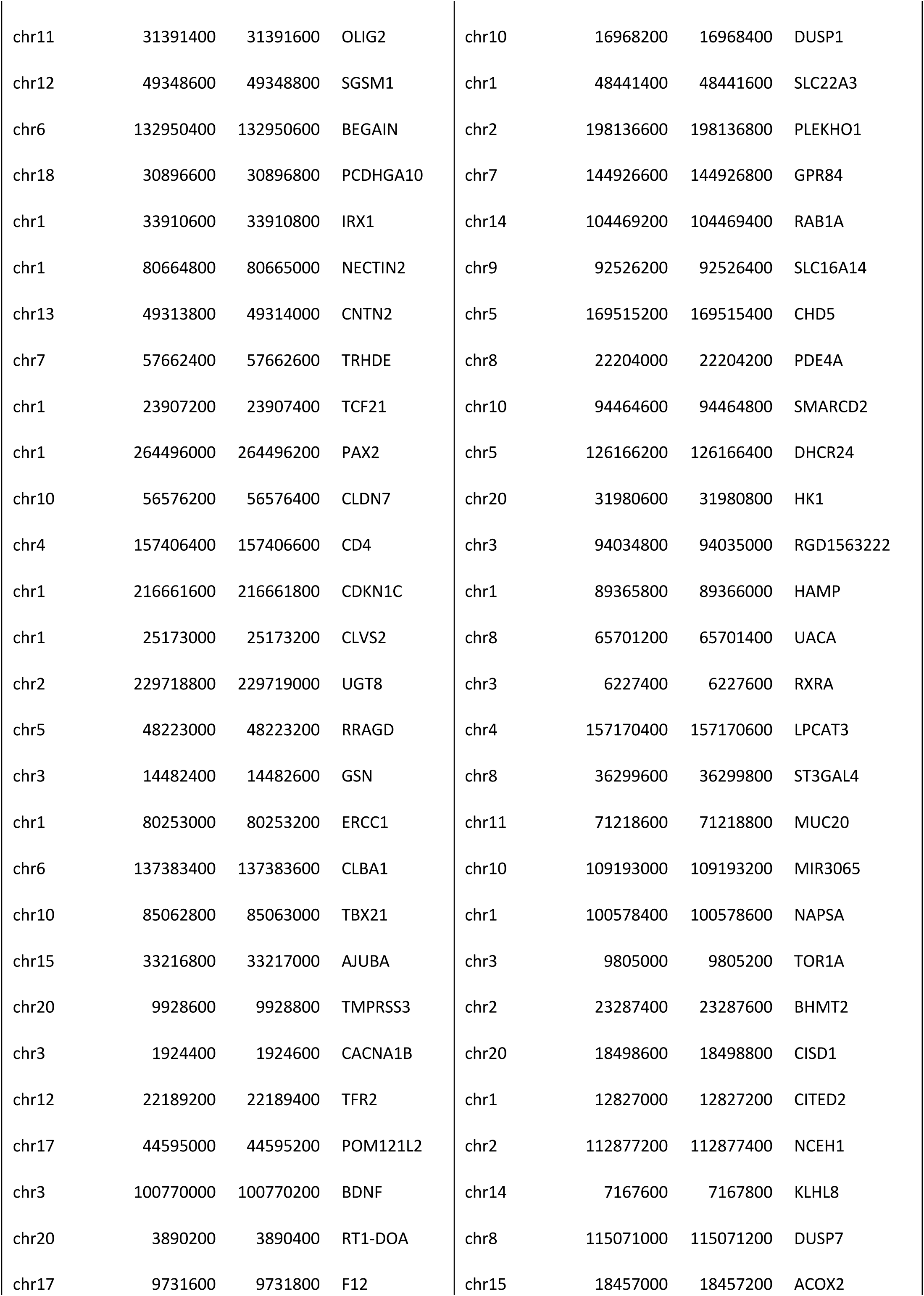
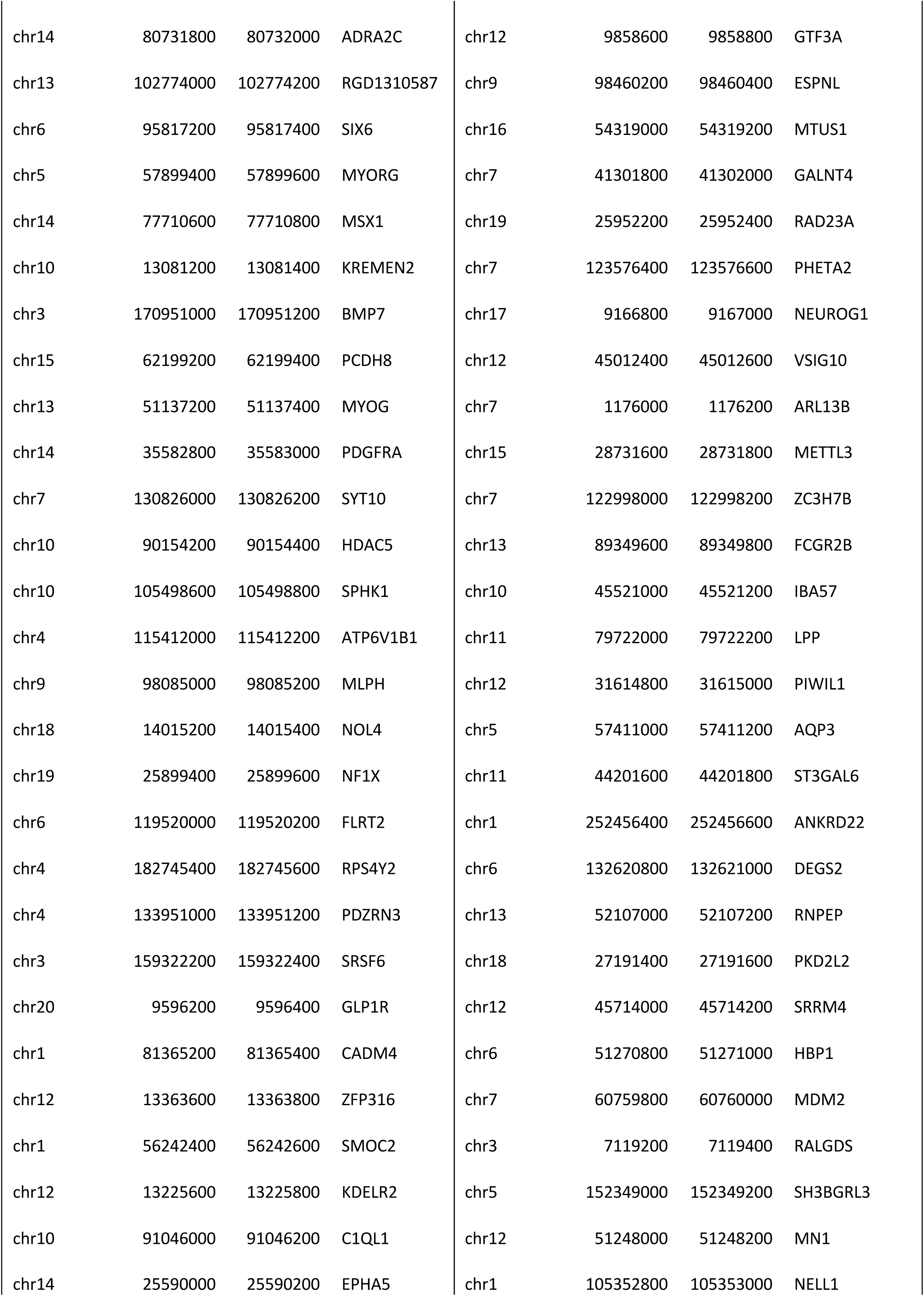
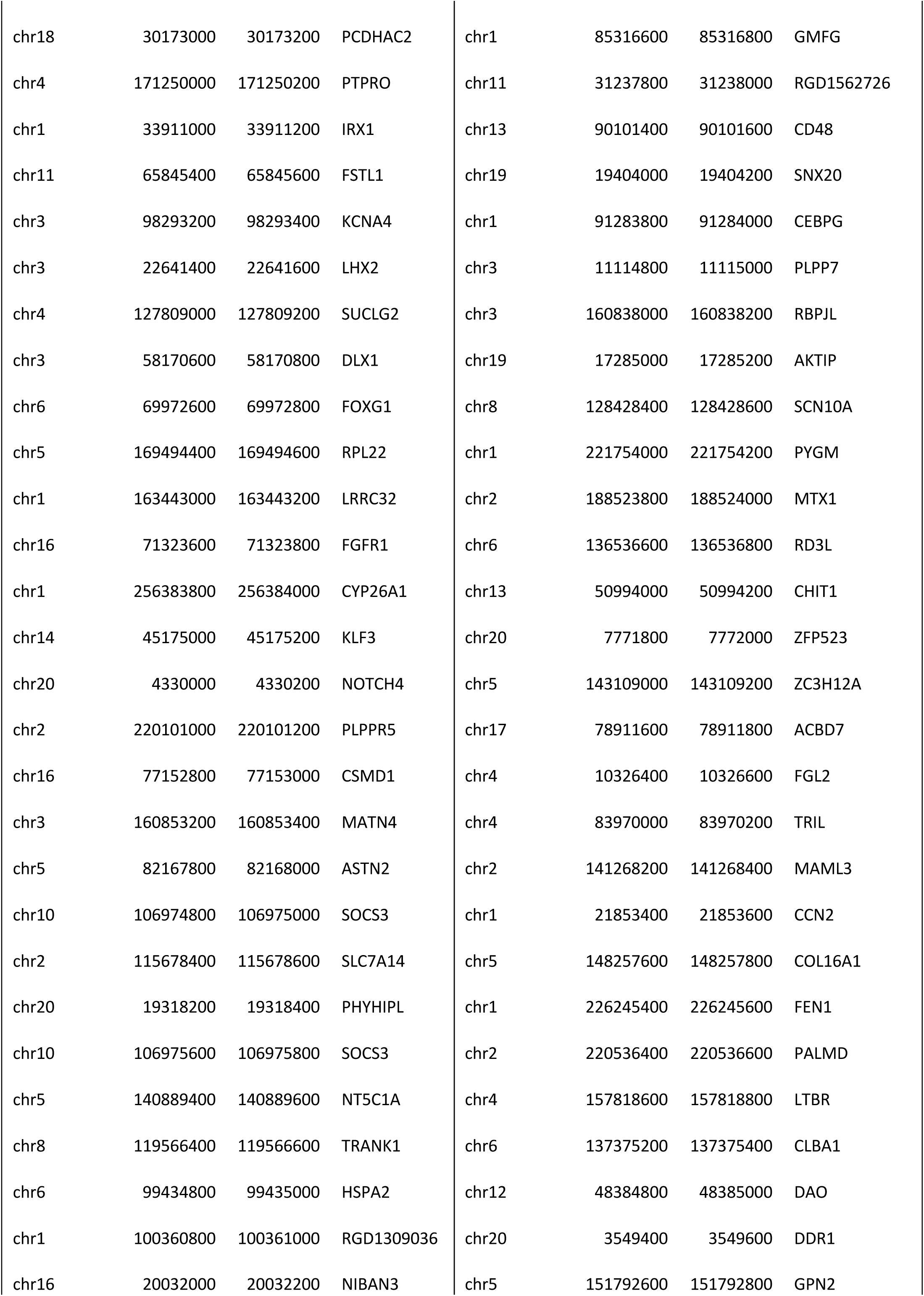
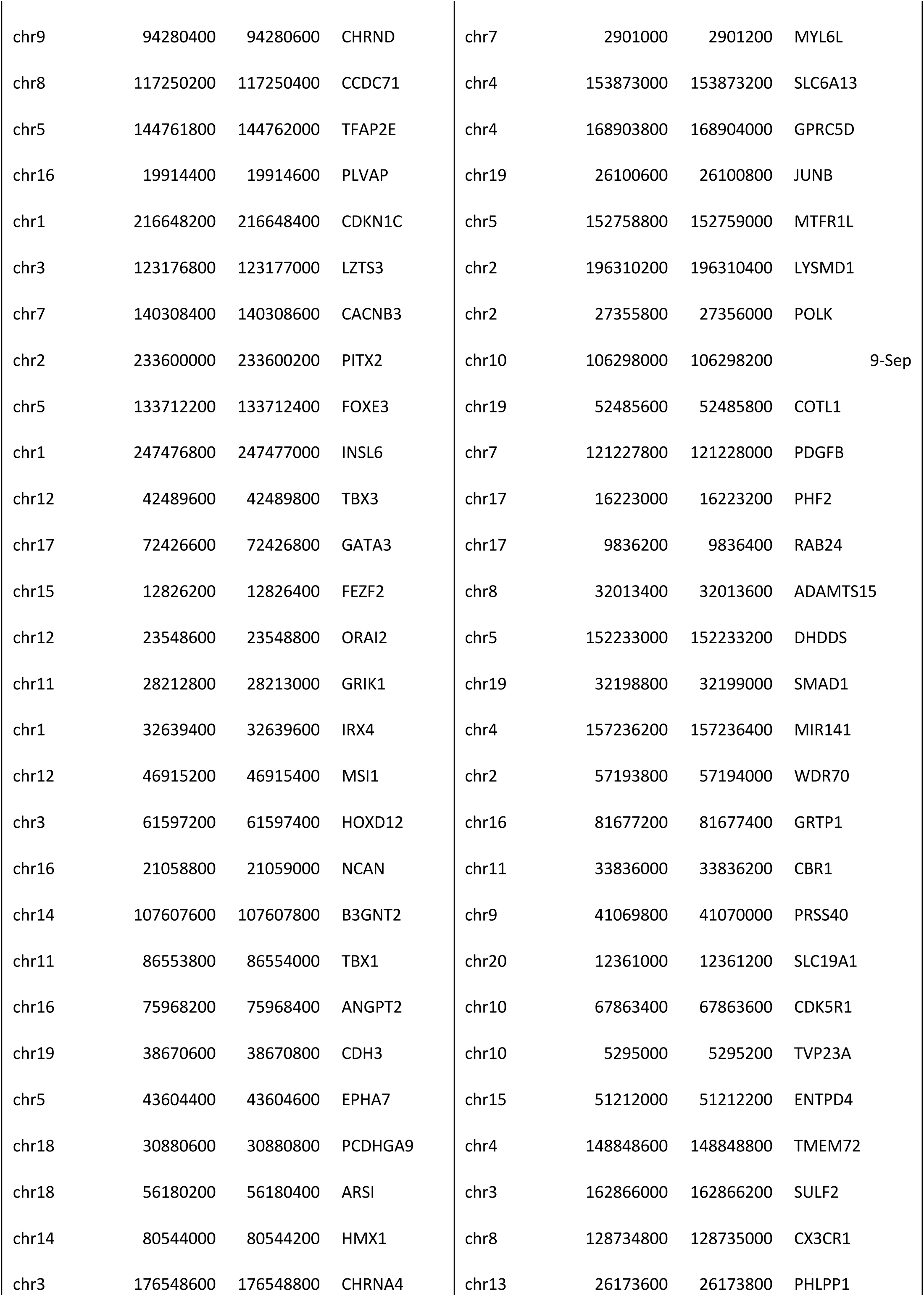
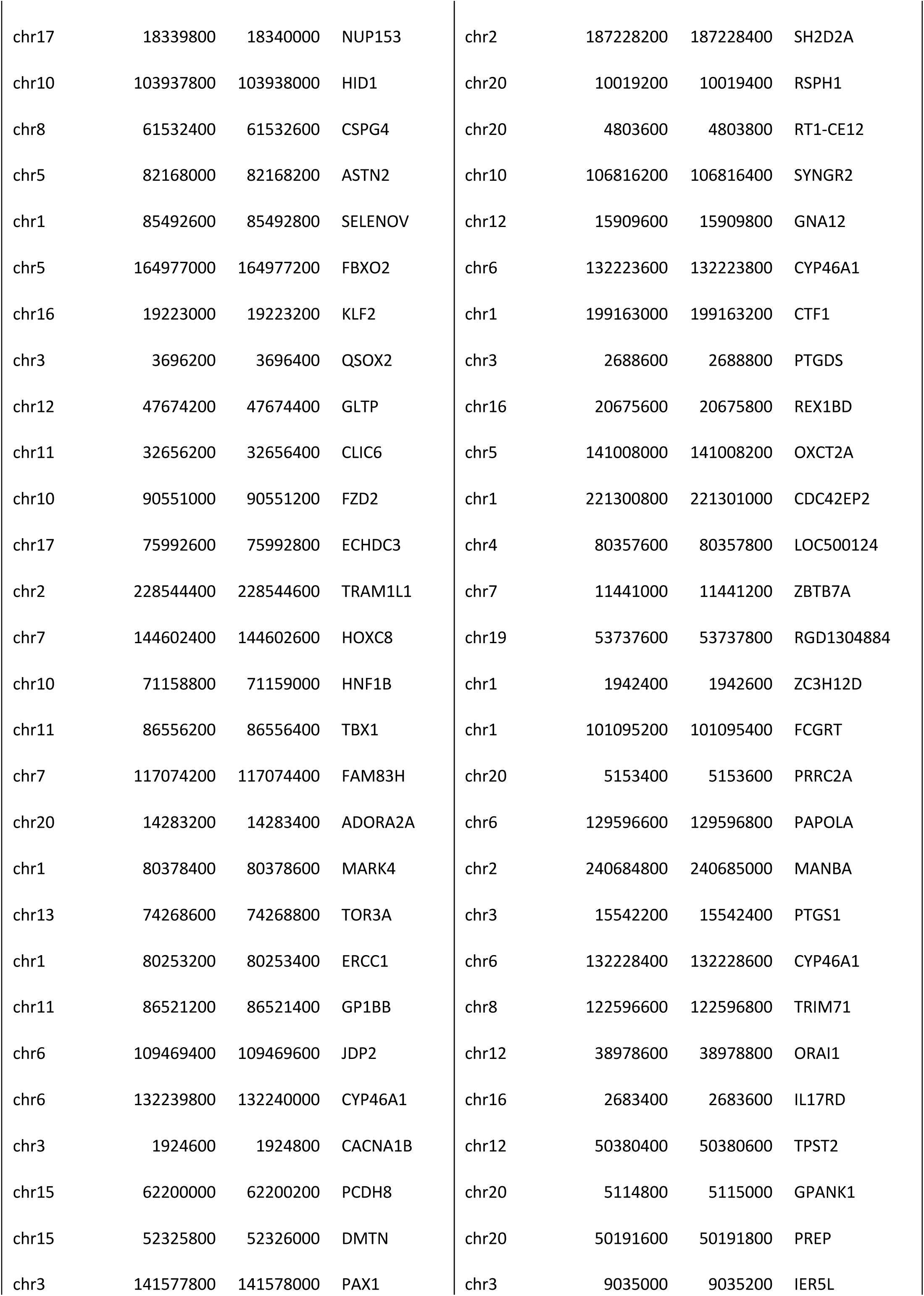
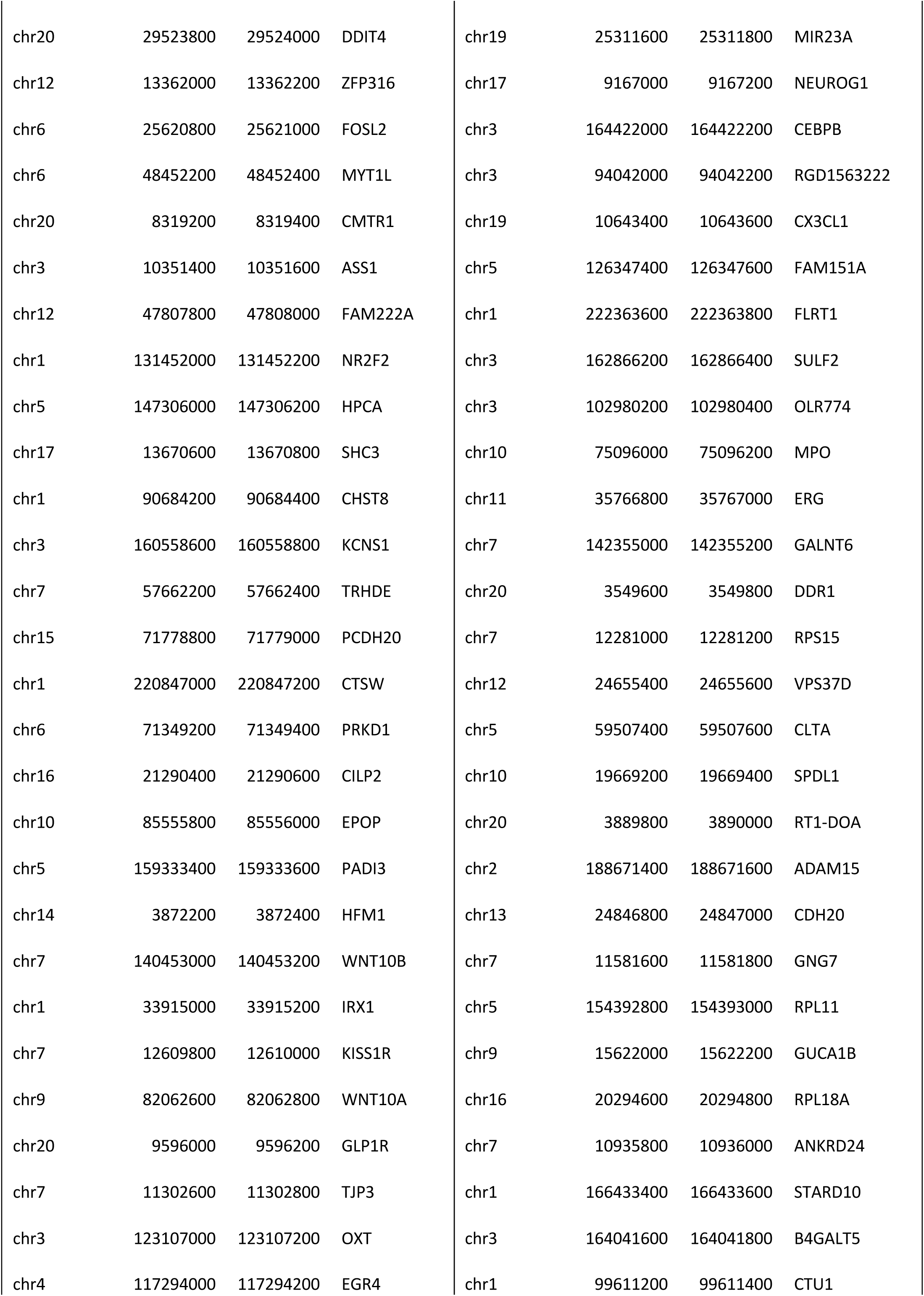
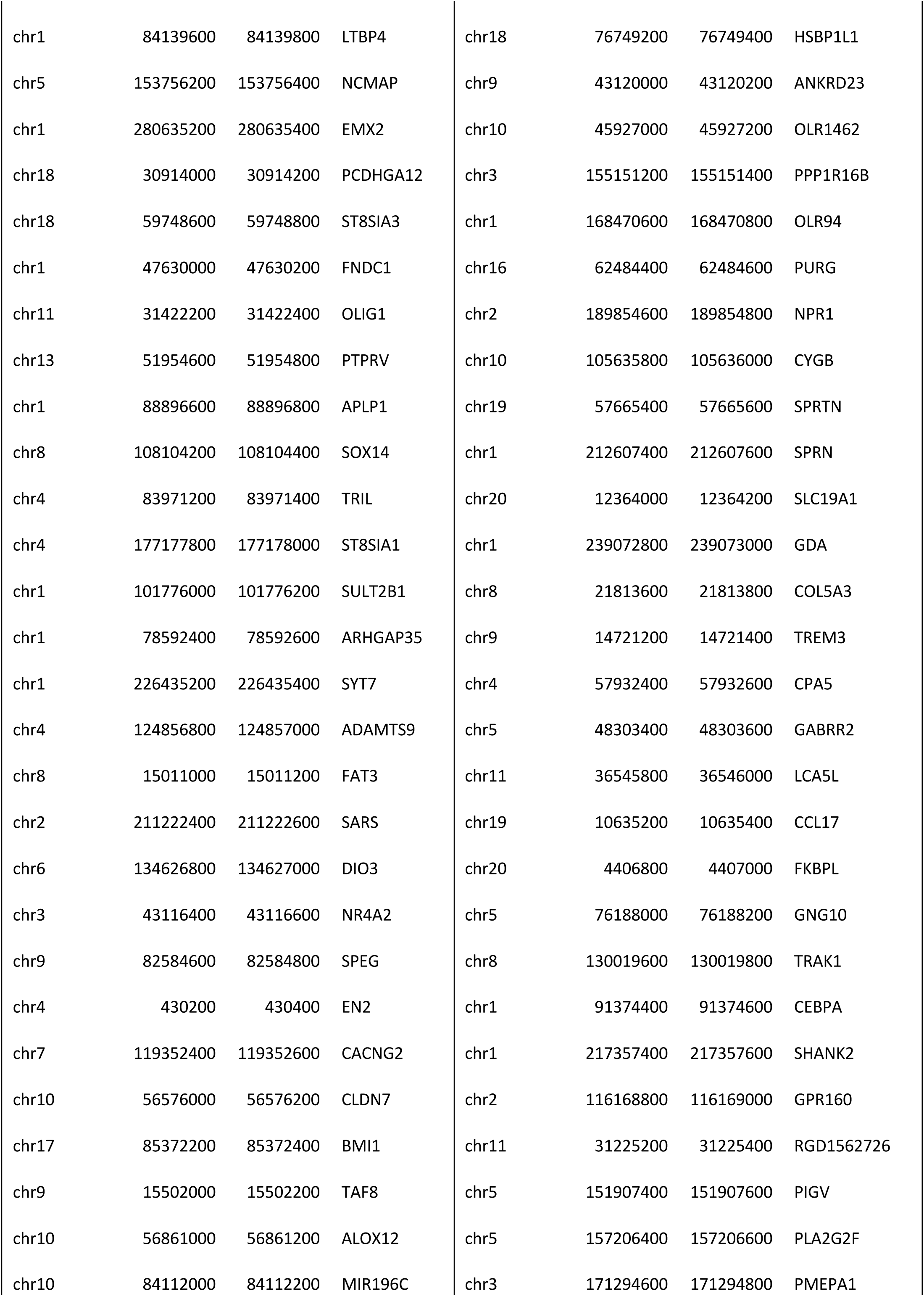
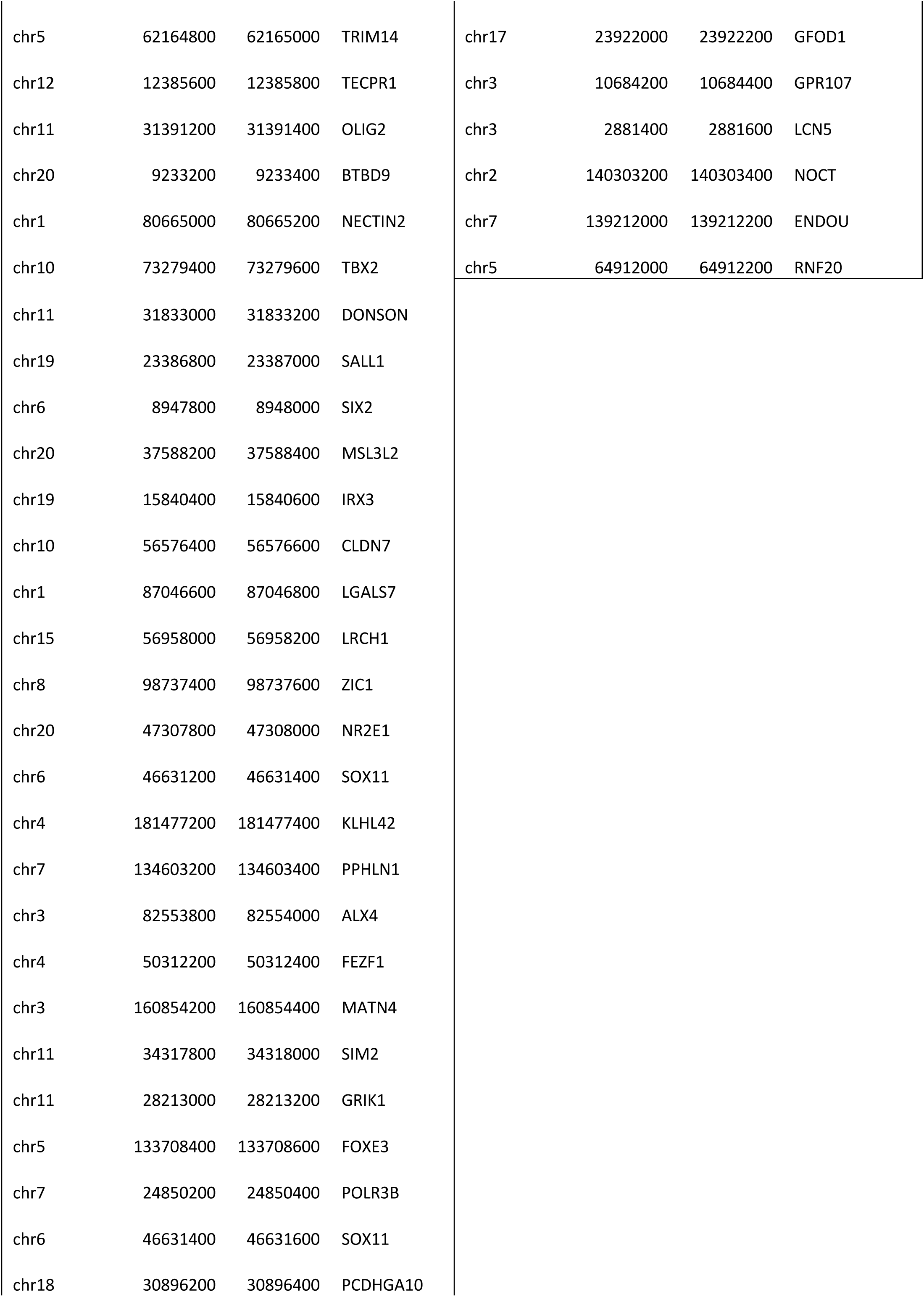
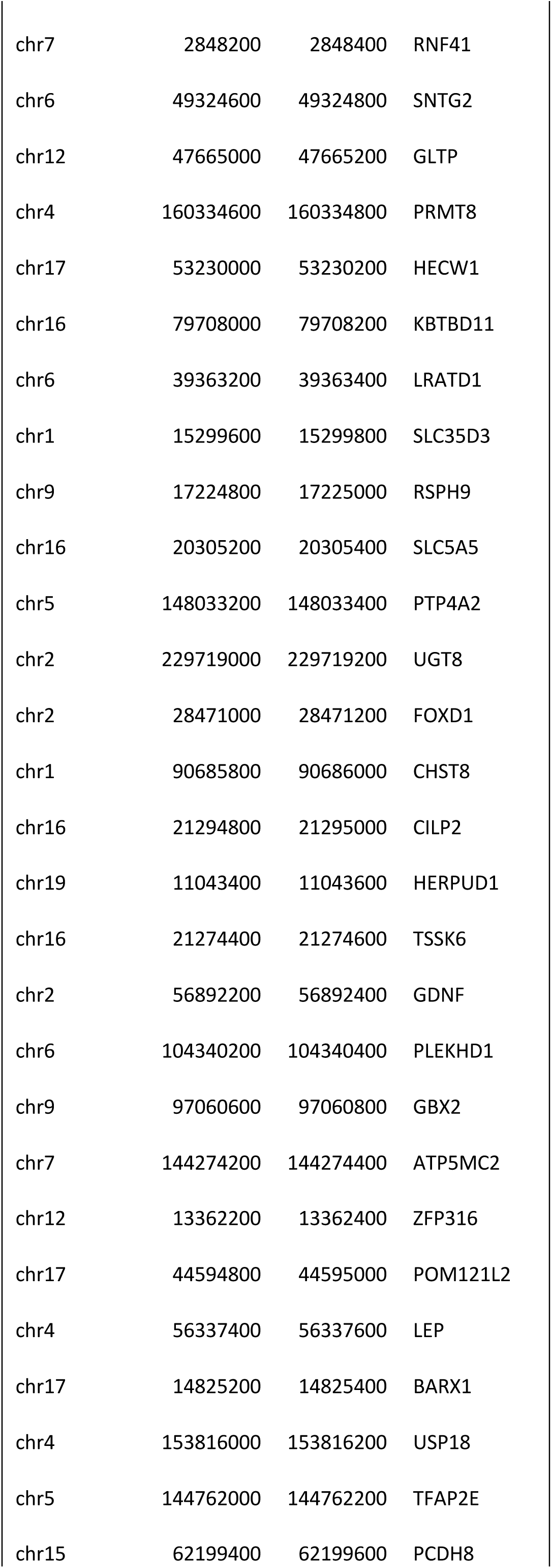
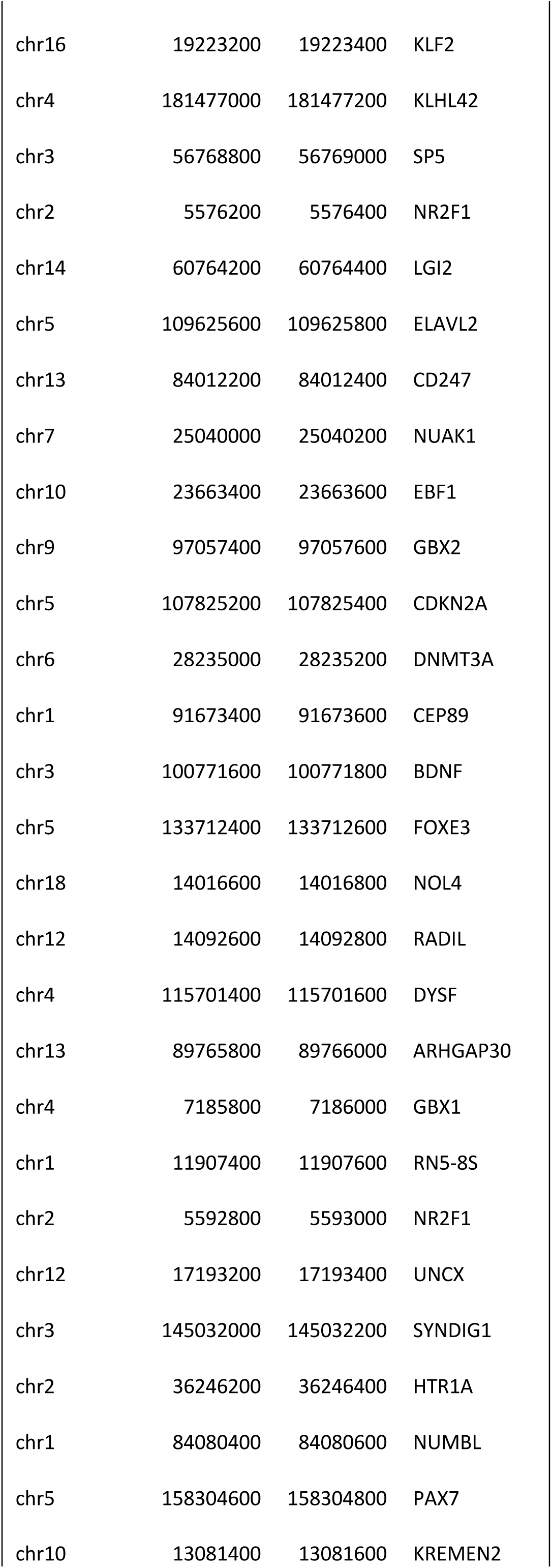
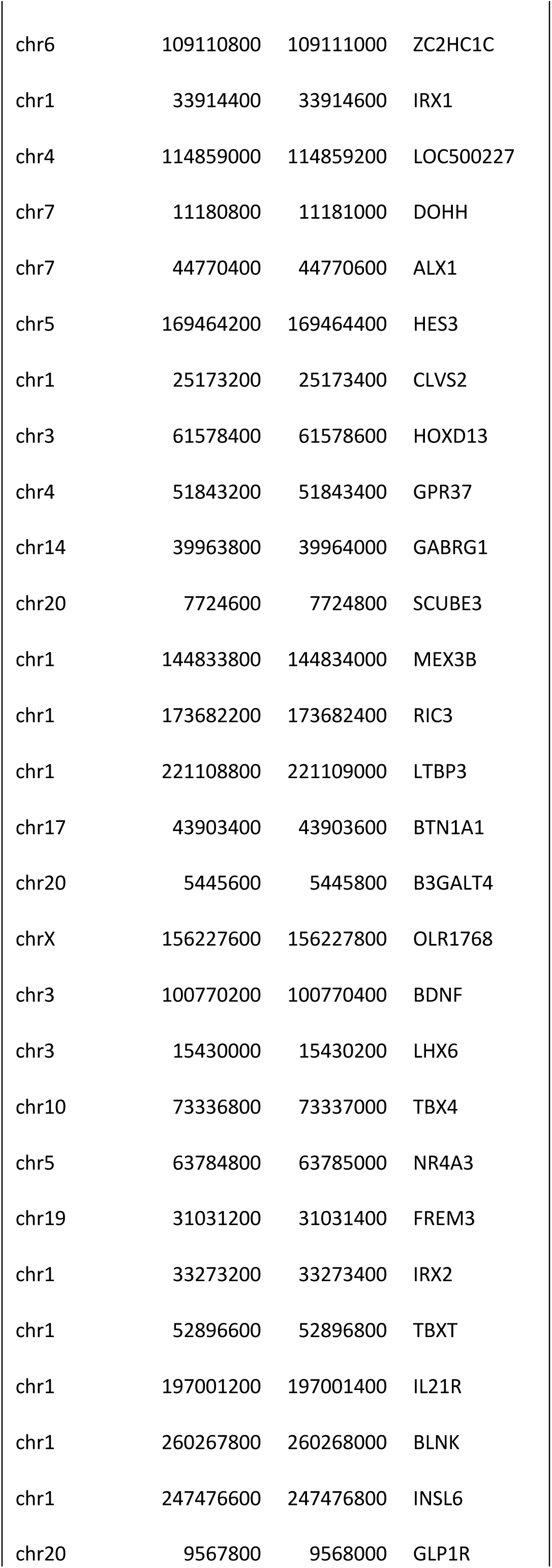
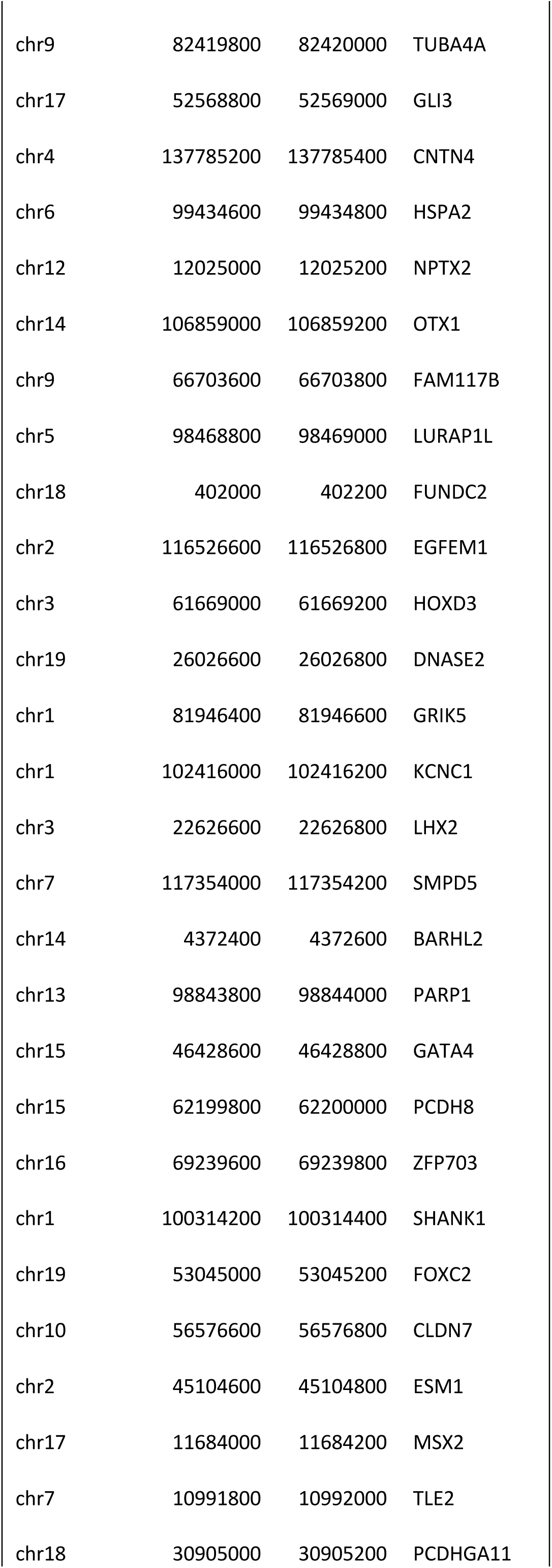
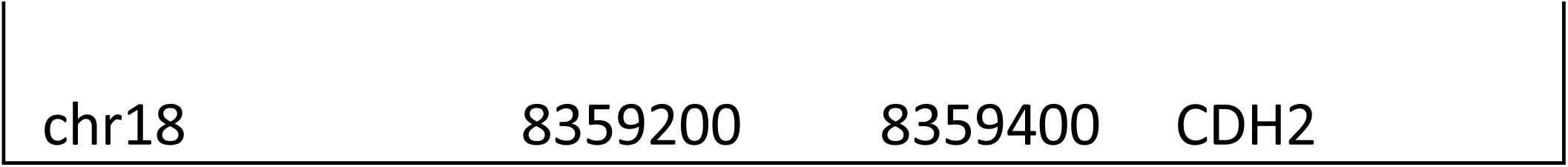
List of differentially methylated regions (DMRs) that are located <20kb from a transcription start site (TSS)

**Suppl. Table 4.**
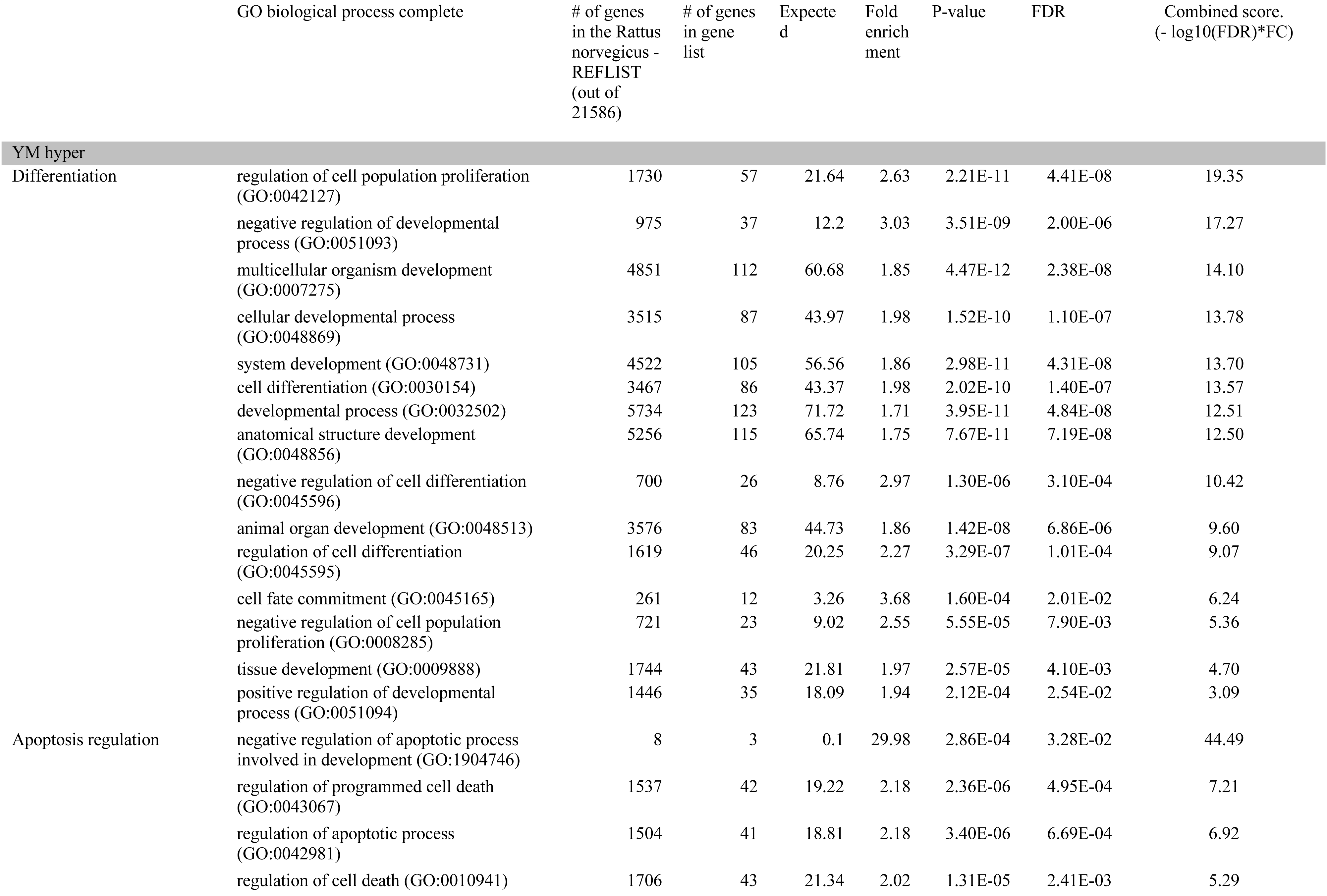
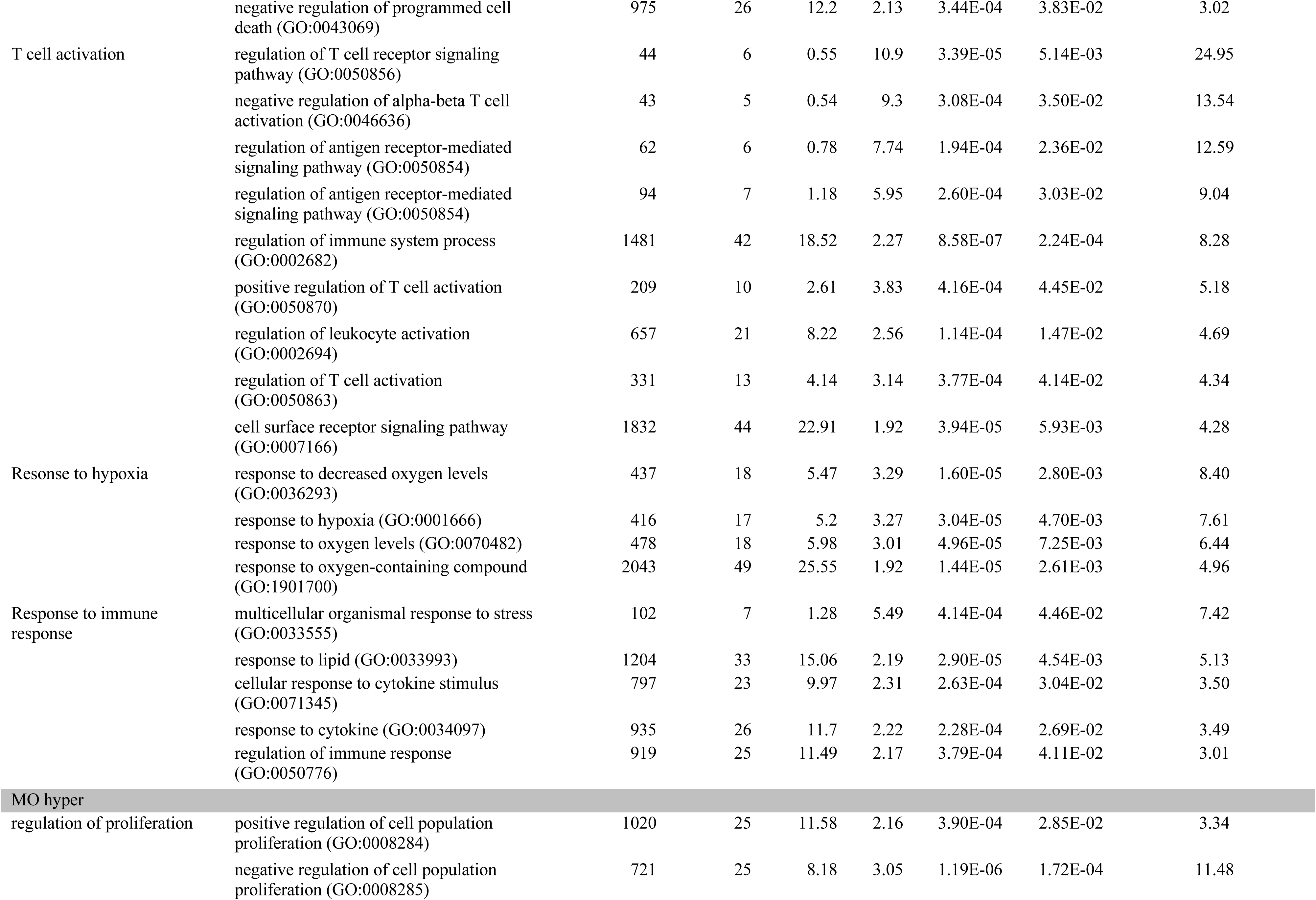
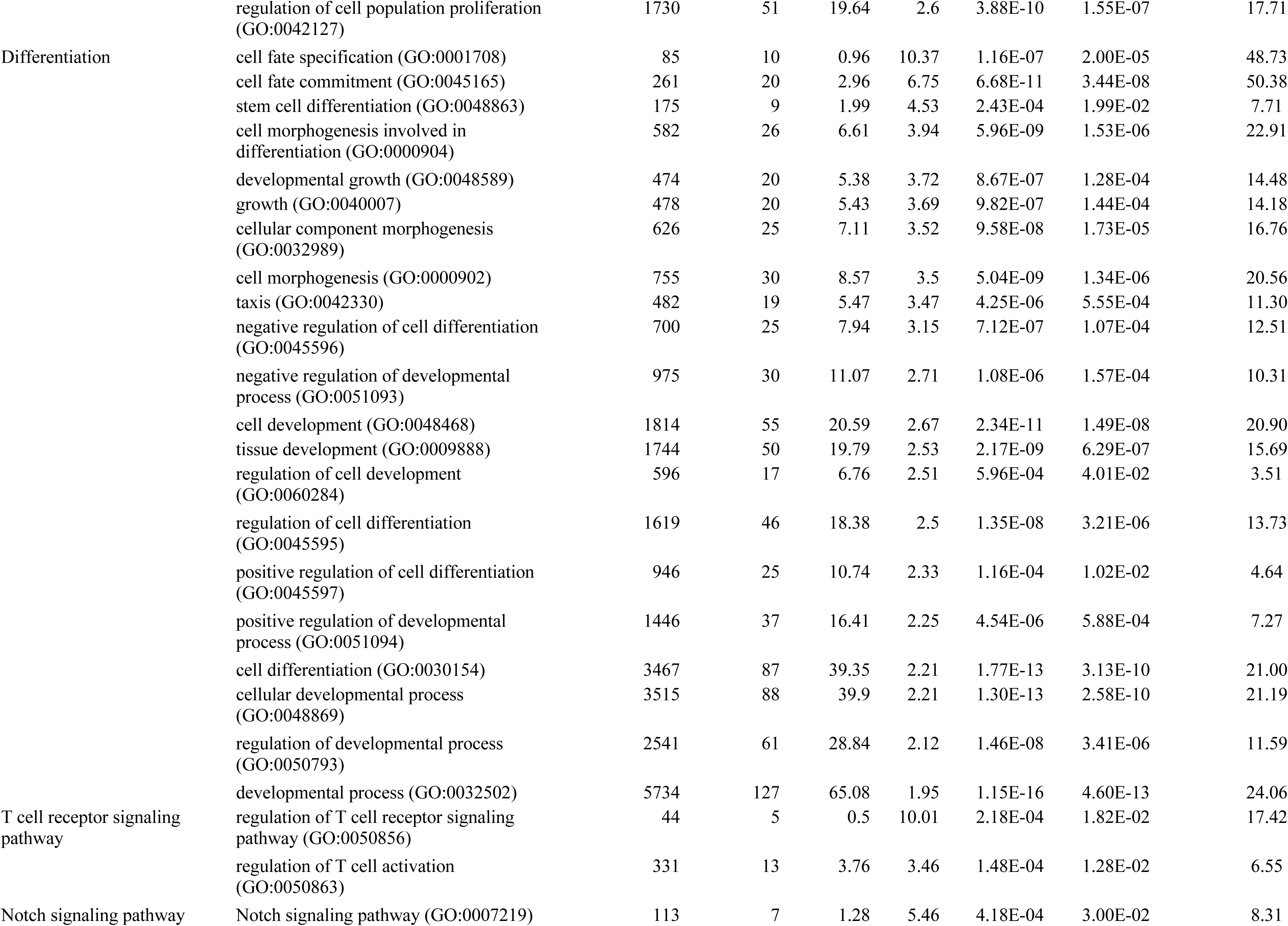
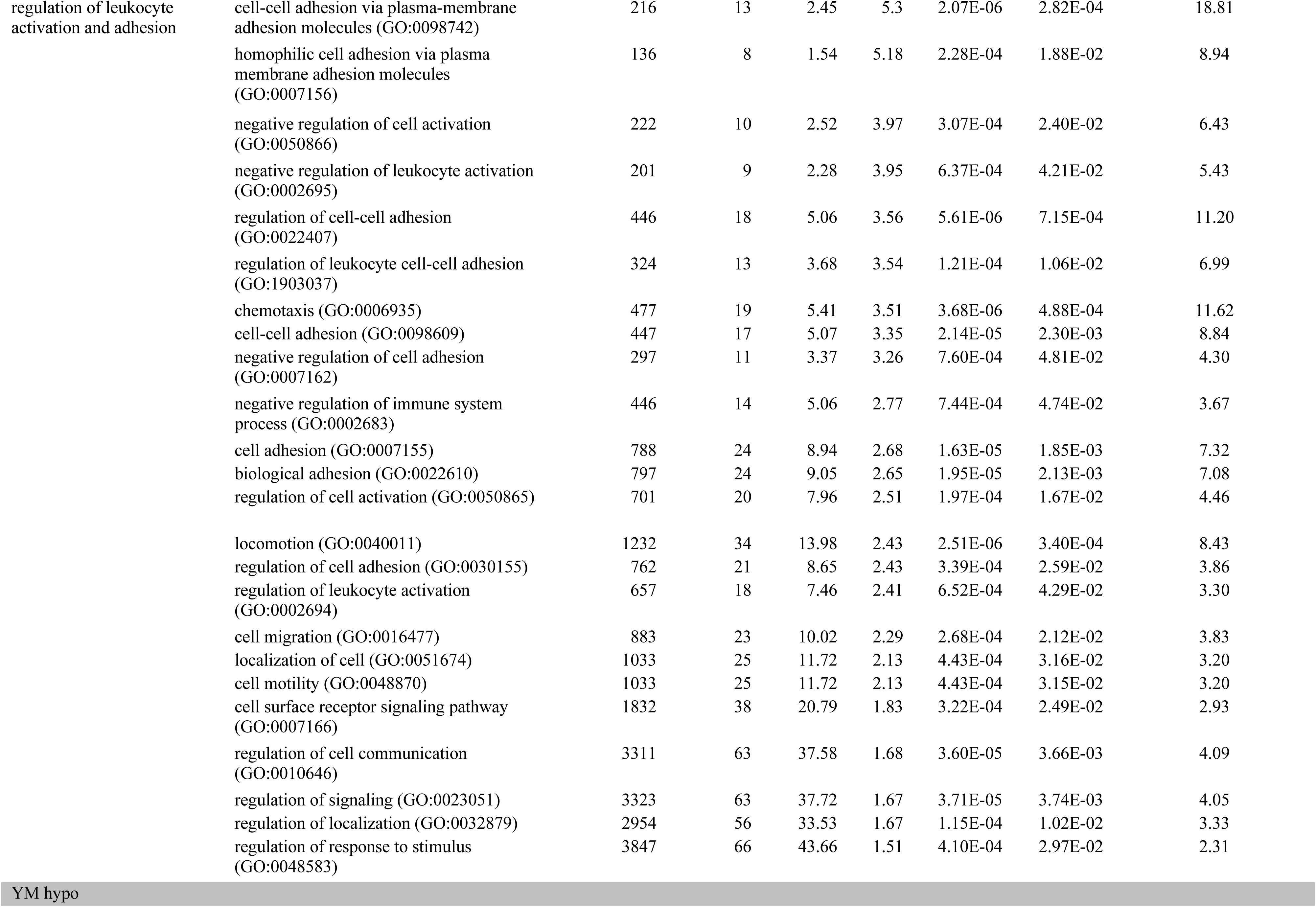
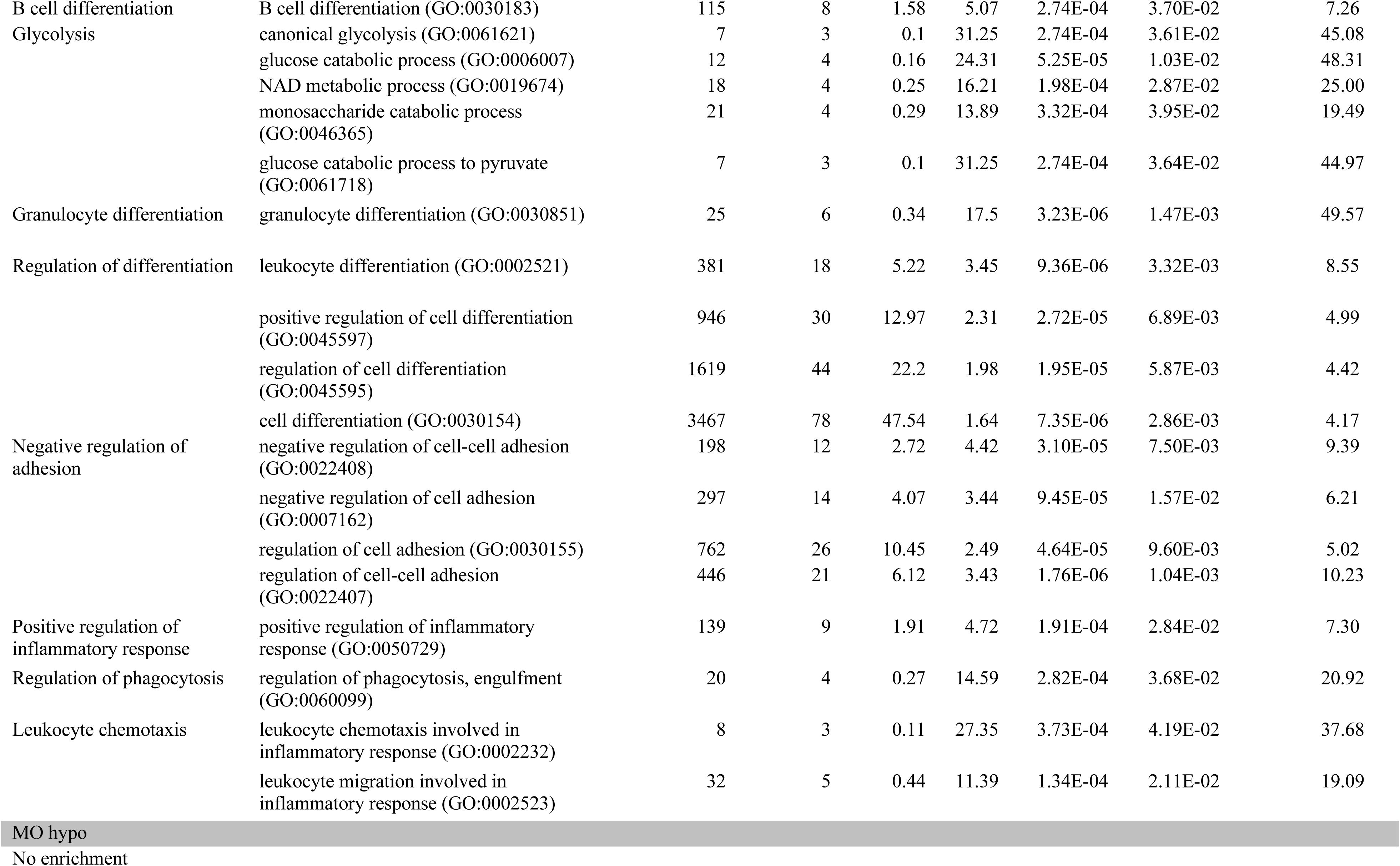
Go enrichment. Analysis was performed using Gene Ontology (http://geneontology.org/docs/go-enrichment-analysis/)

